# (Re)defining the human chromatome: an integrated meta-analysis of localization, function, abundance, physical properties and domain composition of chromatin proteins

**DOI:** 10.1101/2025.10.10.680841

**Authors:** Anna K. Gribkova, Grigoriy A. Armeev, Mikhail P. Kirpichnikov, Alexey K. Shaytan

**Affiliations:** Department of Biology, Lomonosov Moscow State University, Moscow, Russia; Vavilov Institute of General Genetics, Moscow, Russia; Shemyakin–Ovchinnikov Institute of Bioorganic Chemistry, Russian Academy of Sciences, Moscow, Russia; International Laboratory of Bioinformatics, AI and Digital Sciences Institute, Faculty of Computer Science, HSE University, Moscow, Russia

**Keywords:** chromatin, chromatome, epigenomics, proteomics, meta-analysis, genome functioning, protein domains, AI-based protein structure prediction, multivalent protein-protein interactions, intrinsically disordered proteins

## Abstract

The full complement of chromatin-associated proteins—collectively referred to as the *chromatome*—enables genome functioning in eukaryotes by participating in a wide range of physico-chemical processes. These include mediating diverse specific and non-specific intermolecular interactions, catalyzing *in situ* synthesis and modification of macromolecules, facilitating ATP-dependent chromatin remodeling, *etc.* Despite considerable progress in epigenomics and the structural characterization of many nuclear proteins and their complexes, our understanding of chromatin organization at the proteome scale remains incomplete. This gap hinders the development of a holistic view of genome regulation. In this study, we present a state-of-the-art characterization of the human chromatome based on an integrative meta-analysis of diverse data sources describing the composition, abundance, and sub-nuclear localization of chromatin proteins. This effort is complemented by original analyses of their physico-chemical properties, domain architectures, and interaction patterns. To support and streamline these analyses, we developed a reference dataset of chromatin proteins, integrated with an empirical, function-based classification ontology and an associated interactive web resource — **SimChrom** — accessible at https://simchrom.intbio.org/. The reference dataset was carefully curated by reconciling data among protein databases, localization, and mass spectrometry-based experimental studies. Sequence-based and AI-assisted structural analyses revealed previously unannotated domains within chromatin proteins that warrant experimental validation, as well as the widespread use of multivalent interaction strategies that underpin chromatin organization. Together, our findings establish a robust framework for future studies aimed at elucidating genome function through detailed analysis of protein–protein and protein–nucleic acid interactions within chromatin.

**KEY POINTS:**

- The first comprehensive meta-analysis of human chromatin proteins that bridges diverse data types
- Established an interactive SimChrom framework for chromatome research available for the community
- Identified functionally relevant hallmarks of chromatin protein organization

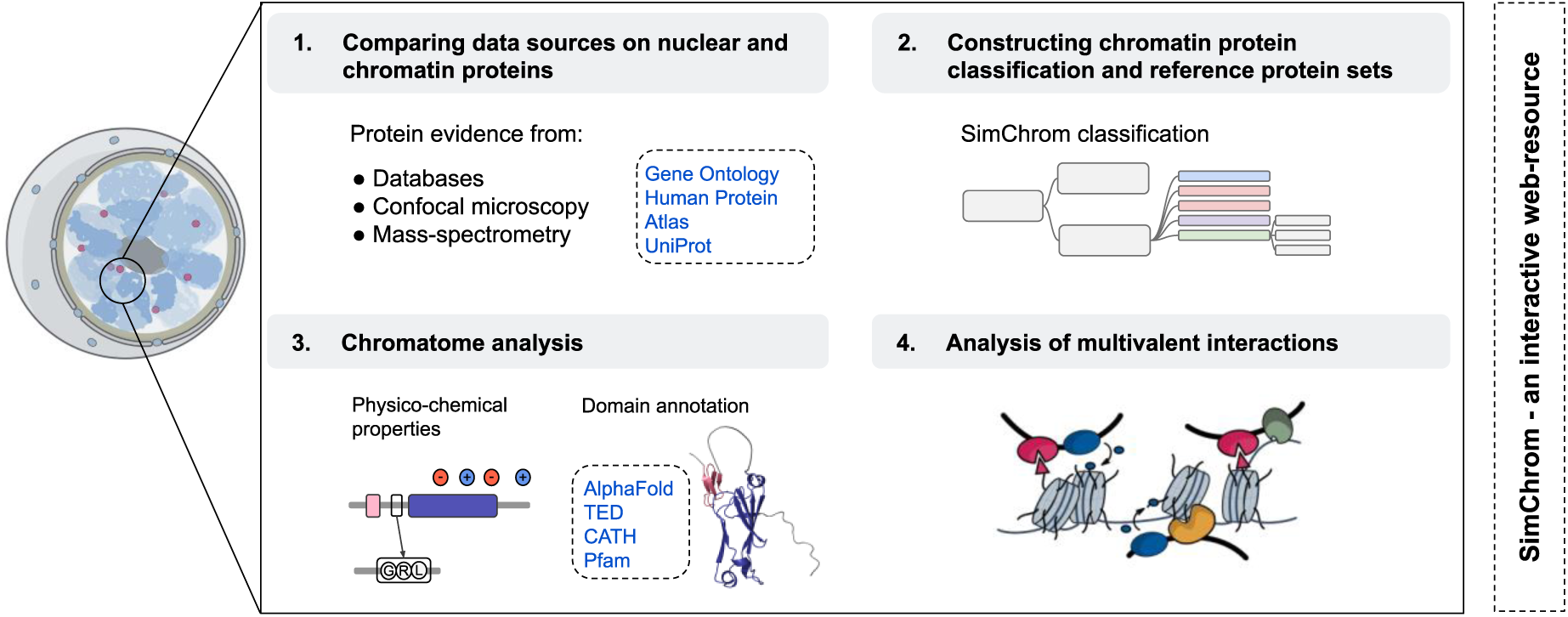
GRAPHICAL ABSTRACT.

## 1. INTRODUCTION

Chromatin, according to the generally accepted definition, is the complex of DNA, proteins, and associated RNA molecules found in the nuclei of eukaryotic cells [1,2] (**Fig. 1A**, **Interactive Fig. 1** at https://simchrom.intbio.org/#nucleus). However, in the crowded nuclear environment, it is challenging to establish stringent criteria that clearly distinguish between macromolecules that form a complex and those that do not, leaving room for interpretation of this definition. Chromatin proteins, collectively called the chromatome [3,4], enable genome functioning in space and time through active ATP-dependent processes and passive protein-DNA/RNA interactions. This functioning employs non-trivial physical phenomena such as liquid-liquid phase separation [5,6], topological constraints on the DNA, DNA looping and loop extrusion [7,8], diffusion in the crowded macromolecular environment [9], multivalent cooperative interactions [10,11], *etc.*, all of which are regulated by the chromatome composition at specific locations and the properties of individual proteins including their post-translational modifications (PTM), domain architecture and intrinsically disordered regions (IDR).

**Figure 1.**
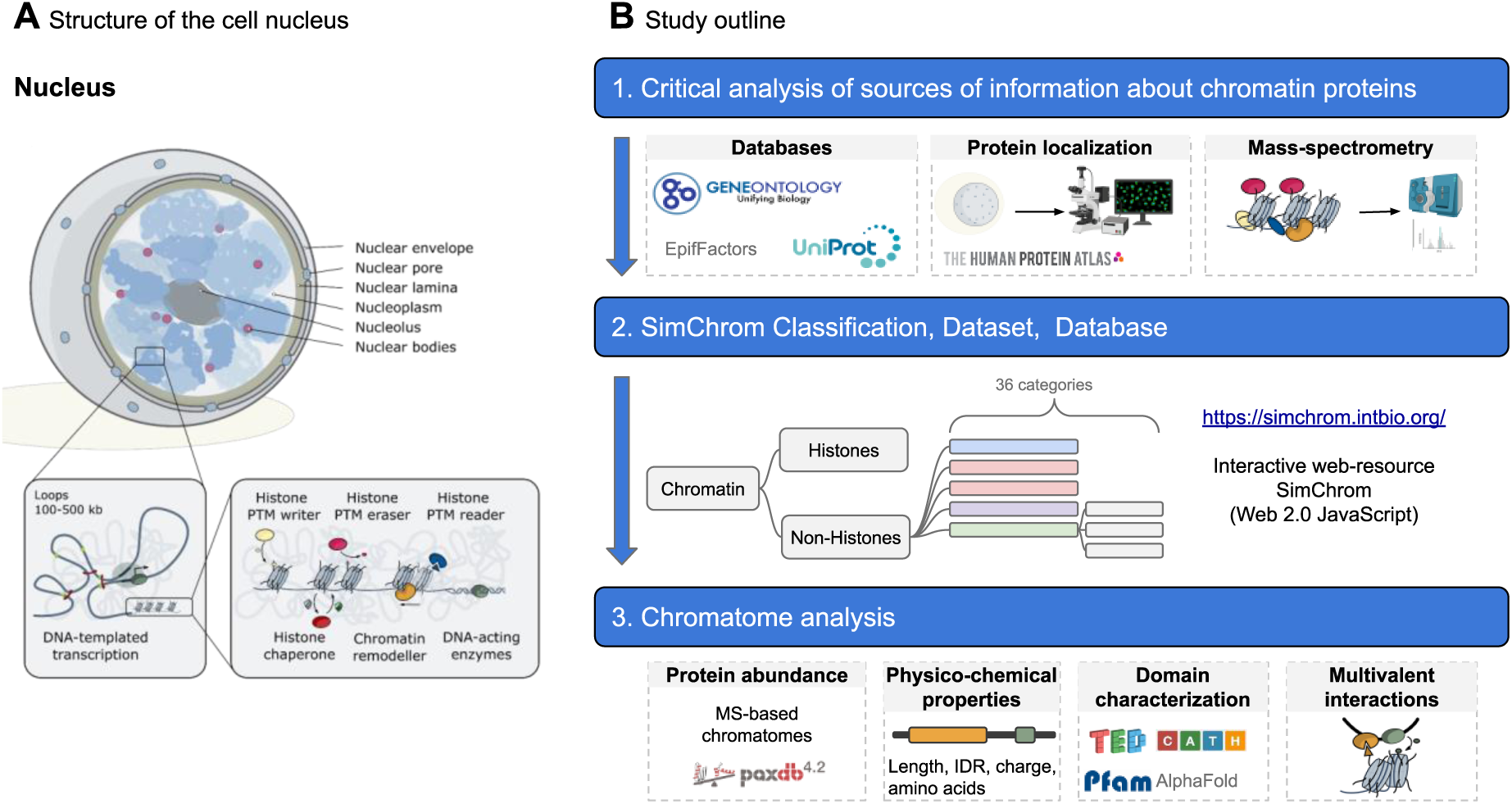
(**A**) The structure of the nucleus with details of chromatin organisation at the levels of chromatin domains and chromatin regulatory proteins. **(B)** The overview of this study shows sources of information about chromatin proteins (*e.g.*, their functions, subcellular localization, identification by MS-based methods or specific protein functional domains) and their use in the current study.

After the discovery of nucleosomes in 1970-ies chromatin research focused on elucidating molecular underpinnings of the genome organization and function at the nucleosome and supranucleosome levels [1]. During recent decades through the contributions of cryo-EM, epigenomics and 3D genomics much more details on the organization of large macromolecular assemblies [12], protein-DNA interactions and DNA topology [13,14] within chromatin have become available. We are at a point when holistic quantitative or at least qualitative models of the genome functioning based on integrating our knowledge about numerous molecular interactions and processes may seem to be within reach [15,16]. The scope of the data required for such models would lie beyond the one provided within the typical frameworks of genomics and epigenomics, and should also rely on what is sometimes referred to as “chromatomics” [3] – the systematic study of the entire content of the eukaryotic nucleus, including chromatin proteins, their spatio-temporal distribution and interactions. However, our understanding of the protein content of chromatin and its functioning at the “omics”-level is currently lagging behind our ability to probe DNA sequence, its epigenetic markup and 3D contacts. It faces certain challenges, which we detail below, with the human chromatome in mind.

The first set of challenges lies in precisely defining the set of proteins that make up the chromatome. Historically, the definition was operational in nature relying on experimental chromatin extraction followed by the analysis of the protein content via physico-chemical methods (see a historical account by K.E. van Holde [1]) and later by various flavours of mass spectrometry analysis combined with different chromatin extraction and treatment techniques (review by van Mierlo and Vermeulen 2021 [17]). Unsurprisingly, the results of such studies depend on several factors – the details of the chromatin extraction techniques (*e.g.*, non-strongly associating proteins may be not extracted), alternatively cytoplasmic proteins may contaminate the sample [18,19], the sensitivity and the resolution of the analysis method (*e.g.*, low abundant proteins may be not detected, variations in post-translational modifications, alternative splice isoforms) [20–22], and the transient nature of the expression of some nuclear proteins. An additional complication is the heterogeneous and dynamic composition of chromatin (sometimes called a fuzzy organel) – it depends on the cell type, cell cycle phase, as well as on the conditions experienced by the cell [23]. One has to keep in mind also that many proteins shuttle between nucleus and cytoplasm. Recent proteomics studies have estimated the number of chromatin proteins to be around 200 – 3800 [4,23–33]. Despite the above mentioned challenges, for many chromatin analysis tasks having a list of chromatin-associated proteins in the starting point.

Since the human chromatome contains at least several thousand entries, any attempt at the rational understanding and description of its functioning requires some dimensionality reduction approaches. Hence, certain grouping or classification of chromatin proteins that considers their functional properties is desirable. Yet, obtaining such a classification is currently challenging. There are certain historically established classes of chromatin proteins that can be clearly defined (*e.g.*, histones, high mobility group proteins, *etc*.) [1], however, others have become obsolete (*e.g.*, nuclear matrix proteins [34] or cannot be easily defined (*e.g.*, nucleosol proteins). GeneOntology (GO) currently provides the most comprehensive set of annotations related to different aspects of gene products and is routinely used to interpret large-scale biological data, such as transcriptomics and proteomics results [35,36]. However, it cannot *per se* provide a straightforward and easy to comprehend classification of chromatin proteins due to the presence of a large number of chromatin related GO terms connected into a complex cumbersome hierarchy, which may be incomplete in some cases (*e.g.*, lacks a term for histones) or include obsolete terms in certain cases (*e.g.*, nuclear matrix). While ultimate functional classification of chromatin proteins may likely not be possible (due to the complexity of the genome functioning, different proteins contributing to many different functional processes, *etc.*), some approximation is at least needed to establish a framework for a rational reductionist-wise understanding of chromatin by us humans.

The third set of challenges, in our mind, relates to the need for developing systems biology approaches to describe and study chromatin in a holistic way as a complex functioning system [37,38]. Considerable advances in methodology are currently needed to move from studying the structure of individual macromolecular complexes and analyzing sequence-level (albeit genome-wide) epigenomic data towards the quantitative models of chromatin operation that can grasp the emergence of complex organismal functions. Chromatin functioning relies on complex dynamics networks of multivalent interactions between macromolecules. These interactions depend on the abundance of chromatin proteins in a given compartment, their physico-chemical properties, and domain architectures that mediate specific or non-specific interactions. Understanding these issues at the chromatome-wide scale is a prerequisite for building holistic functional chromatin models.

Motivated by the above mentioned challenges and the overall need to build complex models of chromatin functioning, in this work we attempted to provide the state-of-the-art meta-analysis of what is known about chromatin proteins, their localization, abundance, and properties. The uniqueness of this study is in cross-comparison of different data sources including database information, mass spectrometry data, and protein localization data. Our analysis was challenged by a common problem – the limited congruence between different data sources. To address it we developed several reference datasets for chromatin and nuclear proteins based on cross-comparison of different datasets and manual curation. Next we developed a relatively simple empirical function-based hierarchical classification of chromatin proteins (SimChrom classification) which was instrumental for all downstream analyses by allowing to compare properties between different groups of chromatin proteins. Using this framework we: 1) systematically analyzed the abundance of chromatin proteins, identified potential pitfalls in MS-based datasets, and using whole cell proteomics data quantified the presence of different chromatin proteins and chromatin protein groups in the cell; 2) characterized the interplay between amino acid composition of chromatin proteins, the prevalence of intrinsically disordered regions and specific distribution of charged amino acids in their sequences; 3) analyzed the current state of structural characterization and domain annotation of chromatin proteins and based on novel AI-enabled protein structure prediction tools identified more than 200 domains in chromatin proteins that belong to currently unknown structural superfamilies and await experimental characterization, 4) characterized typical patterns of multivalent interactions employed by chromatin regulator proteins mainly engaging combinations of histone methylation, acetylation and DNA binding modes.

Finally, we supplement our analyses with an interactive web resource — **SimChrom** — accessible at https://simchrom.intbio.org/. SimChrom harmonizes data from different sources and may be used for exploration of different chromatin protein groups and properties of individual proteins.

The overall scheme of our work is outlined in **Fig. 1B**. Although our analysis in fact required many iterations to achieve self-consistency (*e.g.*, development of SimChrom classification was performed concomitantly with the development of reference chromatin protein sets, and classification was employed in analysis of the quality and content of different data sources) below we present the logic of our analysis as a series of consecutive steps to the extent possible, offloading detailed consideration of certain aspects that may require the familiarization with the whole manuscript to the **<u>Supplementary Results and Discussion</u>** (Suppl. R&D) sections.

## 2. MATERIAL AND METHODS

For the purpose of all analyses in this study we used the set of human protein identifiers and corresponding amino acid sequences representing the canonical protein isoforms (usually corresponding to the major splice isoforms) as provided by the UniProtKB/Swiss-Prot database (also known as the reviewed section of the UniProt Knowledgebase) (UniProt proteome ID UP000005640, release 2022_2) [39]. The set contained 20,272 gene entries corresponding to 20,225 unique protein IDs (some genes code for identical protein sequences). Wherever needed the original datasets were mapped to the above described set of protein UniProt IDs.

### 2.1. Collection and processing of data about chromatin and nuclear protein repertoires, as well as other protein groups, from databases and MS-based studies

#### 2.1.1. Protein localization data sources (UniProt, HPA, OpenCell)

Protein subcellular localization annotations were obtained from UniProtKB [39] (release 2022_2, subcellular location section), Human Protein Atlas (HPA) [40] (version 22, https://www.proteinatlas.org/download/subcellular_location.tsv.zip, accessed on 09.06.2022), the OpenCell dataset [41] was retrieved from the website (https://opencell.czbiohub.org/, accessed on 10.06.2022). To ensure high-confidence subcellular localization annotations, localization terms were filtered according to database-specific reliability criteria. In UniProt, only annotations supported by at least one evidence tag were retained. In HPA, only annotations with a reliability score exceeding the ‘uncertain’ threshold were included. In OpenCell, only annotations scoring above the lowest quality grade were retained.

For the analysis of protein multiple localization, localization annotations from HPA and UniProt were grouped into the following generalized categories: Nucleus, Cytoplasm, Endomembrane system, Other (including chromosome, secretory, and extracellular proteins in UniProt). Detailed grouping information is provided in **Suppl. Table ST3**.

To estimate overrepresentation or underrepresentation of a particular localization for a list of proteins enrichment analysis based on hypergeometric test (also known as Fisher’s exact test) was used with significance threshold p-value < 0.05. To estimate the congruence of subnuclear localization annotations between UniProt (protein set denoted by A) and HPA (protein set denoted by B) following metrics were used: TP = |A ∩ B|; FP = |B ∩ ¬A|; FN = |A ∩ ¬B|; Union = |A ∪ B|; Jaccard similarity coefficient = TP / Union. Performance measures: Precision = TP / (TP + FP); Recall = TP / (TP + FN); and F1-score = 2 × Precision × Recall / (Precision + Recall).

#### 2.1.2. GO and function-oriented databases

In **Section 3.1**, for a selected set of Gene Ontology (GO) terms, lists of associated human proteins were retrieved from the QuickGO database [42] (GO annotation set created on 2025-03-06) using its REST API. To capture all relevant proteins, annotations were obtained also for all their descendant terms with the relationships “is_a”, “part_of”, and “occurs_in”. Information on human histone and non-histone epigenetic regulators was obtained from the EpiFactors database [43] (version 2.1, released September 10, 2024). DNA-binding transcription factors (dbTFs) were obtained from https://www.ebi.ac.uk/QuickGO/targetset/dbTF, accessed on 01.09.2024) [44].

#### 2.1.3. MS-based studies

We conducted a comprehensive search on PubMed using keywords “chromatome”, “chromatin proteins”, “experimentally obtained chromatin proteins”, “nuclear proteins”, and “nuclear proteome” to gather relevant informational resources containing data on nuclear and chromatin proteins.

All protein entries identified in MS-based studies were mapped to the UniProt human reference proteome (release 2022_02), protein isoforms were collapsed to canonical entries. Gene names were used as secondary identifiers to facilitate mapping for records without a direct match. Outdated UniProt entry identifiers were updated to their current counterparts in the used proteome. Below the specific details of obtaining the protein lists from respective MS-studies are provided (see **Table 1**). From Kustatscher et al. (2014) study [23], which provides interphase chromatin probability scores (ICP) for 7,635 human proteins, proteins with ICP > 0.5 were selected. From Alabert et al. (2014) study [26] entries with missing UniProt ID, gene name, or intensity were excluded, unmapped entries were matched by gene name. No filtering by nascent enrichment or chromatin probability was applied. From Ginno et al. (2018) study [25] proteins quantified with at least two unique peptides and consistent signal across all three replicates of at least one cell cycle stage (G1, S, or M) were selected. Although mitotic chromatin was included, the protein composition across stages was nearly identical, and excluding M-phase would not significantly alter the dataset. From Shi et al. (2021) study [27], protein entries were taken from the experimental run that used conditions involving HaeIII digestion and no 1,6-hexanediol treatment (“condition 2” in the study), representing chromatin-associated proteins extracted under native conditions using the Hi-MS protocol. Only proteins with non-zero iBAQ values in all three replicates were retained. From Torrente et al. (2011) study [4] chromatin-associated protein lists obtained by three three extraction methods were combined (total chromatin extraction, high-salt extraction, and micrococcal nuclease digestion). Protein GI accession numbers were converted to UniProt reviewed entries. From Itzhak et al. (2016) study [45] all entries annotated as “mostly nuclear” by the “Global classifier 2” were taken. From Alvarez et al. (2023) study [28] a combined list of proteins obtained by NCC (Nascent Chromatin Capture) in HeLa S3 cells and iPOND (isolation of Proteins On Nascent DNA) in TIG-3 fibroblasts was taken. From the Ugur et al. (2023) study [24] proteins with non-missing raw Log2 intensity values in all three chromatin replicates for human embryonic stem cells were taken.

**Table 1.** A representative list of chromatome and nucleome MS-based experimental studies (full description available in **Suppl. Table ST1**, datasets can be downloaded in **Interactive** Table 1 at https://simchrom.intbio.org/#download).

| Reference | Studied protein fractions | Chromatin purification methods and additional computation filtration | Type of cells / tissue | Proteins identified (by authors) | Number of proteins in our analysis |
| --- | --- | --- | --- | --- | --- |
| Torrente et al., 2011 | Total chromatin, euchromatin, heterochromatin | (1) Total chromatin extraction with hypotonic lysis, Triton X-100 permeabilization, low-speed centrifugation, and EDTA-mediated nuclear lysis. (2) Salt extraction using high salt buffer (420 mM KCl), sonication, centrifugation, followed by dialysis. (3) Micrococcal nuclease (MNase) digestion, including total digestion and partial digestion to separate euchromatin and heterochromatin fractions. | HeLa S3 | 1038 (total chromatin extraction), 1388 (salt), 949 (MNase); 751 (partial MNase); 1912 (all identified chromatin proteins) | 1501 |
| Kustatscher et al., 2014 | Total interphase chromatin | Chromatin Enrichment for Proteomics (**ChEP**): in vivo crosslinking with 1% formaldehyde, followed by differential extraction under denaturing conditions (SDS, urea), RNase A treatment, centrifugation-based chromatin pelleting, and sonication. To assess chromatin association, the study applied Multiclassifier Combinatorial Proteomics (MCCP), which integrates SILAC-based quantitative proteomics from 35 biochemical and biological perturbation experiments. A random forest machine learning algorithm was trained on curated chromatin and non-chromatin reference proteins to assign each detected protein an interphase chromatin probability score (ICP). | HeLa, MCF-7, HepG2, HEK293, U2OS, DT40 | 1980 (chromatin proteins with ICP>0.5); 7635 (total chromatin proteins with ICP values) | 1956 |
| Alabert et al., 2014 | Nascent vs. mature chromatin | Nascent Chromatin Capture (**NCC**): biochemical isolation of newly replicated chromatin using biotin-dUTP incorporation. Cells were pulse-labeled with biotin-dUTP during DNA replication and fixed after either 20 min (nascent chromatin) or 2 h (mature chromatin). Chromatin was crosslinked with 2% formaldehyde, nuclei were isolated, and chromatin was sheared to 2–3 kb by sonication. Biotin-labeled DNA–protein complexes were isolated using streptavidin beads. For proteomic analysis, nascent and mature chromatin were metabolically labeled by SILAC and processed together. | HeLa S3 | 426 (nascent-enriched); 3995 (all identified chromatin proteins) | 3861 |
| Itzhak et al., 2016 | Total nuclear proteome | Cells were metabolically labeled with SILAC and gently lysed under hypo-osmotic conditions to preserve organelle integrity. Post-nuclear supernatants were fractionated by differential centrifugation into five sub-organellar fractions plus cytosolic and nuclear pellets. Protein abundance profiles across SILAC fractions were processed by PCA and classified using a supervised SVM algorithm trained on curated organelle markers. In parallel, total intensities in the nuclear, cytosolic, and organellar fractions (from label-free MS) were used to assign each protein to global classes such as "mostly nuclear", based on relative signal distribution. | HeLa | 1133 (nuclear); 672 (nucleo-cytosolic); 8710 (total) | 1092 |
| Ginno et al., 2018 | Total chromatin: time-resolved (G1, S, M) | Density-based enrichment for mass spectrometry analysis of chromatin (**DEMAC**): formaldehyde-fixed cells were sonicated, subjected to cesium chloride (CsCl) gradient ultracentrifugation to isolate DNA–protein complexes by buoyant density (1.39 g/cm<sup>3</sup>). Chromatin fractions were collected, dialyzed, decrosslinked, digested with DNase I. | Human T98G (glioblastoma) | 3065 (chromatome); 6242 (total proteome) | 3051 |
| Shi et al., 2021 | Promoter-proximal chromatin | **Hi-MS** (Hi-C-based proteomics, adapted from BL-Hi-C): cells crosslinked with 1% formaldehyde; genomic DNA digested with HaeIII (GGCC sites); ligated with biotinylated bridge linkers; nuclei lysed in 0.2% SDS; chromatin sonicated; chromatin-DNA complexes captured on streptavidin beads. Quantified sensitivity to 1,6-hexanediol evaluated via AICAP index (Anti-1,6-Hexanediol Index of Chromatin-Associated Proteins). | K562 | 3228 | 2848 |
| Ugur et al., 2023 | Total chromatin | Chromatin Aggregation Capture (**ChAC**): nuclei fixed with 1% formaldehyde, lysed with SDS and urea, sonicated, and purified by protein aggregation capture (PAC) on magnetic beads. DIA-MS with DIA-NN used for quantification. | Human ESCs (H9) | 2487 | 1730 |
| Alvarez et al., 2023 | Time-resolved (nascent, G2/M, early and late G1) chromatin | Nascent Chromatin Capture (**NCC**) method, which relies on pulse-labeling newly replicated DNA with biotin-dUTP, followed by formaldehyde crosslinking and sonication-based chromatin fragmentation. Biotinylated DNA-protein complexes were affinity-purified using streptavidin magnetic beads. HeLa S3 cells were synchronized and harvested at five post-replication time points (Nasc, Late S, G2/M, early G1, late G1) across six biological replicates. | HeLa S3 | 1454 (present at all time points in all 6 replicates; from total of 5770) | 1478 (2894 total) |
|  |  | Isolation of Proteins On Nascent DNA (iPOND): formaldehyde crosslinking (1%), EdU labeling for 15 minutes, click chemistry with biotin-azide, chromatin fragmentation, streptavidin bead enrichment. | TIG-3 fibroblasts | 2351 (detected in ≥4 of 5 replicates) | 2397 (2894 total) |

#### 2.1.4. Other sources

Housekeeping proteins were defined as proteins detected in all analyzed tissues by RNA-seq and downloaded from HPA (version 23) [46] (in total 8899 proteins). Manually curated protein complexes were downloaded from Complex Portal (accessed on 7 January 2025) [47]. Only complexes exclusively containing chromatin proteins were selected for domain co-occurrence analysis.

### 2.2 Construction of reference chromatin and nuclear protein datasets

#### 2.2.1. The SimChrom ontology, SimChrom protein dataset, and SimChrom/SimChrom-SL classification

The SimChrom chromatin proteins classification ontology was developed simultaneously with the corresponding set of human chromatin proteins that were collected according to the developed classification. To this end, for almost every SimChrom classification term we attributed specific terms from GO classification that were manually selected to represent molecular functions and biological processes that happened exclusively inside the cell nucleus and were related to the respective SimChrom term. In certain cases cellular component GO terms were also used when they were directly related to the respective SimChrom term (*e.g.*, complexes of chromatin remodelers). The list of the GO terms attributed to every SimChrom term is given in **Suppl. Table ST4**. The dataset was then supplemented by proteins defined as chromatin proteins by several databases, review papers, and original studies (*e.g.*, HistoneDB 2.0 – histone proteins, Histome2 – for PTM writers, see details and corresponding data sources in **Suppl. Table ST4**). At the last step for certain SimChrom categories the contents of the dataset were additionally filtered to either remove the potentially non-nuclear proteins (in the case of RNA-binding proteins) or histones from the categories belonging to the “non-histone proteins” group (see details in **Suppl. Table ST4**).

To ensure unambiguous categorization, we constructed a SimChrom-SL (single labeled) classification, where each protein was assigned to exactly one category of the same SimChrom ontology. Assignment followed a predefined priority order: “Molecular function” and “Physico-chemical properties” categories were prioritized, followed by others (“Biological processes” and “Genomic location”). Within these groups, categories containing fewer proteins were ordered first (see priority hierarchy in **Suppl. Fig. SF3_2**). Each protein was labeled with its first eligible category in this sequence.

To assess the quality of the SimChrom dataset, Gene Ontology enrichment analysis was performed. Two groups of proteins were analyzed separately: (1) proteins present in SimChrom but absent from NULOC_CS (see below), and (2) proteins present in NULOC_CS but absent from SimChrom. GO enrichment was assessed using g:Profiler [48], with multiple testing corrections applied (Bonferroni correction, significance threshold 0.05). Only driver GO terms (defined by g:Profiler) were selected for further interpretation.

#### 2.2.2. Construction of reference localization datasets

Reference datasets of nuclear (abbreviated as NULOC), non-nuclear (NON_NULOC) and cytoplasmic (CYTLOC) protein entries at different levels of confidence and uniqueness of localization were constructed (see set of localization terms in **Suppl. Table ST3**). The datasets were created using the combination of localization information from UniProt and HPA (full details are provided in **Suppl. Table ST6**). Datasets of proteins having only one specific localization (*i.e.* no other localization reported in the source databases) are denoted by the UL suffix. Datasets, where localization is simultaneously supported both by UniProt and HPA are denoted by CS (consensus) suffix, alternatively, if localization is supported by at least one database the JT (joint) suffix is given. The NECF (“no evidence code filtration”) suffix denotes datasets where no preliminary filtration of localization information provided by the databases based on evidence codes or confidence levels were applied. We noticed that many histone protein entries in UniProt lack evidence codes for their localization (in release 2022_2). This is apparently due to the fact that these entries appeared in UniProt before the manually curation of evidence attribution was introduced, and they are still awaiting manual review and retrofitting. We manually added histone proteins in all constructed nuclear reference datasets.

### 2.3. Protein abundance data processing

#### 2.3.1. PaxDB data

Two dataset from PaxDB [49] version 4.2 with protein abundance data were used: the dataset with the highest proteome coverage (“*H. sapiens* – Whole organism (Integrated)” – covers 99% of human proteome according to PaxDb, referred to as “PaxDb_INT’’ in this paper) and the dataset with the highest interaction consistency score (“Whole organism, SC (Peptideatlas,aug,2014)” – covers 84% of human proteome according to PaxDb, referred to as “PaxDb_PA’’ in this paper), see also **<u>Suppl.</u> <u>R&D</u> Sec. 3.1**. An abundance unit is protein per million, ppm, which describes protein abundance relative to all expressed molecules in the proteome. Protein abundance was obtained by aggregating abundance of individual genes with similar protein sequences (*e.g.,* in the case of canonical histones). Cumulative abundance was defined as the sum of protein abundances for a group of protein entries; cumulative weight was calculated as abundance multiplied by the protein molecular weight.

#### 2.3.2. Abundance from MS-based studies

Protein abundance values were derived from mass spectrometry (MS) intensities or iBAQ values reported in individual studies. When a single intensity value was associated with multiple protein identifiers, it was divided equally among them. For proteins represented by multiple entries within a study, the median intensity (or abundance) value was used to obtain a single estimate per protein. For Kustatscher et al. (2014) study protein abundance was defined as the summed MS intensity. For Alabert et al. (2014) study protein abundance was based on MS intensity values. For Ginno et al. (2018) study abundance was defined as the median MS intensity across six replicates (G1 and S phases, three replicates each), based on the “Chromatome Reporters” dataset. For Shi et al. (2021) study abundance was calculated as the median iBAQ value across three replicates. For Itzhak et al. (2016) study for proteins with a single isoform, the “Estimated copy number per cell” was used as the abundance value. For proteins with multiple isoforms, the median copy number across isoforms annotated as “mostly nuclear” was used. For Ugur et al. (2023) study abundance was calculated as the median of the three replicate raw log2 intensity values.

### 2.4. Protein physico-chemical properties

Intrinsically disordered regions (IDRs) were identified based on solvent-accessible surface area (SASA) profiles, smoothed using a 20-residue sliding window, calculated from AlphaFold2-predicted protein structures [50]. Regions were classified as IDRs if they contained at least four consecutive residues with accessibility values greater than 0.55, and were separated from other IDRs only by non-IDR fragments longer than four residues. Protein tails were defined as N-or C-terminal IDRs. Protein charge was classified as positive (>=1), negative (<=1) and neutral (0) according to the number of positive charged amino acids (lysines and arginines) and negatives (aspartates and glutamates). Protein tail mean net charge was calculated as a sum of amino acid charges averaged per each amino acid position (first 80 amino acid position was analysed). Protein tail charge profile was constructed with averaged charges per amino acid residue with rolling window (size=10 aa).

To get the individual number of protein molecules of the respective charge we adjusted the mentioned number of charged protein entries by protein abundance (PaxDb_PA values were used). Additionally we took into account the overall net charge of every protein, resulting in the analysis of the cumulative charge conferred by positively or negatively charged proteins.

To characterize differences between chromatin and cytoplasmic proteins with respect to their amino acid composition we used the Uniform Manifold Approximation and Projection (UMAP) nonlinear dimensionality reduction technique. Protein amino acid fractions were standardized using StandardScaler and reduced with UMAP with default parameters. The median values of the amino acid fractions in the chromatin proteins and the proteins that were uniquely localized in the nucleus or cytoplasm were compared using the Mann–Whitney test with Bonferroni correction. The statistical significance threshold was an adjusted p-value of less than 0.05. The fold enrichment of the amino acid fraction was calculated as the difference between the median value in chromatin or nuclear proteins and the median value in cytoplasmic proteins.

### 2.5. Protein structural characterization and protein domain analysis

Chromatin protein sequence coverage was assessed by identifying all experimentally resolved amino-acid residues for each target protein entry using structures deposited in the PDB database using PDB API (accessed on 15.01.2025). CATH structural domain assignments were used from [51] (CATH v4.3.0, AlphaFold v2). For Pfam domains we used Pfam-A regions from Pfam version 37.0, only the following type of regions were used: Domain, Family, Repeat, Coiled-coil. Data about protein domain annotation in other DBs (*e.g.*, InterPro, PANTHER, *etc*.) was obtained by InterPro API (InterPro version 103.0), only following types of domains were selected: domain, homologous_superfamily, family, repeat, coiled_coil. The percent of Pfam models presented in PDB was obtained by processing data from InterPro. TED domain annotation was downloaded using TED API v1 [52]. The domain annotations from different sources (TED, CATH and Pfam) were intersected if the length of intersection was more than half of the length of shortest annotation.

To look for and map TED domains and Pfam models to PDB or CATH database entries we used Foldseek Search (database versions with following labels were used “PDB100 20240101”, “CATH50 4.3.0”) [53] with easy-search module, –-exhaustive-search option and default parameters with following filtration by probability (estimated probability for query and target to be homologous (*e.g.*, being within the same SCOPe superfamily)) higher than 0.9 and query coverage more than 0.5. To characterize representation of chromatin proteins’ TED domains in PDB, the best match PDB structure (with the FoldSeek highest probability score) for every TED domain was identified. TED domains were then classified by the level of sequence identity with their best PDB match. The taxa of the best matches were aggregated into several groups: ‘Homo Sapiens’, ‘Mammalia’, ‘Other Vertebrata’, ‘Protostomia’, ‘Viridiplantae’, ‘Fungi’, ‘Archaea’, ‘Bacteria’, ‘Viruses’, ‘Other’, ‘N/A’ (synthetic constructs or unclassified sequences). In the case of Pfam models, for each Pfam model sequence the best match by probability was selected, sequence identities for each Pfam model were averaged by median value. Additional searches of TED domains in CATH were also performed by sequence search using CATH/GENE3D annotations.

Novel structural domains (potentially representing new types of structural superfamilies or protein folds) were defined as TED domains that could not be matched to any known protein structure or protein structure superfamily (by FoldSeek search in PDB and CATH50). For these domains available functional or other annotations were downloaded from InterPro (only matches where more than half of the TED domain was mapped to the annotated region were considered). DeepFri [54], a Graph Convolutional Network for predicting protein functions, was used to predict GO MF and GO BP terms for novel structural domains by both sequence (CNN model) and structure (GCN model).

Domain diversity of a SimChrom-SL category was calculated as the number of different Pfam domain models found in the proteins belonging to the respective SimChrom-SL category, relative domain diversity – domain diversity divided by the number of proteins in the corresponding SimChrom-SL category.

EMVI-domains were defined as those that were found in multiple copies or in combination with another Pfam domain in at least three chromatin regulator proteins (defined as the following SimChrom-SL categories: Histone chaperones, Histone PTM erasers, Histone PTM writers, Histone PTM readers, Histone modification, Chromatin remodelers, Methylated DNA binding, DNA (de)methylation, RNA modification). EMVI-domains were manually classified based on the information currently available in the literature into the following functional subgroups: Histone methylation, writer; Histone methylation, eraser; Histone methylation, reader; Histone acetylation, writer; Histone acetylation, eraser; Histone acetylation, reader; Histone phosphorylation, writer; Histone binding; DNA binding; DNA methylation; PPI; Dimerization/oligomerization; Chromatin remodeling; RNA binding; Other. The histone modification subgroups were additionally grouped together according to the type of the respective histone modification *(e.g.,* histone methylation). For domain composition scatter plots (*e.g.*,interactive graphs available at: https://simchrom.intbio.org/#domain_composition) Pfam domain models found in more than five chromatin proteins were selected. The conditional probability of finding a corresponding domain A in a chromatin protein given that another domain B is already present was estimated as P(A|B) = P(A ∩ B) / P(B).

## 3. RESULTS

### 3.1. Sources of information about chromatin and nuclear proteins and their critical evaluation

Given the ambiguity in exactly defining the criteria that delineate chromatin proteins from non-chromatin proteins, in analyzing and comparing the sources of information about chromatin proteins we chose to expand our analysis to include nuclear proteins. Nuclear localization data provides an independent way of validation since at least interphase chromatin proteins are also expected to be nuclear proteins. Hence, as the first step of our study we used database and literature mining to analyze the currently available sources of information about *human* chromatin/nuclear proteins and evaluated their congruency through cross-comparison.

The relevant sources of information were grouped into three classes: function-oriented protein/gene annotation databases, protein localization studies/databases, MS-based studies relying on chromatin extraction (**Fig. 2**). **Suppl. Table ST1** contains a representative list of 63 sources compiled for this study that include both historic and state-of-the-art datasets. It can be seen that although in the post-genomic era the size and completeness of the chromatin related proteomic datasets has grown considerably, there is still a lot of variation (*e.g.*, recent MS-based chromatin studies list from ∼1000 to ∼4000 chromatin proteins, the coverage of high-throughput localization studies ranges from ∼1000 to ∼12000 proteins), warranting datasets’ validation, comparison and harmonization.

**Figure 2.**
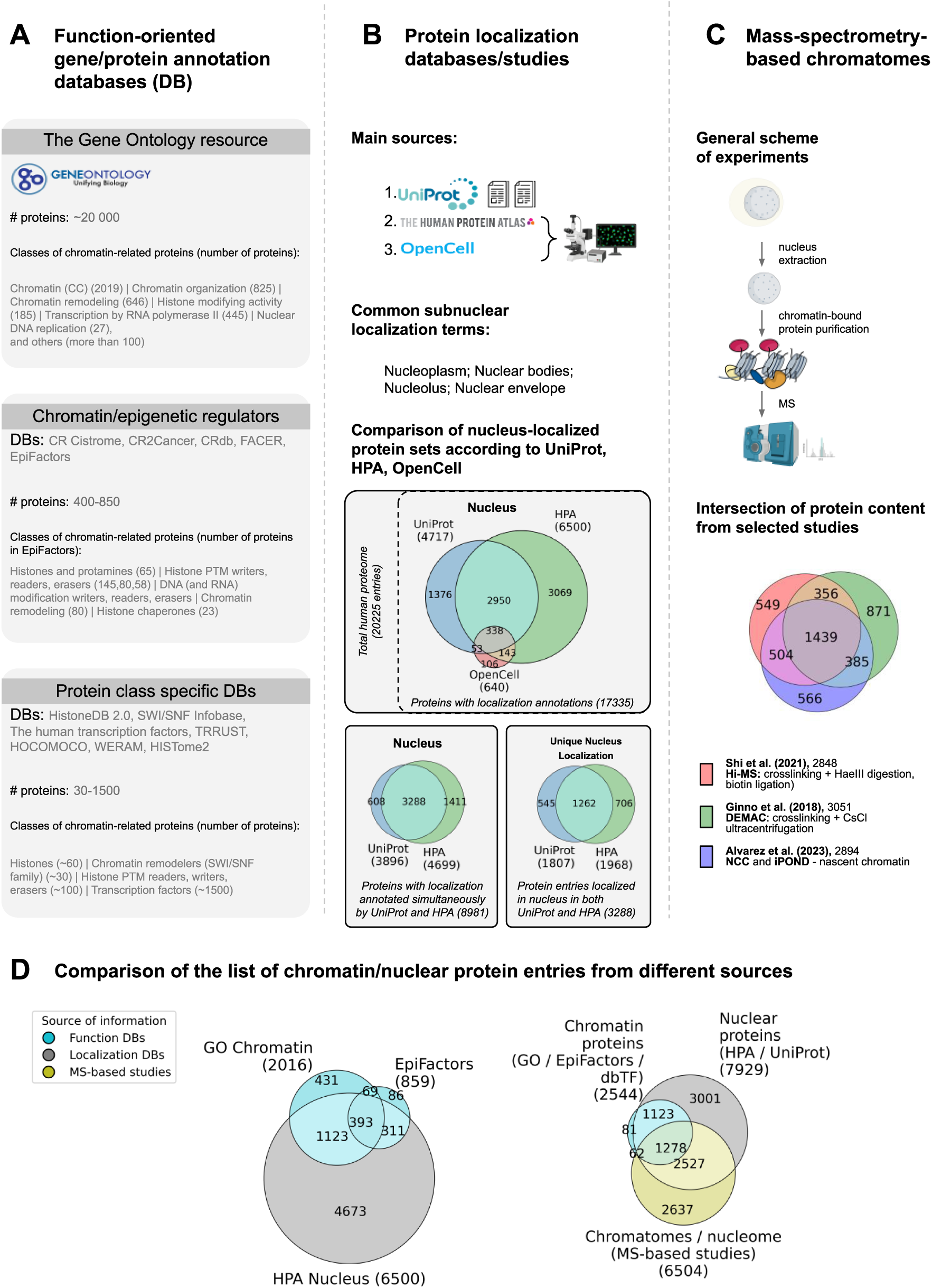
Source of information about chromatin proteins and their critical evaluation. **(A)** Function-oriented annotation databases include Gene Ontology, chromatin or epigenetic DBs (*e.g.*, EpiFactors), and protein class specific DBs (*e.g.*, HistoneDB, HISTome2). **(B)** Protein localization information was extracted from UniProt Knowledgebase (curated information about protein localization mainly from literature or computational analysis), the Human Protein Atlas (HPA, high-throughput immunohistochemical analysis of target proteins in fixed cells), and OpenCell (live-cell microscopy of proteins fused with fluorescent tags). The list of common terms describing subnuclear localization in the three databases is given. The Venn diagram **(top**) shows the intersection of nuclear proteins’ lists from the three databases (with filtration by evidence codes**/**confidence levels, see Methods). The Venn diagrams at **the bottom** show the congruence among annotations in UniProt and HPA (see text). **(C)** The general scheme of mass-spectrometry-based studies for chromatin proteins’ identification (left), the Venn diagram of chromatin proteins’ lists from four different MS-based studies. **(D)** Comparison of chromatin/nuclear proteins’ lists from different sources. **Left**: data from function-oriented DBs (proteins annotated by the term “Chromatin” in GO database and from EpiFactors database) versus localization databases (protein from the HPA database annotated by the term “Nucleus”), **right**: chromatin MS-based studies (proteins identified by three or more studies from the six analyzed chromatin MS-based studies, see Methods) versus data from function-oriented (proteins annotated by GO term “Chromatin”) and localization databases (HPA term “Nucleoplasm”).

For our detailed analysis we chose (1) a number of general (GeneOntology) and specific (*e.g.*, EpiFactors [43], The Human Transcription Factors [55], Histone Database [56], *etc*.) protein/gene annotation databases or datasets that contain information about protein function (see **Fig. 2A**), (2) three major databases and/or high-throughput experimental studies that provide protein localization data (UniProt, HPA, OpenCell) (see **Fig. 2B**), and (3) eight state-of-the-art MS-based chromatin/nucleome studies (see **Fig. 2C**, **Table 1**). Below we proceed with presenting the results of our assessment of each class of information sources, followed by cross-comparing data between them.

Among the protein/gene annotation databases the GeneOntology (GO) database stands out as a comprehensive attempt in describing the functions of gene products in an ever growing number of organisms [35,36]. Within the GO framework genes are annotated according to their involvement in certain molecular functions, biological processes, and cellular components. The ontology itself forms a complex interlinked hierarchy with more than 40,000 GO terms and offers annotations to nearly the entire human reference proteome. However, despite its apparent comprehensiveness the GO database could not *per se* provide answers to the questions that were instrumental to this study, namely, to provide a set of chromatin genes/proteins and a relatively simple functional classification of these proteins that could be used for further analysis.

The GO cellular component term “chromatin” is defined broadly as “the ordered and organized complex of DNA, protein, and sometimes RNA, that forms the chromosome” and encompasses around *2000* proteins. Comparisons with other databases suggest that this number is a rather conservative estimate. For instance, up to 528 proteins listed in specialized databases of epigenetic factors (EpiFactors) and transcription factors (GO catalogue of TFs [44]) are missing from this set (see **Suppl. Fig. SF2_1A**); concomitantly the HPA protein localization database suggests that there are around *6000* proteins located in nucleoplasm (see **Suppl. Fig. SF2_2A**). Furthermore many proteins annotated by GO terms that are *bona fide* related to chromatin (*e.g.*, chromatin binding) are missing from those annotated by the GO term “chromatin” (see **Suppl. Fig. SF2_1C**). This stems in part from the complexity of the relations between the GO terms belonging to different annotation aspects, in this example the terms “chromatin organization” and “chromatin remodeling” are not connected to the term “chromatin” within the ontology tree. The manual search and identification of the chromatin related GO terms and relevant proteins is challenging because (1) of the shear number of GO terms (*e.g.*, the word “chromatin” is found in the names of more than 60 terms, see **Suppl. Fig. SF2_1C**), (2) the fact that apparently relevant terms may include also non-chromatin associated entries (*e.g.*, transcription may also include mitochondrial transcription), (3) the fact that terms describing certain historically established chromatin protein groups may be missing (*e.g.*, histones, HMG proteins), (4) the fact that GO database is not chromatin-specific and may not be up-to-date in certain aspects (*e.g.*, contain obsolete terms such as “nuclear matrix” or lack annotations for proteins that are available in recent literature reviews). **<u>Suppl. R&D</u> Sec. 1.1** provides further details and examples from our analysis.

A number of epigenetic/chromatin regulators/factors databases (*e.g.,* EpiFactors [43], CRdb [57]) provide carefully curated information about chromatin proteins that are involved in what is historically assumed to be molecular mechanisms of epigenetic regulation **(Fig. 2A, Suppl. Table ST1)**. However, they annotate only around 400-800 chromatin proteins, which is much less than is expected to be in chromatin (see **Fig. 2D**). Unlike GO, these databases introduce a rather simple classification of proteins (the respective categories are highlighted in **Fig. 3**), but lack many essential chromatin categories (*e.g.*, histones, histone chaperones, *etc.*). Protein class specific databases and datasets available in published papers provided an even more trustworthy but narrow sets of information about certain classes of chromatin proteins. Of particular impact here by the number of provided entries are the databases of transcription factors. Recent databases (*e.g.*, The Human Transcription Factors [55], GO catalogue of TFs [44]) comprise around 1500 transcription factors. Additionally, several chromatin-related protein classes have been reviewed in the literature but lack dedicated database resources [58–64].

**Figure 3.**
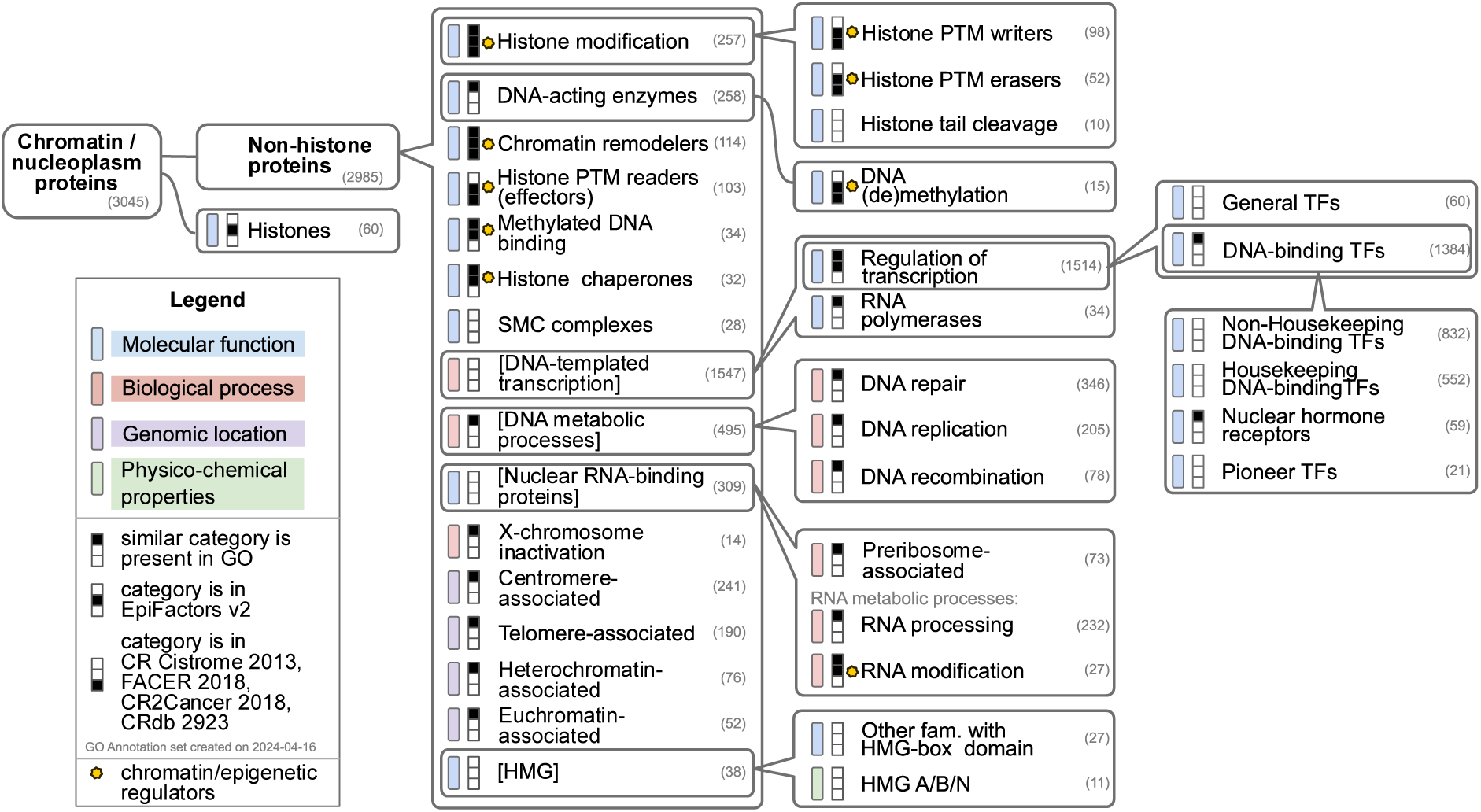
The SimChrom empirical chromatin classification ontology and the SimChrom chromatin proteins dataset. The hierarchical tree-like classification organizes 39 SimChrom categories. For each category the respective number of proteins from the SimChrom dataset is given in parentheses (proteins can simultaneously belong to more than one category with the exception of histones). The pictograms on the left of each category name provide the information about the presence of similar categories in the ontologies of other databases (see legend). The colored bars show whether the specific category was derived from grouping the proteins according to certain aspects: the similarity of their molecular functions, physico-chemical properties, involvement in similar biological processes or localization in similar genomic locations (see legend). Note: the latter annotations may deviate from GO annotation aspects.

The other alternative and powerful source of information about nuclear/chromatin proteins is localization databases and proteome-wide studies – UniPot, HPA and OpenCell projects are currently regarded as the most comprehensive and trusted resources on protein intracellular localization (see **Fig. 2B)**. From each resource we extracted the sets of proteins whose localization was annotated (only annotation with sufficient confidence levels was considered for our analysis – see **<u>Methods</u> Section 2.1.1**) as belonging to the nucleus or sub-nuclear compartments according to the localization ontologies specific to each resource. The detailed cross-comparison of the datasets is presented in **Suppl. Fig. SF2_2**, its interactive version (**Interactive Fig. 2** available at https://simchrom.intbio.org/#localization), and at length discussed in **<u>Suppl. R&D</u> Sec. 1.2** using **Suppl. Fig. SF2_2, <u>SF2_3</u>, <u>SF2_4</u>.** In summary there is a considerable degree of variation both between the sets of nuclear proteins, their sub-nuclear localization annotation and the annotation ontologies themselves between the resources.

It has to be kept in mind in the first place that localization annotation coverage is not complete – collectively the three resources cover 86% of the human reference proteome (*with sufficient confidence – see above*), while only 44% of proteins are simultaneously annotated by UniProt and HPA. The resources also differ by the number of localization annotations they provide on average for each protein (median number is two and one, for HPA and UniProt, respectively), suggesting HPA is more complete with respect to annotating multilocalization of proteins. Hence, although the protein space coverage by UniProt is larger compared to HPA (70% vs 60%), the nucleome provided by the former is considerably smaller (∼4700 vs 6500 proteins). Together the three resources annotate ∼8000 proteins as having nuclear localization (see Venn diagrams in **Fig. 2B**), which *amounts to 47% of proteins* that have localization information according to at least one resource. *Hence, as a rough estimate it is tempting to conclude that current protein localization databases suggest that around half of human proteins have some evidence of nuclear localization.* However, it has to be kept in mind that the congruency between the resources remains mediocre. Among ∼9000 proteins whose localization is simultaneously available in UniProt and HPA, ∼3300 are annotated as nuclear by both HPA and UniProt, while another ∼2000 are annotated as such only by one of the two resources (∼600 by UniProt and ∼1400 by HPA) (see **Fig. 2B**, lower left Venn diagram). The discrepancies are in part due to (1) incomplete annotation of multiple localization possibilities by the databases (among the ∼2000 proteins, ∼60% have a matching localization annotation between the databases other than nuclear), (2) potential biases in localization annotations (HPA tends to label nuclear proteins as vesicular proteins and UniProt tends to label nuclear proteins as secreted proteins and proteins of the extracellular matrix) (see **<u>Suppl.</u> <u>R&D</u> Sec. 1.2**, **Suppl. Fig. SF2_3**). Extrapolating the above estimates to the whole proteome (with all caveats in mind about the non-uniform annotation coverage of different protein groups) one can suggest that *between 35 to 60% gene protein products may be ascribed nuclear localization depending on the chosen degree of certainty*. Hence, a combination of the datasets provided by the resources may be used to construct reference nucleome datasets of varying confidence (see **<u>Results</u> Sec. 3.2**).

Many nuclear proteins were found to have multiple localization annotations belonging to different cellular compartments (see **Suppl. Fig. SF2_2B**, **<u>Methods</u> Section 2.1.1**, and **<u>Suppl. R&D</u> Sec. 1.2**). On one hand this reflects the functionally important property of nuclear proteins to shuttle between compartments. For example, many transcription factors and coactivators (*e.g.*, NF-κB, STAT, p53, TAF7, YAP/TAZ) regulate their action through cytoplasm/nucleus shuttling [65–67], even some histones, such as H2B, may relocalize to cytoplasm under stress and perform unconventional functions [68,69]. On the other hand, functionally irrelevant multiple localization annotations may arise due to experimental artefacts or suboptimal signal-to-noise thresholds, keeping in mind that all nuclear proteins are in fact synthesized in cytoplasm and imported to the nucleus. According to our analysis UniProt and HPA estimate separately that ∼50% of proteins with nuclear localization may be also localized in other compartments (∼40% in cytoplasm, 12%–22% in the endomembrane system), annotating around 48%-50% to be localized solely in the nucleus. However, once the annotations of UniProt and HPA are compared within the shared common set of proteins (nuclear localization annotation is available in both databases) it turns out that only for ∼40% of proteins the two databases reach consensus for their unique nuclear localization (see **Fig. 2B**, lower right Venn diagram). In other words (see **<u>Suppl. R&D</u> Sec. 1.2**), *approximately for every five proteins identified as uniquely localized in the nucleus by one database, it is likely that two of them will have non-nuclear localization annotation in the other database (per se or in addition to the nuclear localization)*. The same tendency was observed for the annotations of uniquely localized cytoplasmic proteins. The obtained estimates likely reflect the suboptimal specificity of the localization information provided by the databases (additional localizations of nuclear proteins may not be always captured) and potential presence of spurious localization annotations (artifacts or incorrect localization assignments). It is, however, non-trivial to deconvolute between these two types of errors.

Our analysis of sub-nuclear localization ontologies showed that the one of Uniprot is more diverse comprising 20 terms (this number includes chromosome localization which is distinct from the nuclear localization according to UniProt, although the majority – 84% – of chromosome proteins are also annotated as nuclear), while HPA and OpenCell comprize, 9 and 6 terms, respectively. However, in terms of annotation specificity only 19% of nuclear proteins in UniProt are annotated with sub-nuclear localization terms, while for HPA all of the nuclear proteins have some sub-nuclear localization (although 92% are considered a part of nucleoplasm, 33% bear localization annotations other than nucleoplasm). There are certain parts of the ontologies that do not match between the resources or to the state-of-the-art knowledge. For instance, HPA considers mitotic chromosomes as a part of nucleoplasm, while UniProt uses outdated “nucleus matrix” term. OpenCell is the only resource that explicitly considers “chromatin” as the possible localization for nuclear proteins, while UniProt explicitly lists “Chromosome” as the possible localization, which was in turn inherited from GO cellular compartment ontology where “chromatin” has a child-parent relation with the term “chromosome”. The above-mentioned discrepancies reflect the dynamic complexity of cellular organization, our constantly evolving understanding of nuclear organization, and the resulting difficulty in describing subcellular localization in a form of a simple hierarchical tree-like ontology. While the exact names may differ, all resources converge on the presence of the following localization terms: Nucleoplasm; Nuclear bodies; Nucleolus; Nuclear envelope. Among these terms the nucleoplasm localization is the one most related to chromatin proteins (according to HPA nucleoplasm is what is found within the nuclear membrane, but excludes nucleoli according to the respective localization ontology). If one interprets the definition of chromatin broadly (treats proteins that localize with the interphase chromosomes to be part of the chromatin “complex”) the set of proteins with nucleoplasm localization is a direct source of information about chromatin proteins. HPA lists around six thousand nucleoplasm proteins (**Fig. 2B**). The analysis of subnuclear multilocalization is available in **<u>Suppl. R&D</u> Sec. 1.2** and **Suppl. Fig. SF2_4**.

MS-based studies of chromatin extracts are another key source of information about the protein content of chromatin. Despite being the ultimate direct source of data about the composition of chromatin it unfortunately has certain limitations (see **<u>Introduction</u>**, and relevant reviews [17,70]). To gain quantitative understanding into the utility of MS-based studies for our goals we have selected data from several studies in human cell lines (see **Table 1** and **<u>Methods</u> Section 2.1.3**) for analysis. The selected datasets included *five* studies that aimed at total interphase chromatin characterization using different methods of chromatin purification and post-MS data analysis, *two* studies characterizing nascent chromatin, and *one* study characterizing total nuclear proteome. The more than twofold variation (from 1.5 to 3.5 thousand entries) in the number of detected chromatin proteins in various MS-based studies highlights the varying sensitivities of different chromatin purification/MS-detection setups (**Table 1**). The pairwise comparison of different chromatin datasets of comparable size (having around 3000 proteins) suggests that for any given set its fraction overlapping with any other set does not exceed 68% (see **Fig. 2C**). The number of chromatin proteins present simultaneously in all total chromatin datasets is 179 (**Suppl. Fig. SF2_5B**). These facts highlight considerable variation of MS-based data due to different sample sources and chromatin extraction techniques.

We next thoroughly analyzed these protein datasets through cross-comparison between themselves, comparison with protein localization data, and tested enrichment of different chromatin protein categories (according to SimChrom classification described in **<u>Results</u> Section 3.2**). The detailed analysis is provided as **<u>Suppl. R&D</u> Sec. 1.3**, and we only succinctly summarize our conclusions below. From 10% to 38% of proteins identified in MS-based chromatin datasets currently do not have any support from localization databases through their annotated nuclear localization (see **Suppl. Fig. SF2_5A,C**), suggesting that even for chromatin purification protocols based on protein-DNA cross-linking there still might be a certain degree of contamination with non-nuclear proteins, mainly cytoplasmic ones (see **Suppl. Fig. SF2_6**). Yet, MS-based techniques may have predictive power to identify new chromatin proteins that are not annotated in the localization databases. For example, among 195 proteins reported simultaneously by at least five out of seven chromatin MS-based studies we estimated that around ∼30% of proteins may have indications in the literature supporting their nuclear localization. MS-based studies are biased towards identifying the housekeeping proteins – more than 80% of nuclear/chromatin proteins reported by the MS-based studies were from the housekeeping pool, while the average expected fraction of nuclear housekeeping proteins is around 62% (see **Suppl. Fig. SF2_7A,B**). This is expected since many non-housekeeping proteins are conditionally expressed. However, MS-based studies tend to miss the housekeeping transcription factors too (and even to a greater extent non-housekeeping TF) apparently due to their low abundance and dynamic nature of interactions (**Suppl. Fig. SF2_7C, <u>SF2_8A</u>,** see also **<u>Results</u> Section 3.3.1** for discussion of chromatin protein abundance). MS-based studies also struggle to recover as separate gene products proteins with very similar sequences, *e.g.* canonical histone isoforms (see **Suppl. Fig. SF2_7C**, **<u>SF2_8B</u>**).

To finalize our analysis we compared the datasets from three types of data sources about chromatin proteins examined above (**Fig. 2D, Suppl. Fig. SF2_9**). One can see that localization databases are currently leading by the number of proteins that may be considered as chromatin proteins in the broad sense (*e.g.,* the proteins of the nucleoplasm). However, there is still limited congruence with the other data sources. For instance, 25% of GO “Chromatin” proteins are not localized in the nucleus according to HPA, moreover, of these 500 proteins, only 115 have any localization information in HPA. Notably, 42% (2699) of the proteins identified in MS-based chromatome and nucleome studies lack nuclear localization annotations in both UniProt and the HPA, whereas only 254 proteins remain entirely unannotated for subcellular localization in these databases.

Taken together our analysis of different chromatome data sources revealed considerable heterogeneity of information and limited congruence between the available datasets. The available functional databases while providing functionally supported data are either limited in scope or suffer from historically-contingent complexity and sometimes discrepancies in their classification ontologies that are not tailored to provide comprehensive straightforward information about interphase chromatin proteins. The localization databases is a powerful alternative source of information that can give an upper bound for the set of chromatin proteins (since they should have nuclear/nucleoplasm localization), provide a relatively reliable estimate of the lower bound for the number of nuclear proteins, however, they suffer from an incomplete coverage of the proteome-localization space and hence difficulties in estimating false-positive and false-negative annotation rates (keeping in mind the multi-localization of proteins) and limited congruence of subnuclear localization ontologies. The MS-based studies of chromatin extracts are the most direct source of information about chromatome, they may identify new chromatin proteins not annotated currently into the databases, however, they are limited in scope (many proteins are conditionally expressed or have low expression levels) and suffer from contamination with non-nuclear proteins.

### 3.2. The SimChrom chromatin protein classification, the SimChrom dataset and other reference datasets

Taking into account the advantages and disadvantage of different sources of information about chromatin proteins presented above, we aimed at constructing a reference set of chromatin proteins together with a classification ontology and several supplementary nuclear localization protein datasets that can be later used in analyzing the repertoire, abundance, functional, structural, and physico-chemical properties of chromatin proteins. Our aim was to create a relatively simple classification ontology that while potentially sacrificing the details will enable a holistic human-understandable overview of the chromatome (see **<u>Suppl. R&D</u> Section 1.1** for the discussion of GO complexity and ensuing challenges). The current version of SimChrom classification focuses on classification of chromatin/nucleoplasm proteins leaving aside the classification of the nuclear envelope proteins, which are historically not considered to be a part of chromatin. The SimChrom classification ontology was created by manually analyzing, critically evaluating, selecting and combining into a tree-like classification scheme information from (1) the historically established consensus on chromatin proteins classification (*e.g.*, histone, non-histone proteins, HMG-proteins [1]), (2) classification used in major databases of chromatin and epigenetic regulators (*e.g.*, EpiFactors, FACER), (3) classification used in the three aspects of Gene Ontology, (**Suppl. Fig. SF3_1**). Our hierarchical SimChrom classification is presented in **Fig. 3**. The majority of classification terms used in SimChrom was inspired by GO-based classification, yet only a small subset of terms was used. The main focus of the classification was to classify chromatin proteins according to their functions and biological processes that they are involved in, but genomic-location (which is also indirectly related to function) and physical properties (*e.g.*, high-mobility group proteins of A, B and N families) were also considered (**Fig. 3** highlights what classificational aspects are most relevant for each term using a color bar). The SimChrom ontology was developed simultaneously with the SimChrom dataset in an iterative manner by obtaining sets of proteins annotated by various GO terms, extracting them from literature and domain specific databases, manually curating, validating and filtering (see **<u>Methods</u> Section 2.2**). Only major splice isoforms of genes are included in SimChrom. The resulting SimChrom dataset contains 3045 proteins, is available as a **Suppl. Table ST5** and viewable in the **Interactive Fig. 3** at the SimChrom web-site (https://simchrom.intbio.org#classification). The descriptive details about the SimChrom dataset are available in **<u>Suppl. R&D Sec. 2</u>**.

In the default SimChrom classification (depicted in **Fig. 3**) every protein from the SimChrom dataset may belong to more than one SimChrom ontology category. This provides the needed degree of flexibility since many proteins indeed may *bona fide* belong to several categories due to their complex functional, physico-chemical or structural properties. However, in certain cases of holistic analysis an even simpler classification may be useful, which ascribes every protein to only one category. Such single label classification (**SimChrom-SL**) based on the same SimChrom ontology was also developed (see **<u>Methods</u> Section 2.2**, **Suppl. Fig. SF3_2**). Briefly, if the protein belonged to several categories by default it was ascribed to the category with the least number of other proteins (*i.e.* the most specific category for this protein) with functional categories taking priority (see **Suppl. Fig. SF3_2** for category priority order).

As auxiliary datasets based on the results of **Section 3.1** we have compiled several reference datasets of nuclear and non-nuclear proteins at different levels of support (depending on whether nuclear localization is supported by one or several localization databases), confidence (depending on the evidence codes and reliability scores provided by the databases), and also whether proteins are uniquely localized in the nucleus or have multiple localization in the nucleus and other cellular compartments (see **<u>Methods</u> Section 2.2.2**). The list of the datasets and their definition is presented in **Suppl. Table ST6**, the datasets are available for download in the **Interactive Table 2** at https://simchrom.intbio.org/#download. Instrumental to our further analysis will be the “nuclear localization consensus” (NULOC_CS) dataset – the set of nuclear proteins, whose nuclear localization is supported (with sufficiently good confidence levels) both by UniProt and HPA and does not contradict the data from OpenCell, and the “nuclear localization joint dataset with no evidence code filtering” (NULOC_JT_NECF) dataset – the maximally broad set of nuclear proteins, which includes proteins whose nuclear localization is supported by any of the localization databases at any levels of confidence. The NULOC_CS dataset contains 3296 entries, while NULOC_JT_NECF contains 8912 entries.

To evaluate the contents of our SimChrom dataset we performed its cross-comparison to the localization based datasets described above (NULOC_CS and NULOC_JT_NECF) (see **Suppl. Fig. SF3_3**). Detailed discussion of the results is provided in **<u>Suppl. R&D Sec. 2</u>**. Briefly, almost all SimChrom proteins had some evidence of nuclear localization (95% were present in NULOC_JT_NECF dataset, 60% in NULOC_CS dataset, see **Suppl. Fig. SF3_3**). For the SimChrom proteins that did not have high confidence support of nuclear localization (non present in NULOC_CS) GO enrichment analysis of SimChrom-exclusive proteins revealed minimal association with non-nuclear functions, with only a minor subset (∼10 centromere-associated proteins) linked to such categories (**Suppl. Fig. SF3_4**, **Suppl. Table ST8**). Moreover, no additional chromatin-related GO categories were found to be underrepresented in SimChrom, indicating its broad coverage of chromatin-associated functions (**Suppl. Table ST9**). 60% of SimChrom proteins were successfully identified in MS-based chromatomes and nucleomes (see **<u>Suppl. R&D</u> Sec. 1.3, Suppl. Fig. SF2_7)**. The remaining 40%, predominantly low-abundant transcription factors, were likely undetected due to their transient nature and dynamic interaction properties, which pose challenges for MS-based detection. Furthermore GO enrichment analysis of MS-derived proteins absent in SimChrom or nuclear reference sets did not reveal a *bona fide* chromatin-associated category (see **Suppl. Table ST7**). Together, these results support the quality of the SimChrom dataset, suggesting that SimChrom is sufficiently comprehensive in its coverage of chromatin-related proteins and its categories.

### 3.3. Analysis of the human chromatome

Equipped with the datasets described above we aimed at a comprehensive characterization of the chromatome, including characterization of it composition (numbers of proteins belonging to different chromatin categories), abundance (the number of individual protein molecules present in the cells), physico-chemical properties of the amino acid sequences of the proteins, their domain architectures and interaction patterns (including engagement in multivalent interactions). The full discussion of the results is presented in **<u>Suppl. R&D Sec. 3</u>** and **Suppl. Fig. SF4_1 – SF 4_4, SF5_1 – SF5_5, SF6_1 – SF6_3, SF8_1, SF8_2.** The sections below summarize our analysis.

#### 3.3.1. The chromatome composition and abundance of chromatin proteins

To understand chromatin functioning it is important to know the chromatome content not only in terms of the set of proteins associated with chromatin, but also in terms of their abundance (*i.e.*, the (relative) number of proteins per cell or organelle). The analysis of MS protein intensities from the experimental chromatome/nucleome studies discussed above, revealed a high degree of variability (see **Fig. 4A**, **<u>Supp. Fig. SF4_1</u>**). For instance, the estimated relative mass of histone proteins varied from 0.1% to 58% depending on the study, suggesting a high degree of bias due to different experimental techniques and analysis pipelines used to process raw mass spectrometry data (see **Fig. 4A)**. Hence, for further analysis we relied on the “whole-organism” protein abundance information available in PaxDb for *H. sapiens*, which for every protein reports its relative abundance in ppm (parts-per-million) [71]. See **<u>Methods</u> Section 2.1.3** and **<u>Suppl. R&D</u> Sec. 3.1** for the discussion of the quality and applicability of the data.

**Figure 4.**
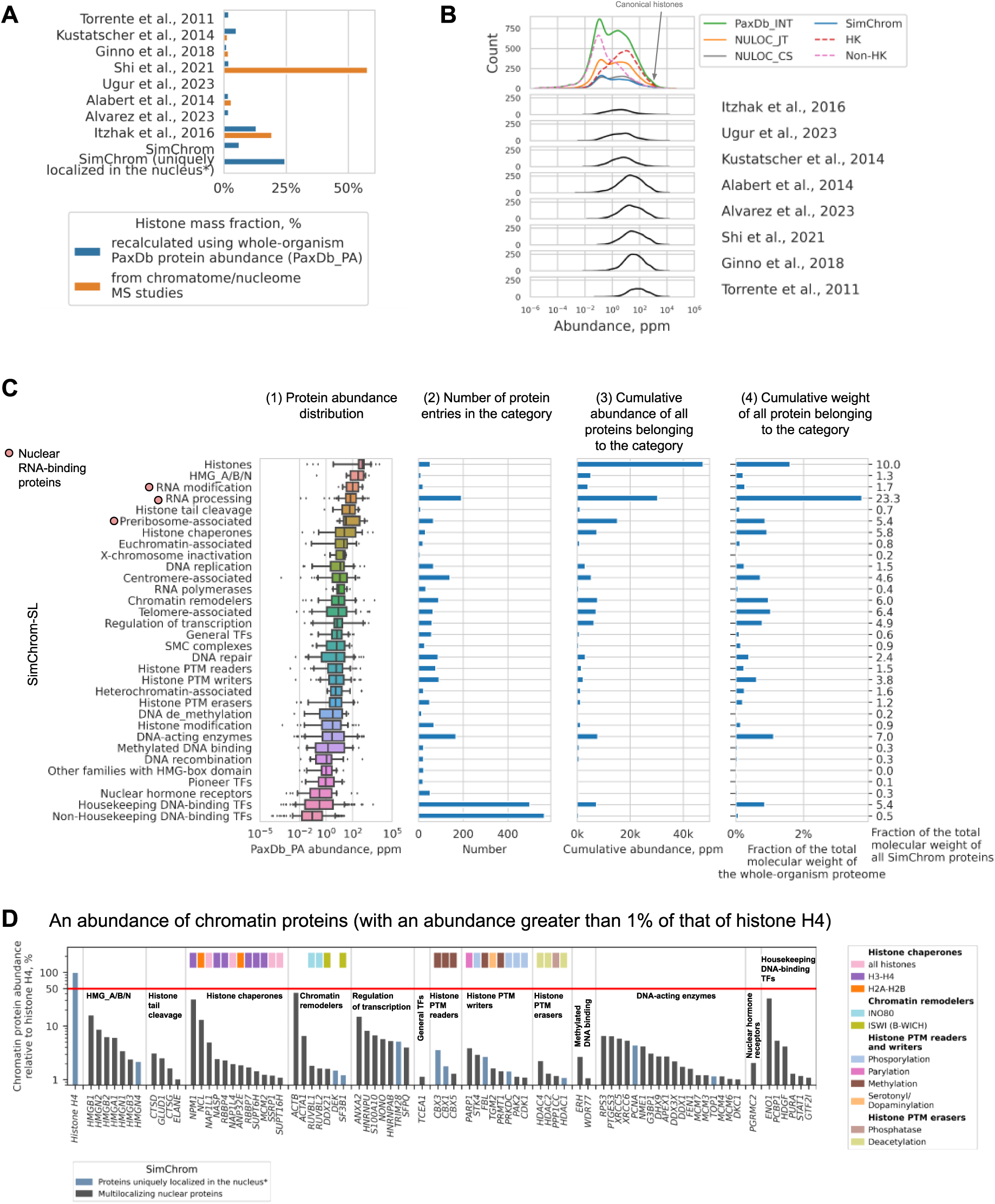
Analysis of the chromatome composition and abundance of chromatin proteins. **(A)** Fraction of histone proteins by mass in different datasets (according to experimental MS-based intensities and recalculated using abundance values from PaxDb_PA). (**B)** Distribution of proteins from different datasets according to their relative abundance values from PaxDb_INT. The distribution was constructed by taking the logarithm of the abundance values, making a histogram (bin size of 0.15) and smoothing it with a gaussian kernel for visual clarity. (**C)** Characterization of protein abundance (according to PaxDb_PA) for each SimChrom category and its contribution to the total abundance and mass of proteins. Each protein appears only in one category – SimChrom-SL classification is used. Subpanels show: 1 – distribution of chromatin proteins according to their relative abundance, box plots represent 25th and 75th percentiles, whiskers represent 5-95 percentiles, 2 – number of protein entries in each SimChrom-SL category, 3 – cumulative relative abundance of all proteins in the category, 4 – cumulative weight of all proteins belonging to the category relative to the whole human proteome and to the weight of all SimChrom proteins (numbers on the right side of the plot). The SimChrom categories are sorted according to the decrease of the median value of protein abundance. (**D)** The most abundant non-histone chromatin proteins. The barplot represents abundance (according to PaxDb_PA) for the most abundant non-histone chromatin proteins (with relative abundance of more than 1% of the histone H4 abundance). Proteins are referred to by the names of their genes grouped by SimChrom categories. The upper panel of colored rectangles provides additional functional classification (see legend). The red line at 50% abundance indicates the presumed abundance of nucleosomes (each containing two copies of the H4 histone).

The overall distribution of the human proteome with respect to protein abundance is bimodal (**Fig. 4B**), with roughly half of the human proteins having low abundance (LA) values below 1 ppm. The corresponding distributions of chromatin and nuclear proteins follow the same distribution (the fraction of LA proteins is 48% in SimChrom and 40-45% for NULOC_CS/JT). On the contrary, the sets of chromatin/nuclear proteins extracted from the MS-based studies demonstrate abundance distributions that are mainly unimodal and centered around high abundant proteins (**Fig. 4B**, **Suppl. Fig. SF4_1C**). The fraction of LA proteins identified in MS-based studies ranges from 1 to 27%, suggesting that while there is an overall bias towards highly abundant proteins, there is certain variation due to the methods of protein extraction and analysis (see **<u>Suppl. R&D</u> Sec. 3.1**, **Suppl. Table ST10**).

Our analysis of the number of house-keeping (HK) and non-house-keeping (non-HK) genes (see **<u>Methods</u> Section 2.1.4**) showed that the set of chromatin proteins is enriched in HK-gene products (the proteome wide fraction of HK genes is 40%, while in chromatin the fraction is 60%). It has to be kept in mind that around half (45%) of proteins in SimChrom are transcription factors, among which the non-HK TF constitute 60%. Hence, the non-TF fraction of SimChrom is even more enriched in HK-genes (74%). With respect to all-proteome distribution the increase in HK-proteins in chromatin is both due to the increase in LA and HA proteins (see **Suppl. Table ST10**). The bias towards HA proteins in MS-based studies of nuclear/chromatin proteins may be traced both due to the increased proportion of HK-proteins (which in turn have more HA representatives) and the difficulties in detecting the LA proteins among both HK and non-HK-proteins. The studies whose datasets are highly biased towards HA proteins mainly suffer from the latter problem (see **<u>Suppl. R&D</u> Sec. 3.1**).

We next aimed at understanding the abundance of different chromatin protein groups and individual chromatin proteins in the cell relying on our SimChrom-SL classification using PaxDb abundance data. The resulting diagrams depicting abundance variations of chromatin proteins, belonging to different SimChrom-SL categories, the number of proteins belonging to the respective categories, and the cumulative abundances (calculated both as the total number of protein molecules and the total molecular weight of protein molecules belonging to each SimChrom-SL category) are presented in **Fig. 4C**. To gain additional insights into the functioning of chromatin in **Fig. 4D** we plotted the abundance values of highly expressed chromatin proteins (abundance of more than 1% of the H4 histone abundance) belonging to SimChrom-SL categories of the “Molecular function” or “Physico-chemical properties” type. It is important to note that many chromatin proteins have additional localization in other cellular compartments, hence the presented data reflects the overall abundance of the chromatin proteins in the cell rather than their abundance in the nucleus (see **Suppl. Fig. SF4_3A** for an analogous analysis of chromatin proteins with unique nuclear localization).

As seen in subpanel 1 of **Fig. 4C** chromatin categories vary substantially by their median abundance from 0.09 ppm to 570 ppm and there is still considerable variation in the abundance values within the categories. The most abundant chromatin protein is histone H4 (∼11000 ppm), and it is convenient to measure the abundance of all other proteins in fractions of its abundance. For detailed analysis of histone protein abundance see **Suppl. Fig. SF4_2A,C** and **<u>Suppl. R&D</u> Sec. 3.1**. Briefly, the most abundant histone variants are H3.3, H2A.X, H2A.Z, the least abundant are H2A.B and H1.7 (see **Suppl. Fig. SF4_2C**). Despite the relatively small number of protein coding human histone genes (108), many of which code for identical sequences, the cumulative abundance of histone proteins exceeds that of all other chromatin protein categories even if proteins with multiple localization are taken into account (see panel 2,3 in **Fig. 4C**). However, when the total molecular weight of proteins belonging to different categories is compared, the relatively small size of histone proteins (median ∼15 kDa) results in them yielding the first place to RNA modification proteins (see panel 4, **Fig. 4C**). Collectively the cumulative weight of proteins belonging to “Nuclear RNA binding proteins” category (that combines Preribosome-associated, RNA modification, and RNA processing categories) amounts to 30.4% of all SimChrom proteins weight (4.8% of whole-organism proteome weight). However, many proteins from these categories are also localized in cytoplasm, and the major contribution to their cumulative molecular weight likely comes from the cytoplasmic fraction. Among the proteins uniquely localized in the nucleus (**Suppl. Fig. SF4_3**), the mass fraction of histones is in the first place (38%), however, the mass fraction of RNA processing proteins (around 70 uniquely localized proteins, including splicing factors, pre-mRNA binding proteins, etc.) remains high (27%) it part due to higher average molecular weight of proteins in this group (see **Suppl. Table ST11** for a list of these proteins).

Other functional chromatin protein groups (or groups with specific properties) <u>with high values</u> <u>of median abundance</u> include HMG A/B/N, histone tail cleavage, histone chaperones, RNA polymerases, Chromatin remodelers and other categories (see **Fig. 4<u>С</u>**). The high mobility group proteins (HMG A/B/N) are the second group after histones ranked by their median abundance, the estimated ratio of HMG proteins to nucleosomes is 1:8, 1:2, 1:3 for proteins of main HMG superfamilies, HMGA, HMGB, or HMGN proteins, respectively. Among histone chaperones the H3-H4 histone chaperone NPM1 and H2A-H2B histone chaperone NCL have the highest abundance, 32% and 13% of H4 abundance, respectively.

<u>The groups with least median abundance</u> are those related to “DNA-binding transcription factors” (pioneer TFs, nuclear hormone receptors, housekeeping and non-housekeeping TFs). Non-housekeeping DNA binding transcription factors have an abundance between 0.00008 and 23.9 ppm, suggesting their expression at minimal levels when averaged across all body tissues. The majority of housekeeping transcription factors is also expressed only marginally (median abundance 0.314 ppm). However, the abundance of certain proteins classified as housekeeping TF may reach 91.8 ppm (RURB1 gene, 0.84% of H4) for TF uniquely localized in the nucleus or 3667 ppm for proteins that have multiple localization (ENO1 gene, 33.6% of H4). *The DNA-binding transcription factor groups have the largest number of genes in SimChrom-SL (1385 genes in total), however, their contribution to the cumulative weight of chromatin proteins is rather small (6.3%)*. See **<u>Suppl. R&D</u> Sec. 3.1** for a full discussion of the abundance of other protein groups in SimChrom.

#### 3.3.2. Physico-chemical properties and amino acid composition

The physico-chemical properties of chromatin proteins encoded in their primary sequence were analyzed in comparison with those of the uniquely localized nuclear and cytoplasmic ones (SimChrom, NULOC_CS_UL, and CYTLOC_CS_UL datasets were used), see **Fig. 5C-I** and **Suppl. Fig. SF5_1 – SF5_5**. **Fig. 5A,B** summarizes the properties that were under study and the main differences of chromatin proteins from cytoplasmic ones.

**Figure 5.**
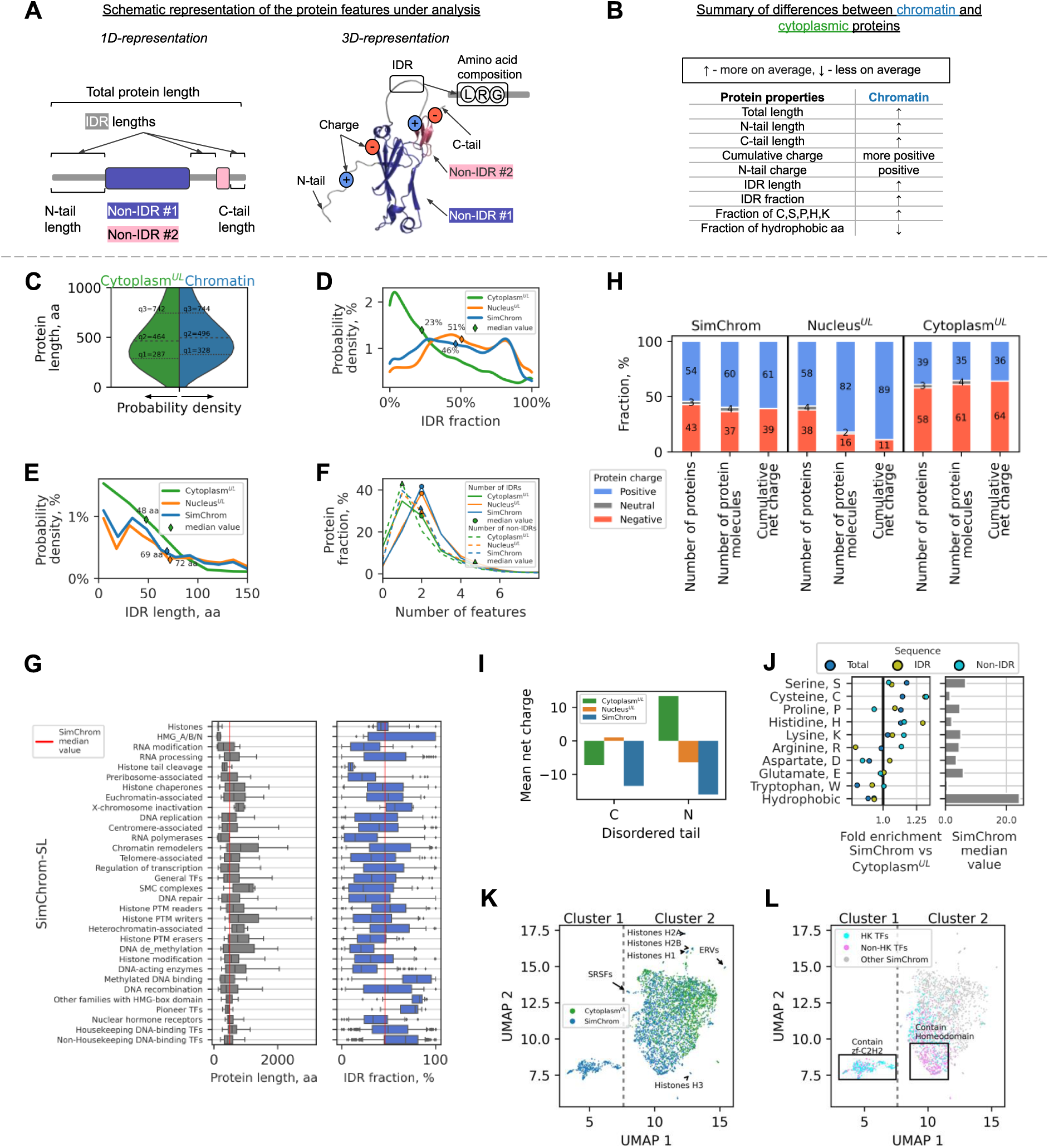
Physico-chemical properties and amino acid composition of chromatin proteins. *(**A**)* Schematic representation of the analyzed protein features/properties and (**B**) summary of the differences in properties of chromatin versus cytoplasmic proteins. Following datasets and designations are used in this figure: SimChrom – chromatin proteins, CYLOC_CS_UL (Cytoplasm^UL^ – uniquely localized cytoplasmic proteins), NULOC_CS_UL (Nucleus^UL^ – uniquely localized nuclear proteins). (**C-E**) Comparative distributions of chromatin, cytoplasmic and nuclear proteins with respect to their (**C**) protein length, **(D)** IDR fraction, **(E)** IDR length, (**F**) number of IDRs/non-IDRs. (**G**) The distribution of proteins by length and IDR fraction for individual chromatin protein categories (based on SimChrom-SL classification). Box plots represent 25th and 75th percentiles, whiskers represent 5-95 percentiles. (**H**) Ratio of the number of: (1) negatively and positively charged protein entries in the datasets (SimChrom, uniquely localized nuclear and cytoplasmic proteins), (2) individual protein molecules of the respective charge (estimated using PaxDb abundance data), and (3) ratio of the total net charges contributed by these molecules (calculated separately for all positively and all negatively charged protein molecules). (**I**) The average net charge of C– and N-disordered protein tails for different protein groups (see **<u>Methods</u> Section 2.4**). (**J**) Difference in amino acid composition of proteins (including subdivision into IDR and non-IDR regions) for chromatin proteins in comparison with cytoplasmic ones. The left panel represents statistically significant fold enrichment of median values (see Methods and **Suppl. Fig. SF5_4**). The right panel represents the median values of amino acid or group of amino acids fraction in chromatin proteins. (**K, L)** Grouping of chromatin and cytoplasmic proteins by their amino acids composition using UMAP dimensionality reduction technique. Different groups and individual proteins that stand out in the UMAP space are highlighted in panels **K** and **L** (mainly transcription factors). The dashed line denotes a visible border between clusters 1 and 2: cluster 1 mostly contains TFs with zf-C2H2 domains, whereas cluster 2 contains other chromatin proteins, including TFs with Homeodomain (see text for details).

First we analyzed the total protein length, and presence of intrinsically disordered (IDR) and non-IDR regions (**Fig. 5C-G, Suppl. Fig. S5_1**). The average chromatin protein is somewhat longer than a cytoplasmic one (median length value increases from 460 to 490 amino acids (aa)), however, there is considerable variation among the SimChrom categories (**Fig. 5G**). The smallest proteins are histones and HMG group proteins (median values of 130 and 109 aa), while the SMC complexes and chromatin remodelers are on average much longer (median values of 1096 and 835). The longest chromatin proteins reach the length of five thousand aa (*e.g.*, preribosome associated protein MDN1 – 5596 aa, histone methyltransferase KMT2D – 5537 aa). It has to be kept in mind that the largest contributors to all average characteristics of chromatin proteins are transcription factors (TF) because of the number of proteins in these categories. Interestingly, the non-HK TF are on average shorter (median 432 aa) than cytoplasmic proteins, while HK TF are longer (median 563 aa). Chromatin and nuclear proteins differ drastically from cytoplasmic proteins in the fraction of their length occupied by IDR regions (∼50% vs ∼25%). This is both due to the increase in the number of IDRs (median is two vs one) and the length of individual IDRs (∼70 vs ∼50 aa). Interestingly, non-HK TFs have an especially high fraction of IDRs (median ∼68%, and contribute a pronounced peak in the distribution at around 80%, **Fig. 5D, Suppl. Fig. S5_1B**), some other groups are also especially rich in IDRs (HMG, pioneer TFs, Methylated DNA binding proteins). More details can be found in **<u>Suppl. R&D</u> Sec. 3.2**.

It is generally assumed that chromatin proteins are on average positively charged to compensate for the negative charge of the genomic DNA. In our datasets the entries related to positively charged proteins dominate among nuclear and chromatin proteins, while the ones related to negatively charged proteins dominate among the cytoplasmic ones (see **Fig. 5H**). The dominant charge group in each case consists of around more than half of protein entries (54-58%), while the opposite one consists of 38-43% of protein entries. This metric, however, does not account for protein abundance or total positive and negative charge conferred by the proteins. The data adjusted for protein abundance (see **<u>Methods</u> Section 2.4** and **Fig. 5H**) suggests that among the proteins that are uniquely localized in the nucleus, positively charged protein molecules dominate (82% vs 16%). The total amount of net positive charge contributed by such proteins exceeds the net negative charge contributed by the negative ones (the ratio of the values is 89:11). Among different chromatin protein groups there are those that are significantly enriched in negatively charged proteins (see **Suppl. Fig. S5_1E**). Among the categories having the most number of negatively charged proteins are Histone chaperones (84%), RNA polymerases (76%), Histone PTM erasers (71%). These categories, however, have many proteins that are localized both in the nucleus and the cytoplasm. This surprising presence of many negatively charged proteins in these categories is likely explained by their preferential association not with DNA, but rather with positively charged histones.

To further elucidate the peculiarities of charge structure in chromatin proteins we analyzed the length and average charge profiles of protein N– and C-terminal tails (**Suppl. Fig. SF5_2**). For chromatin proteins both N– and C-terminal tails are negatively charged, while for the proteins uniquely localized in the nucleus N-tail is negatively charged (see **Fig. 5I**). Interestingly there is a clear difference with the cytoplasmic proteins, whose N-tails are positively charged. The average charge of the N-terminal tails was –16 for SimChrom proteins, –6 for nuclear proteins, and +13 for cytoplasmic ones. The analysis of the N– and C-terminal tail charge for different SimChrom categories revealed that they varied between the different categories (**Suppl. Fig. S5_2E,F**). Histones had the most positively charged tails, while histone chaperones and HMGs had the most negatively charged ones. Transcription factors on average also had negatively charged protein tails. This is an interesting fact because most transcription factors are positively charged (**Suppl. Fig. S5_1E**).

We next set to analyze in detail the amino acid composition of chromatin/nuclear proteins with respect to cytoplasmic ones and the variability of amino acids composition between different groups of chromatin proteins (see **Fig. 5J,K<u>,L</u>**, **Suppl. Fig. SF5_4, <u>SF5_5</u>**, **Suppl. Table ST13**). To this end we first used the UMAP nonlinear dimensionality reduction technique to see if significant variations between chromatin proteins can be identified in the space of their amino acids composition. The resulting 2D projections onto the main UMAP components revealed that (1) chromatin and cytoplasmic proteins occupied overlapping domains on the 2D map, but with a visible shift between their centers, suggesting there is an overall difference in the average amino acid composition, (2) certain chromatin protein groups formed dedicated clusters on the map, suggesting significant distinctness in their composition (see cluster 1, 2 and outliers shown by the arrows in **Fig. 5K,L**). Further analysis revealed that in the 2D UMAP map transcription factors, containing zinc finger domains and homedomains formed distinct clusters (see **Suppl. Fig. SF5_3A,B**). The most distinct group (cluster 1) was **almost exclusively** (415 out of 422) composed of zinc-finger containing DNA-binding transcription factors (240 housekeeping and 175 non-housekeeping) with the median number of zinc-finger domains (ZFD) of around 10 (**Suppl. Fig. SF5_3C**). Zinc-finger containing DNA-binding transcription factors were also present in cluster 2, but the median number of zinc-finger domains (ZFD) in that cluster was only three, hence containing a lower proportion of amino acids specific to ZFD (**Suppl. Fig. SF5_3D**). ZFD are enriched in histidine and cytosine (see **Fig. 5J** and discussion below). Other protein groups that occupied distinct positions on the UMAP map, included (1) histones, (2) serine/arginine-rich splicing factors (enriched in serine and arginine), and (3) reverse transcriptases of endogenous retroviruses (enriched in isoleucine and threonine) (see **Fig. 5K**, **Suppl. Fig. SF5_3D**).

The detailed analysis of amino acids composition of different chromatin protein groups is presented in **<u>Suppl. R&D</u> Sec. 3.2**. Briefly, among the top four enriched amino acids in chromatin proteins are serine, cysteine, proline, and histidine (**Fig. 5J**). The enrichment of cysteine and histidine is solely contributed by the ZFD of transcription factors (**Suppl. Table ST13**, **Suppl. Fig. SF5_4B**, **Suppl. Fig. SF5_5A**). The total enrichment of serine and proline in chromatin proteins is attributed due to their enrichment in the non-IDR regions (relative to IDR and non-IDR regions of cytoplasmic proteins), and more importantly due to the higher proportion of IDR regions in chromatin proteins (46% vs 23%) that in turn have a considerably higher proportion of these amino acids than non-IDRs (**Suppl. Table ST13**). Serine was also enriched in non-IDR regions globally, while the enrichment of proline in non-IDRs was observed only in a few categories (*e.g.*, HMG-proteins) (**Suppl. Fig. SF5_4H**).

The enrichment of positively charged amino acids is only statistically significant for lysine, but not for arginine, and the enrichment is relatively moderate (1.03 in chromatin) (**Suppl. Fig. SF5_4A**). Arginine is highly enriched in non-IDRs, but it is the most depleted amino acid in IDRs of chromatin proteins *versus* the respective regions of the cytoplasmic ones. The depletion of negatively charged amino acids in chromatin/nuclear proteins is statistically significant for aspartate (fold enrichment is around 0.9), while the depletion of glutamate is statistically non-significant. Interestingly, aspartate is enriched in IDRs and significantly depleted in non-IDRs. This suggests that the increased positive charge of chromatin/nuclear proteins has its main contributions in the depletion of aspartate and enrichment of arginine in non-IDRs, and moderate global enrichment of lysine.

Among the most relatively depleted amino acids in chromatin/nucleus are hydrophobic aliphatic amino acids, they are relatively rare in IDRs and hence the large proportion of IDRs in chromatin proteins accounts for their lower total fraction (**Suppl. Fig. SF5_4F,I**). Tryptophan, which is the rarest amino acid (∼1% in proteins), is the most depleted amino acid on average in chromatin/nuclear proteins and almost in all chromatin categories, except for a few.

#### 3.3.3. Domain composition of chromatin proteins and identification of new structural domains

Next we set out to systematically analyze the available data on structural characterization, domain annotation and domain composition of chromatin proteins. We specifically explored the structurally uncharacterized portion of the chromatome (the “dark” proteome) and identified potential new structural domains that are predicted by AI-based protein structure prediction tools (see **Fig. 6A**).

**Figure 6.**
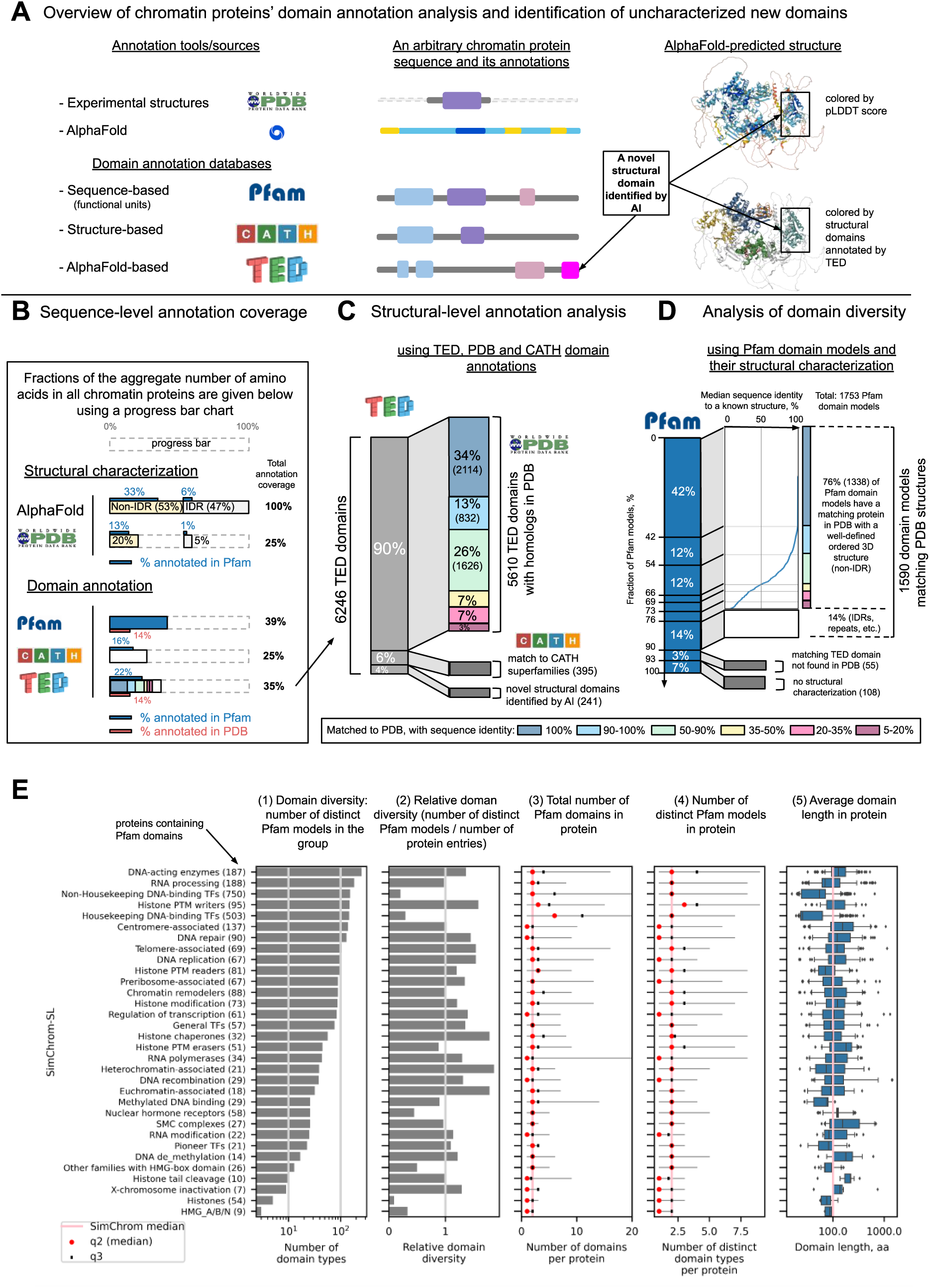
Domain composition of chromatin proteins and identification of new structural domains. **(A)** A schematic overview of chromatin proteins’ domain annotation analysis and identification of uncharacterized new domains. Sources of annotation and typical annotation patterns for an abstract protein are schematically outlined. A structure with a novel domain identified using AI-based annotation pipeline implemented in TED resource is shown on the right. **(B)** Cumulative annotation coverage of all chromatin protein sequences combined at the amino acid level via different resources. Annotation coverage with experimental structures in the PDB database, the AlphaFold database, and three domain annotation databases (Pfam, CATH, TED) is presented. For AlphaFold and PDB additional information about the fraction of annotated amino acids belonging to IDRs and non-IDRs is depicted (see **<u>Methods</u> Section 2.4**). For all annotations additionally the fraction of amino acids belonging to annotated Pfam domain models in the Pfam database is also depicted, for Pfam and TED additionally the fraction of amino acids resolved in PDB is also depicted. **(C)** Analysis of the structural domains in chromatin proteins identified by the TED resource via AlphaFold-based algorithm. The number and fractions of structural domains that have matching structures in the PDB database at various levels of sequence identity are depicted. The structural matches were identified via FoldSeek (see **<u>Methods</u> Section 2.5**). For those domains that were not matched to PDB structures directly a few were annotated by CATH (depicted in orange), the remaining fraction (depicted in magenta) represent novel structural domains present only in TED. **(D)** Analysis of functional domain diversity in chromatin proteins as identified by the Pfam database. 11147 domains belonging to 1753 Pfam domain models were identified. The plot characterizes domain models with respect to the availability of a matching structure in PDB (the median sequence identities of the matches between the chromatin proteins’ domains belonging to the respective Pfam model and their best structural match in the PDB database as identified by FoldSeek are shown), an annotated TED domain, or otherwise the absence of structural characterization (see **<u>Methods</u> Section 2.5**). **(E)** Analysis of functional domain diversity in chromatin proteins as identified by the Pfam database for proteins belonging to different chromatin categories according to SimChrom-SL classification. Subpanels 1-5 represent various characteristics.

Historically, protein domains are loosely defined as evolutionary conserved units with similarities at functional, structural and/or sequence levels [72]. Related individual protein domains may be grouped and aligned to produce domain models, catalogued and annotated by a number of resources/databases such as PFAM [73], CDD [74], CATH [75], InterPro [<u>76, p20</u>], *etc* (see **<u>Suppl.</u> <u>R&D</u> Sec. 3.3** for a thorough discussion). The ultimate experimental structural characterization of chromatin proteins is available in the PDB database, however, recent progress in protein structure prediction spurred by AlphaFold resulted in new approaches to the structural characterization and discovery of new structural domains (*e.g.*, as implemented in the TED database used below [52]) (**Fig. 6A**).

**Fig. 6B** shows the fractions of the aggregate number of amino acids in all human chromatin proteins (referred below to as “aggregate chromatome sequence”, or ACS) which are structurally characterized or have domain annotations in different databases. Detailed discussion is available in **<u>Suppl. R&D</u> Sec. 3.3**. Briefly, despite recent tremendous progress in structural biology many human chromatin proteins still lack direct structural characterization. On one hand only 25% of ACS can be mapped directly to PDB structures and 25% can be mapped to known structural protein superfamilies (through CATH). On the other hand, AlphaFold 2 identifies 53% of ACS as belonging to non-IDRs, and TED predicts that 35% of ACS belong to domains having well-defined 3D structures. The latter is a conservative estimate of structurally characterizable ACS, since both partially ordered and disordered regions can become ordered in protein-protein complexes. For example, 9% of ACS is available in PDB (where protein complexes are present) while not being annotated with TED domains (which rely on single protein chain structure predictions). This is in line with the fact that among 6246 TED domains found in chromatin proteins almost half of them (42%) are directly covered by the PDB database. However, the majority of other domains (56%) can be matched to a PDB structure of a homologous protein at various levels of sequence identity (from 99% to 5%, see **Fig. 6C**). The majority of these homologous domains are in fact different paralogous sequences found within human genes (even for domains with sequence identity of 35-50% the fraction of human sequences among the matches was 51%), for matches with sequence identity above 35% the second largest contribution came from structures of mammalian homologues, for matches with sequence identity below 35% significant contributions were from structures derived from proteins of fungi, protostomia and bacteria (see **Suppl. Fig. SF6_1A** for details). Additionally, 6% of TED domains that lacked direct hits among the PDB structures were mapped to protein structural superfamilies in the CATH database. The remaining 4% (241) represented domains could not be matched to any known protein structure or protein structure superfamilies and potentially represent new types of structural superfamilies/folds. These domains are presented in **Suppl. Table ST14** (see also **Interactive Table 3** at https://simchrom.intbio.org/#novel_structural_domains), ranked via their structural complexity by the number of their secondary structure elements. Among these domains, 123 domains have annotations in Pfam or other domain annotation databases present in InterPro, leaving 118 domains that are completely without annotations. The latter domains belong to 106 chromatin proteins, which may be considered as prospective new targets for experimental studies of their function and structure. Among such proteins are, for example, (1) a protein encoded by the GTF3C1 gene (it has a previously unannotated and uncharacterized structural domain with a length of 233 amino acids, see detailed characterization in **Suppl. Fig. SF6_2A**), (2) the globular domain of the testis specific linker histone H1.7, which has a quite different sequence from other H1 proteins resulting in a predicted structure that has a different topology (the “wing” of the globular domain consists of three beta-sheets rather than two [77], see **Suppl. Fig. SF6_2B** and **<u>Suppl. R&D</u> Sec. 3.3**).

We used the sequence-based Pfam domain annotation to characterize the diversity of different types of evolutionary related protein domains (hereafter referred to as Pfam domain models or Pfam domain types) found in chromatin proteins and typical domain composition thereof. In total 1753 different Pfam domain models matched various parts of chromatin proteins (**Fig. 6D**). 42% of these were considered fully structurally characterized, *i.e.*, every individual domain in chromatin proteins belonging to these models can be found in PDB. 34% of domain models are partially characterized – their domains could be matched to a PDB structure of a homolog (using FoldSeek, see **<u>Methods</u> Section 2.5**). 14% of these Pfam domain models were not matched by FoldSeek to PDB structures with our strict criteria (see **<u>Methods</u> Section 2.5**), but could be still identified in PDB via sequence search methods – these represented more flexible domains with IDR regions, repeats and coiled-coils (34 Pfam models), DNA-binding motifs, *etc*. 3% (55 domain models) could be matched to structural domains predicted by AlphaFold and found in the TED database. These represent prospective targets for validation with structural biology methods and further investigation of their interactions. For instance, among these domain models are domains, potentially associated with chromatin remodeling (SANTA, zf-C3Hc3H), histone PTM writing (DUF7030, COMPASS-Shg1), zinc fingers (zf_CCCH_4, zf-LITAF-like, zf-WIZ, SWIM), *etc*. 7% of Pfam domain models currently have no structural information that can be assigned either through the PDB or TED databases.

We next analyzed the diversity of Pfam domain models in various SimChrom-SL protein categories (**Fig. 6E**, subpanels 1,2) and the domain content of individual proteins belonging to these categories (**Fig. 6E**, subpanels 3-5). Detailed discussion is available in **<u>Suppl. R&D</u> Sec. 3.3**. Briefly, the number of distinct Pfam domain models found in chromatin proteins (∼1700) is comparable to the number of chromatin proteins (∼3000), at the same time an average chromatin protein usually contains two Pfam domains representing two different domain models. The majority of Pfam domain models are present only in a single chromatin protein, but there are also those that are present in dozens or even hundreds of proteins (**Suppl. Fig. SF6_1B**). Certain chromatin groups stand out in terms of their domain composition in some aspects: the number of individual domains is high in housekeeping TF (due to ZFDs); transcription factors, histones and HMG proteins are relatively poor in their domain diversity (*i.e.*, the proteins in these categories harbour a limited number of distinct Pfam domain models); histone PTM writers on average have domains belonging to three different domain models (while this number is one or two for all others). Still a considerable number of chromatin proteins may harbor domains belonging to several domain models. DNA-acting enzymes, histone PTM writers, chaperones, remodelers, transcription factors may have as much as 8-9 Pfam domain models present in their sequence (see **Suppl. Table ST15**). There are 118 chromatin proteins harboring at least five different domain types (see **Suppl. Fig. SF6_1C**, leftpanel). This highlights the multivalency of protein interactions in chromatin, keeping in mind that many proteins further form protein-protein complexes increasing their interaction potential (see next section). The average individual domain length in chromatin proteins is around 65 amino acids (the median is 28 aa), however, this number is biased by the presence of many zinc-finger domains (around 22 aa in length). Subpanel 5 in **Fig. 6E** gives a more balanced view for each SimChrom category. For the majority of protein groups the median domain length in protein is around 100 amino acids (mean is 137, median is 134). Only 70 chromatin proteins had no domain annotation at all.

The birds-eye view of the most frequently matched Pfam domain models in proteins of various functional SimChrom-SL categories is presented in **Fig. 7**. The data is presented for domain models that occur in at least five chromatin proteins and in at least 10% of proteins in a category (the threshold for data point depiction is 5%). The comprehensive interactive analysis figure with the ability to alter these thresholds and switch between SimChrom and SimChrom-SL classifications systems is available at **Interactive Fig. 4** (https://simchrom.intbio.org/#domain_composition). In **Fig. 7** the following categories and their respective domains can be grouped revealing their partially shared domain composition: 1) the categories containing transcription factors and their zinc finger, homeodomains and KRAB domains form the most frequently occurring entities, 2) some chromatin regulators, such as PTM writers, readers, erasers and chromatin remodelers together with their Chromo-, Bromo-, and PHD domains.

**Figure 7.**
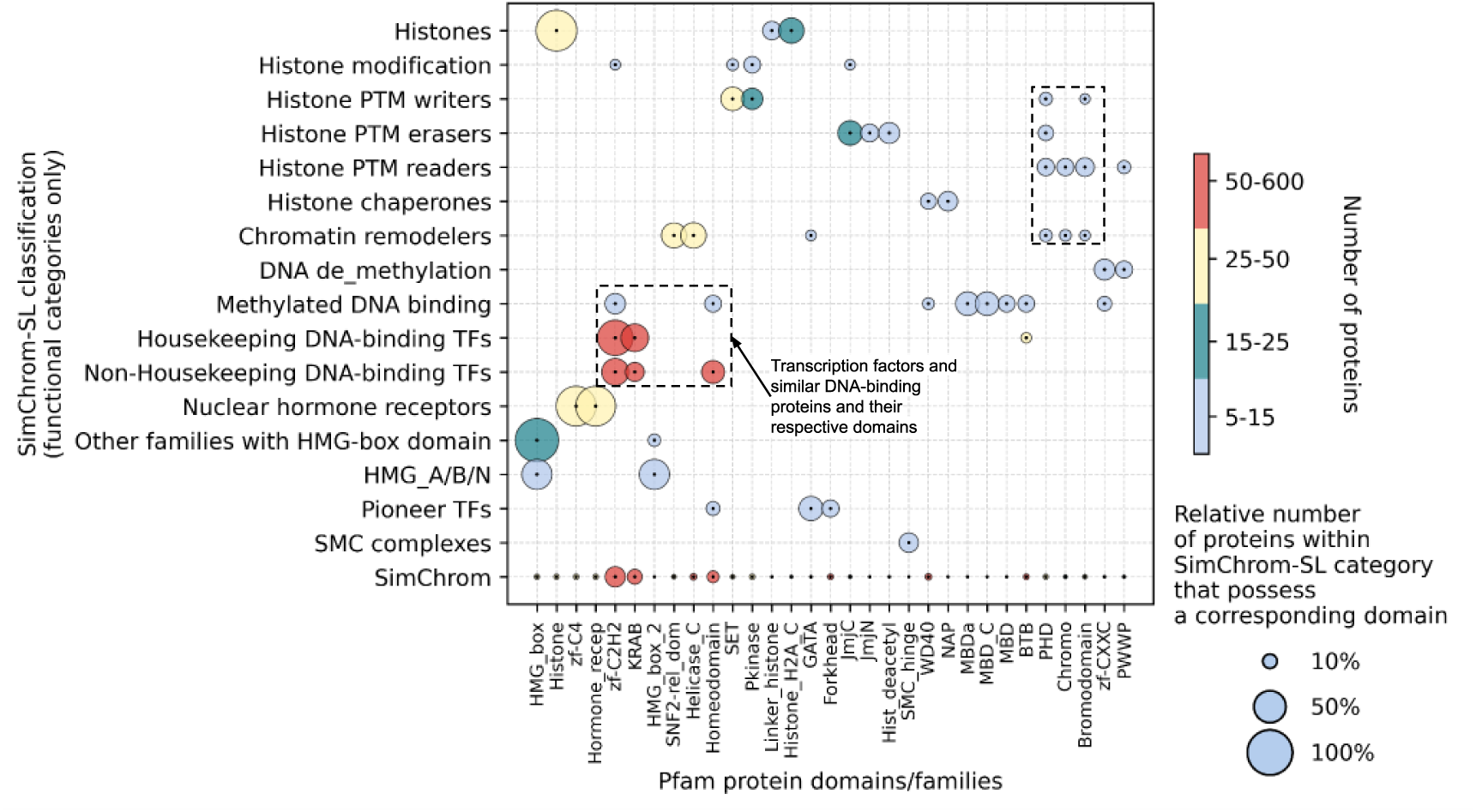
The most representative protein domains/families (according to Pfam) in proteins belonging to functional SimChrom categories. The dashed rectangles highlight the presence of particular groups of domains in certain categories of chromatin proteins (left – transcription factors and similar DNA-binding proteins, right – various histone interacting and modifying proteins). The plot is based on SimChrom-SL chromatin protein classification; only Pfam domain models present in more than five proteins were considered. Only datapoints with the size of more than 5% are displayed. The full size plots based on both SimChrom and SimChrom-SL classifications are available as **Interactive Fig. 4** (https://simchrom.intbio.org/#domain_composition).

#### 3.3.4. Multivalent interactions in chromatin protein

The presence of multiple domains (belonging to the same or different domain models) in chromatin proteins is a known feature contributing to their ability to engage in multivalent interactions (**Fig. 8A**) [10]. Below we present the analysis of such domains engaged in multivalent interactions (referred to as EMVI-domains hereafter) that are found in chromatin/epigenetics regulator proteins (see **Fig. 3** for definition of this group). These proteins often contain many domains (**Fig. 8B**). The median total number of domains found in chromatin proteins and in chromatin/epigenetic regulators is two. Nevertheless, many chromatin proteins contain more (16% – have three domains, 10% – four domains, 7% – five, six to fourteen – 10%). There are 409 Pfam domain models that are found in combination with other models or in multiple copies in at least one chromatin regulator protein. To limit our analysis to a manageable set of EMVI-domains, we selected those that were found in multiple copies or in combination with another Pfam domain in at least three chromatin regulator proteins (94 Pfam domain models in total), and from those we selected 59 domain models that we were able to manually classify based on the information currently available in the literature according to their functional binding modes. The following *functional groups* of domains were used: histone methylation/acetylation/phosphorylation, chromatin remodeling, histone binding, DNA binding, DNA methylation, protein dimerization/oligomerization, PPI, RNA binding. Histone post-translational modifications were further subdivided into readers, writers and erasers *functional subgroups* (see **Fig. 8C** and **Suppl. Table ST16** for the list of domains and their detailed classification). Domains involved in histone methylation are most present in chromatin regulators, followed by DNA binding, Histone acetylation, Histone phosphorylation and Chromatin remodeling associated domains (**Suppl. Fig. SF8_1A**).

**Figure 8.**
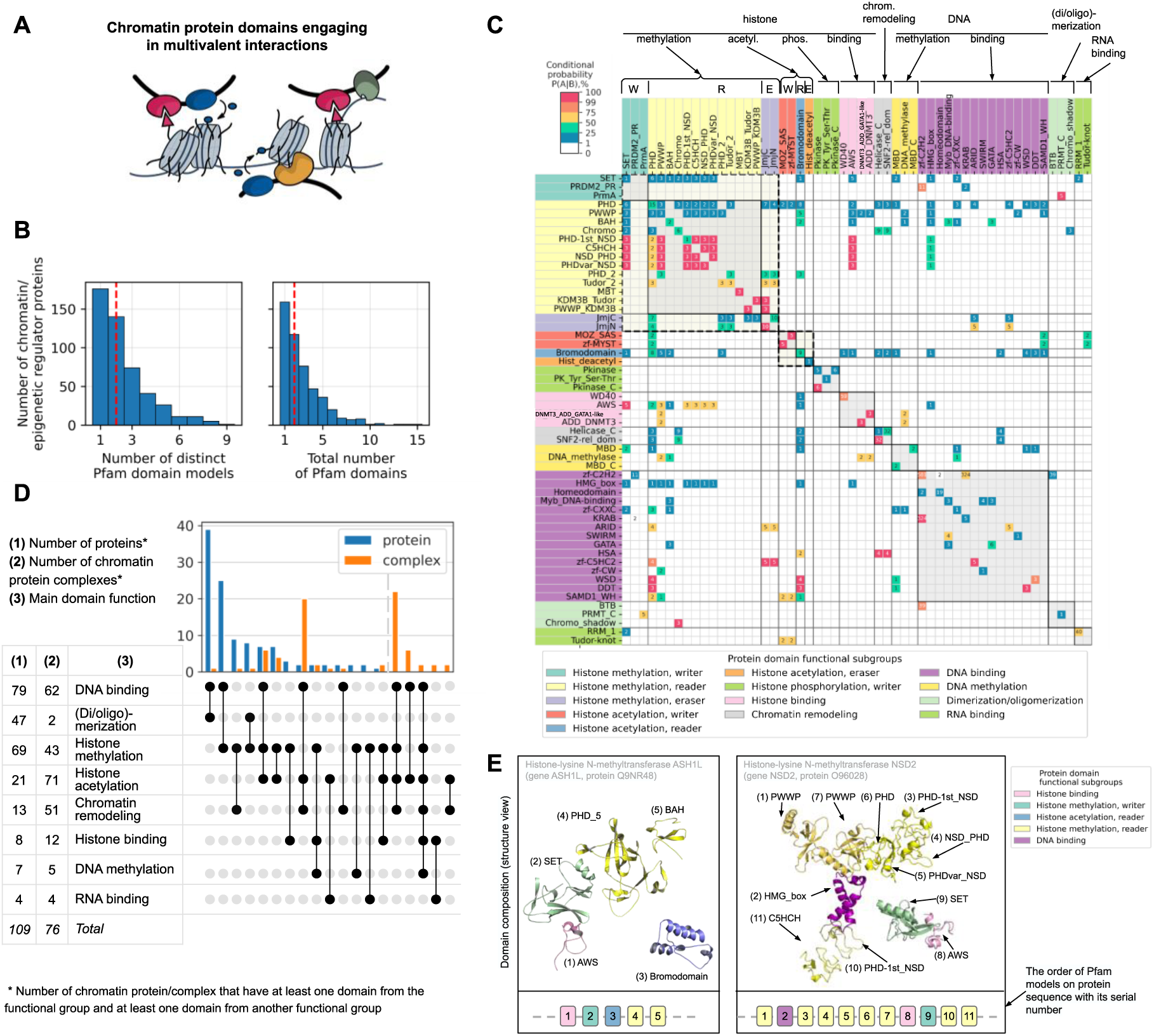
Analysis of multivalent interactions in chromatin proteins. (**A**) Schematic illustration of multivalent interactions. (**B**) Distribution of chromatin proteins from the chromatin/epigenetic regulators group with respect to the total number of Pfam domains (right) and distinct Pfam domain models (left), red lines indicate median values. The distributions of all chromatin proteins are shown in **Suppl. Fig. SF6_2C.** (**C**) Co-occurrence of domains corresponding to different Pfam domain models in chromatin proteins. Only domains found in proteins belonging to chromatin/epigenetic regulator groups are depicted (see **<u>Methods</u> Section 2.5** and **Fig. 3**). Domains are grouped into several functional classes (see description at the top of the plot). The values indicate the conditional probabilities of a domain in column (*A*) occurring alongside a domain in row (*B*) in chromatin proteins. Along the diagonal, data belonging to individual domain groups are highlighted with shading, dashed lines highlight groups associated with histone methylation or acetylation. The following abbreviations are used in the domain subgroup names of the latter: W – writers, R – readers, and E – erasers. (**D**) Co-occurrence of domains from different functional classes in chromatin proteins and protein complexes. Only combinations that are present in more than one protein or complex are shown, see full version in **Suppl. Fig. SF8_2**. (**E**) Examples of domain architectures in chromatin proteins containing the largest number of chromatin/epigenetic regulator domains. The top shows domains from 3D structures colored by their main function; the links between domains are not shown. The bottom shows the order of domains at the sequence level.

We analyzed the co-occurrence of selected EMVI-domains in all chromatin proteins. There were in total 851 chromatin proteins (589 of these are transcription factors) that had more than one EMVI-domain. The conditional probability of finding a corresponding domain A in a chromatin protein given that another domain B is already present was estimated and is presented in **Fig. 8C** (columns and rows correspond to domains A and B, respectively). The **Interactive Fig. 5** is available at https://simchrom.intbio.org/#domain_co-occurrence (also extends the analysis to unclassified potential EMVI-domains found in at least two chromatin regulator proteins). The matrix in **Fig. 8C** allows to trace the interplay between different domains employed in architectures of chromatin proteins. The largest groups of domains in **Fig. 8C** are those involved in histone methylation and DNA binding, suggesting that these mechanisms are the most represented and employed in chromatin functioning regulation. See **<u>Suppl. R&D</u> Sec. 3.4** for detailed discussion of the results. Briefly, in certain cases one can see 100% association between the presence of various domains in chromatin proteins. This may be due to direct structural interactions between the domains or likely due to functional reasons. Among the Pfam domains that co-occur with the most number of other different Pfam domains is the PHD domain (45 other domains), Bromodomain (38), SET (40) and PWWP (28) and chromatin remodeling Helicase_C (33) and SNF2-rel_dom (31), **Suppl. Table ST16**.

A more general view of multivalent interactions may be obtained if we trace the relationships within or between different functional groups of domains. One can see that domains from the same functional group (especially histone methylation) tend to co-occur (**Fig. 8C, Suppl. Fig. SF8_1B,C**) and may also be present in multiple instances in proteins (*e.g.*, in NSD2 there are nine histone methylation associated domains, see below). If a chromatin protein has a domain involved in histone methylation (either writing, reading or erasing) there is an estimated 38% chance that there will be another different functional domain from this group of domains (**Suppl. Table ST17**, **Suppl. Fig. SF8_1C**). For acetylation this estimated probability is 31%, for phosphorylation 18%. The associations between the occurrence of domains from different functional groups can also be observed. Domains involved in histone methylation (one of the most abundant groups by the number of Pfam models and the number of chromatin proteins) may be in a considerable number of chromatin proteins combined with other EMVI-domains (particularly DNA binding domains and histone acetylation), association with histone binding domains, chromatin remodeling, DNA methylation, (di/oligo)-merization and RNA binding domains was also observed (see **Suppl. Fig. SF8_2C**). The same can be said about domains involved in histone acetylation, although in a somewhat smaller number of cases, and with exclusion of their combination with dimerization domains. Notably, domains involved in histone phosphorylation were not found in combination with domains from other functional groups in our analysis. This may reflect an evolutionary strategy whereby combinations of histone methylation and acetylation evolved to delicately regulate gene expression at the epigenetic level, while phosphorylation remained as a more general mechanism affecting a broad number of proteins and pathways in the cell.

For a more comprehensive view of multivalent interactions it is reasonable to (1) analyze not only pairwise co-occurrence of different functional domains, but simultaneous co-occurrence of domains from several functional groups in proteins, (2) extend the analysis to complexes of chromatin proteins. The results of such analyses are presented in **Fig. 8D** (see **<u>Methods</u> Section 2.1.4** for our selection of 513 protein complexes (out of 2266) where all proteins are chromatin proteins from Complex Portal [47]). In our analysis at the level of individual proteins, proteins harbored domains only from up to four functional groups. Particularly, DNA binding domains may be combined with (di/oligo)-merization, chromatin remodeling, histone methylation, methylation and acetylation or histone acetylation and chromatin remodeling domains. The formation of chromatin protein complexes considerably enlarges the available combinations of functional domains. Among the 513 analyzed protein complexes, 181 complexes contained EMVI-domains from the analyzed functional domain groups, 101 complexes harboured more than one domain, 80 complexes harbored domains from different functional groups. From these 80 complexes the majority were various chromatin remodeling complexes (53 complexes), others representatives included histone acetyltransferase (13) and deacetylase (4) complexes, and DNA-methyltransferase complexes (2). One can see that the largest number of analyzed complexes (24) simultaneously contained domains from four functional groups (DNA binding, histone methylation, histone acetylation, and chromatin remodeling), in a select number of complexes domains belonging to up to six functional groups were observed (all the above mentioned together with domains involved in DNA methylation and histone binding). These were all complexes involved in chromatin remodeling. For example, ‘MBD2’ or ‘MBD3/NuRD nucleosome remodeling and deacetylase complex’. Notably, in chromatin complexes histone acetylation domains are found more often than histone methylation domains (unlike in the case when individual proteins are analyzed), this might, however, be biased by the current list of known chromatin complexes and their variability. Taken together chromatin protein complexes expand the multivalency of chromatin protein interactions and expand the functionality of the complexes.

As a final step we looked for chromatin proteins that matched to the most number of Pfam domain models that belonged to different functional groups/sub-groups and thus might engage in interactions of high multivalency (see **Suppl. Fig. SF8_2A**). According to this analysis, chromatin proteins could harbor domains from up to four functional groups/sub-groups. For example, histone-lysine N-methyltransferase ASH1L is involved in reading and writing of histone methylation, reading of histone acetylation and histone binding (**Fig. 8E**). Proteins could harbor up to nine Pfam domain models (some from identical functional categories). One of such examples is Histone-lysine N-methyltransferase NSD2 (**Fig. 8E**). This protein combines domains that likely engage in methylated histone binding (PWWP, PHD-1st_NSD, NDS_PHD, PHDvar_NSD, PHD, C5HCH), histone methylation (SET), histone binding (AWS) and may be DNA binding (HMG_box).

## 4. DISCUSSION

The presented study is a comprehensive effort to summarize, critically evaluate, systematize various types of information sources about chromatin proteins and use this meta-analysis to perform the state-of-the-art characterization of chromatin proteins’ composition and properties in order to describe the functioning of the chromatome as a holistic entity.

To our knowledge this is the first meta-analysis study aimed at identifying which human proteins are present in chromatin (the chromatome) where we cross-compared conceptually different sources of experimental (mass-spectrometry-based and microscopy based) and database derived information (including databases of protein localization and database of known classes of chromatin proteins). The results of our meta-analysis highlight challenges both in establishing the *per se* definition of the chromatome and using various approaches to collect the list of proteins according to the said definition. In fact, historically, chromatome was operationally defined either through the sets of proteins that were known to contribute to the structure and functioning of chromatin (which itself is an evolving term) [1] or through proteomics analysis of chromatin extracts (which depend on the experimental techniques) [17]. In our view, the chromatome—under a broad definition—may include all proteins with nuclear localization that can interact with the genomic DNA either directly or indirectly, with the possible exception of proteins tightly bound to the nuclear envelope. As we showed, within this definition, there is currently limited congruence between the sources of information and further efforts are needed to improve the quality and comprehensiveness of the datasets. Particularly, the MS-based studies suffer from an inherent limitation due to the fact that chromatin is a “fuzzy organelle” [23] (whose composition depends on the type of the cell line used and other conditions, given that many genes are conditionally expressed), supplemented by the technical problems in identifying the low abundant proteins and contamination during chromatin extraction. Localization studies have limited coverage of the proteome space, and may report only certain localizations for proteins, while according to our estimates ∼50% of nuclear proteins may be also localized in other cellular compartments.

The processes happening within the nuclei of eukaryotic cells are perhaps ones of the most complex that Nature has created. While a lot is already known about the molecular mechanisms of processes happening in chromatin, the holistic understanding of its functioning that would be able to explain how complex and tightly regulated organismal traits and behavior originates from these molecular processes remains a big challenge. While a lot of expectations were based on application of cybernetics [78], and now are based on big data and AI technologies, one of the main approaches to rational understanding is the reductionist analysis of the system. Hence, keeping in mind all the known limitations (such as the multiple functions performed by many proteins) in order to understand and to analyze the properties of the human chromatome we proposed the hierarchical SimChrom classification ontology of chromatin proteins based on a manual synthesis and analysis of the established body of knowledge. The main advantage of SimChrom is its relative simplicity which enabled us to analyze and compare the properties of different groups of chromatin proteins without being overwhelmed by the amount of details. Other classifications, such as GeneOntology [35,36] do not offer such an advantage to this end, other databases do not offer the comprehensiveness of SimChrom and focus on only specific protein classes [55,79,43]. The SimChrom classification ontology would not be useful without the actual classification of ∼3000 of chromatin proteins according to this ontology which form the SimChrom dataset. Another approximation/tradeoff useful for the rational analysis of the chromatome introduced in this study is the use of the SimChrom-SL classification, where each protein is ascribed only to one SimChrom category. Such an approach allowed us to highlight the differences in protein properties that may be essential for the particular protein class.

In our analysis of chromatin protein abundance we observed that MS-based studies of chromatin protein extracts yield very fluctuating abundance estimates (due to their analysis of specific cell lines and apparent bias to detect highly abundant proteins) and hence establishing the exact abundance profiles of chromatin proteins inside the cell nucleus is in our opinion an unsolved challenge. Various chromatin MS-based studies provided highly varying intensities for histones. Hence techniques that attempt to convert MS-intensities to protein copy numbers based on known histone mass in the cell (such as “proteomic ruler”) [80] are likely not applicable in the case of chromatin extracts studies. To analyze chromatin proteins’ abundance, we used the normalized data from whole organism proteomes collected by PaxDb, which allowed us to gain certain insights but could provide only aggregate abundance for proteins with multiple localizations. According to our analysis, the distribution of chromatin proteins with respect to their abundance is bimodal with approximately one half representing the highly abundant ones (> 1 ppm) and another half the low abundant ones (< 1 ppm). This is quite similar to that observed for the whole proteome. Chromatin proteins are moderately enriched by those performing housekeeping functions in comparison with the whole proteome (60% of chromatin proteins are housekeeping ones). This suggests that the presence of many low-abundant chromatin proteins in the whole organism proteome may be attributed not only to their mere conditional expression in certain cells, but at least in certain cases to their low levels of expression in the cells *per se* (assuming that housekeeping genes are expressed in all cells). A significant fraction of the chromatome (45 %) is represented by transcription factors. Proteomics and systems-biology studies of TF regulatory networks have shown that TFs can be expressed at very low levels and still play essential roles in cellular processes such as environmental responses, development, and differentiation [81,82]. Using our SimChrom classification and combining data on whole organism protein abundance and protein localization we were able to characterize the main chromatin proteins and chromatin protein groups that dominate by their mass fraction in the cell and the cell nucleus. Early studies established that histones are present in approximately the same mass as DNA, whereas non-histone chromosomal proteins contribute about 0.3-0.8 g per gram of DNA [1]. However, results can vary substantially depending on the experimental approach, methodology, and resolution [83]. To what extent these estimates are accurate within the definition of the chromatome used in this study still remains to be quantified. Our analysis is consistent with the fact that histones (with H4 histone being the sole leader due to the identity of its protein sequence among the family of encoding genes) dominate the protein contents of the cell nucleus (by the number of molecules and likely by the total mass), however, the proteins involved in RNA processing (including splicing factors, pre-mRNA binding proteins) rivals them once the mass fractions are compared (these proteins are on average considerably longer than histones). The latter fact highlights the functional importance of the nucleus as not only the DNA storing and processing entity, but also as a factory to produce and mature RNA molecules. This idea is also supported by the fact that among the most abundant individual proteins in the nucleus are histone chaperones nucleophosmin (NPM1) and nucleolin (NCL), which are also involved in nucleolar organization and ribosome biogenesis. Our analysis is consistent with earlier established knowledge that among the most abundant *bona fide* DNA/nucleosome interacting proteins are those from the HMG protein group. This is a diverse group of small, basic and abundant proteins that modulate chromatin architecture and dynamics through non-sequence-specific and dynamic interactions with DNA and nucleosomes [84].

Using our carefully prepared datasets and chromatin proteins classification we were able to systematically and quantitatively address the questions of chromatin proteins composition both at the amino acid and domain level. The importance of intrinsically disordered regions in chromatin proteins have recently been the focus of many studies, suggesting their properties are essential to the dynamic nature of chromatin functioning. IDRs enable proteins to interact transiently and multivalently with multiple partners [85,86], recruit them through short linear motifs (SLiMs) [87,88] and “fuzzy” interactions where proteins can adapt to multiple partners [89,90], facilitate DNA binding and DNA motif search and recognition [91,92], promote liquid-liquid phase separation and formation on non-membrane organelles enriched with certain proteins [6,93]. We showed that both the fraction and the number of IDRs in chromatin proteins is higher than in cytoplasmic ones, which is reflected in particular changes in their amino acids composition. Specifically, they are enriched in serine and proline and are depleted in hydrophobic aliphatic amino acids. It is generally assumed that chromatin proteins are positively charged to compensate for the negative charge of DNA and RNA molecules. We show that this is true in general, but certain important protein groups are negatively charged (histone chaperones, RNA polymerases, Histone PTM erasers), suggesting charge-charge interactions are an important factor in organizing the structure and functioning of chromatin. Moreover, we find that there are certain specific charge patterns along the protein sequence – tails of chromatin proteins (especially the N-tails) are negatively charged on average. This property may influence translation efficiency and protein expression, consistent with the general relationship between peptide charge and translation [94]. We showed that the increased positive charge of chromatin proteins is conferred through alterations in the occurrence of positive and negative amino acids which is different for IDRs and non-IDRs. Counterintuitively, certain positive amino acids may be depleted in certain regions (*e.g.*, arginine is depleted in IDRs of chromatin proteins), suggesting there may be other factors and physico-chemical properties affecting the presence of charged amino acids along the protein sequence. Arginine residues exhibit a strong affinity for DNA due to their positively charged guanidinium groups, facilitating electrostatic interactions with the negatively charged DNA backbone. This property is advantageous in structured DNA-binding domains but may be detrimental in intrinsically disordered regions. Arginine residues strongly interact with DNA and RNA via their positively charged guanidinium groups, stabilizing structured nucleic acid–binding domains. However, in IDRs, excessive arginine can cause nonspecific nucleic acid binding, restricting flexibility and disrupting LLPS [95–97]. This might be an explanation for the relative depletion of arginine in IDRs of chromatin-associated proteins, balancing nucleic acid interactions with the need for dynamic, phase-separating behavior. Our analysis of domain annotation of chromatin proteins suggests that while non-IDRs are relatively well annotated (∼60% are annotated by Pfam), the IDRs are poorly annotated (∼87% are unannotated) – highlighting the challenges in understanding their functions and interactions. While IDRs resist direct structural characterization with conventional structural biology methods, we envision that the application of molecular modeling in combination with the state-of-the-art machine learning models that can link protein sequence, dynamics, and function is a perspective way to move forward in deciphering the role of IDRs in chromatin functioning [98–101].

The structural characterization of chromatin protein domains is also an important task that is not completely solved. According to AlphaFold ∼50% of the aggregate chromatin protein sequence are non-IDRs, while 35% form *bona fide* structural domains in monomeric proteins (as predicted by AphaFold and TED). Among these domains around 70% have relatively close homologs (with sequence identity of more than 50%) available in PDB. The advances in AI-based protein structure prediction currently open exciting opportunities to bridge this gap in the structural characterization of chromatin proteins. Particularly, using such tools we identified 241 structural domains in chromatin proteins whose 3D structures (or structure of their detectable homologs) were not previously experimentally solved, 106 proteins have domains that are also not functionally annotated by Pfam. These domains await experimental structural and functional characterization. We envision that the development and application of AI-based structure prediction tools will facilitate chromatin research, particular in analysis of protein-protein interactions, which is already happening [102,103].

The importance of multivalent protein interactions in chromatin was highlighted by D. Allis and colleagues around 20 years ago [104]. Here we performed systematic analysis of domains engaged in multivalent interactions present in chromatin regulator proteins (histone PTM readers/writers/erasers, chromatin remodelers, histone chaperones, proteins involved in DNA or nuclear RNA modifications). We showed that chromatin regulators indeed actively use such domains and may contain domains corresponding to up to nine distinct Pfam domain models. In chromatin protein complexes the number of such domains is significantly increased. Chromatin regulators contain an especially large number of domains involved in histone methylation that may be combined with histone acetylation, histone binding, DNA binding and DNA methylation domains. The data collected in this study provide further framework for understanding the engagement of multivalent interaction in chromatin functioning.

The hallmark feature of this work is the SimChrom web-resource which allows the user to explore the results of our analysis interactively, query and compare the information about the proteins the user is interested in from the curated datasets. This is a unique tool that uses interactive web-based data representation to address the multidimensionality and heterogeneity of the available data about chromatin proteins and allows to extract knowledge from the data. Inspired by the best practices of collaborative research in data science, SimChrom is implemented as a GitHub repository that is rendered as a web-site directly from GitHub, making it a secure, reliable, open source tool that may be easily copied and modified by the community.

Taken together we hope that our work establishes a holistic framework for further advances in the field of chromatin research which will help to understand genome functioning though deeper appreciation of the complex role played by the chromatome.

## ACKNOWLEDGEMENTS

We thank A.L. Sivkina, D.K. Malinina, N.S. Gerasimova, A.V. Lyubitelev, S.V. Ulianov, and A.A. Gavrilov for valuable discussions that helped to improve this work.

## AUTHOR CONTRIBUTIONS

AKG: Conceptualization, Data curation, Formal analysis, Investigation, Methodology, Software, Visualization. GAA: Resources, Software. MPK: Conceptualization. Writing – review & editing. AKS: Conceptualization, Formal analysis, Funding acquisition, Methodology, Supervision, Validation, Writing – original draft, Writing – review & editing.

## SUPPLEMENTARY DATA

Supplementary material is available online, including supplementary figures, tables, supplementary results and discussion.

## DATA AVAILABILITY

The SimChrom database including interactive supplementary materials about chromatin proteins’ classification, localization, functions, domain composition are freely available at a GitHub hosted web site https://simchrom.intbio.org/. The SimChrom source code is available at GitHub https://github.com/intbio/SimChrom and archived via Zenodo at https://doi.org/10.5281/zenodo.17314850.

