## Supplementary Information for "(Re)defining the human chromatome: an integrated meta-analysis of localization, function, abundance, physical properties and domain composition of chromatin proteins"

#### Supplementary Tables

**Supplementary Table ST1.** Representative list of chromatome datasets from whole proteome projects, databases, literature sources and computational predictions about chromatin or nuclear proteins. A short description and the number of proteins is provided for every source. Columns: Source of information; DB, link, data availability; Description of source of information, details of experiment; Number of proteins (machine readable), Year; Reference; Doi; Notes about number of proteins (human readable); Number of processed proteins.

**Supplementary Table ST2.** The evaluation metrics for all localized, nuclear and subnuclear (nucleolus, nuclear envelope, nuclear bodies, nucleoplasm) proteins from UniProt (A) and HPA (B). Metrics include the number of proteins with overlapping annotation, union of annotations, Jaccard similarity coefficient (measuring overlap), False Positives, False Negatives, and performance measures (precision, recall, and F1-score).

**Supplementary Table ST3.** Localization terms from UniProt and HPA which were grouped into generalized localization categories: nuclear categories (nucleus, nucleolus, nuclear envelope, nuclear bodies, nucleoplasm), cytoplasm, endomembrane system and others (in UniProt).

**Supplementary Table ST4.** SimChrom classification terms and details about their protein contents: corresponding GO terms, databases and literature sources, additional filters. Columns: SimChrom term, GO terms used to build the SimChrom DB, Explanation of the selected GO terms to build SimChrom DB, Number of proteins initially extracted from GO without proteins in nested classes (before additional filtration), Databases and literature sources used to add proteins and explanation, Additional filtering applied, Total number of proteins (with proteins in nested classes), Additional notes.

**Supplementary Table ST5.** The SimChrom ontology and SimChrom/SimChrom-SL classification of chromatin proteins. The list of SimChrom protein categories is given together with the aspects that they represent, and the lists of proteins belonging to each category given via UniProt IDs. Total number of proteins per category is also given.

**Supplementary Table ST6.** The constructed protein localization reference datasets, their definition and the corresponding number of protein entries. Abbreviations: NULOC - nuclear localization, NON_NULOC - non-nuclear localization, CYTLOC - cytoplasmic localization (see definition in **Supplementary Table ST3**), UL - uniquely localized, CS - consensus (supported by several sources), JT - joint (supported by at least one source), NECF - no evidence code filtering (localization data with any evidence level from the databases was used).

**Supplementary Table ST7.** GO enrichment analysis for the set of protein entries from MS-based chromatomes that are absent in NULOC_JT_NECF and SimChrom (n=2232). Only the driver GO terms that were highlighted by g:Profiler were included. Columns: source, term_name, term_id, highlighted, adjusted_p_value, negative_log10_of_adjusted_p_value, term_size, query_size, intersection_size, effective_domain_size, intersections.

**Supplementary Table ST8.** GO enrichment analysis of SimChrom-exclusive proteins (absent in NULOC_CS), n=1208. Only the driver GO terms that were highlighted by g:Profiler were included. Columns: source, term_name, term_id, highlighted, adjusted_p_value, negative_log10_of_adjusted_p_value, term_size, query_size, intersection_size, effective_domain_size, intersections, unexpected_for_chromatin.

**Supplementary Table ST9.** GO enrichment analysis of NULOC_CS-exclusive proteins (absent in SimChrom), n=1459. Only the driver GO terms that were highlighted by g:Profiler were included. Columns: source, term_name, term_id, highlighted, adjusted_p_value, negative_log10_of_adjusted_p_value, term_size, query_size, intersection_size, effective_domain_size, intersections.

**Supplementary Table ST10.** The number and fraction of housekeeping (HK)/ non-housekeeping (non-HK), low-abundant (LA) and high-abundant (HA) proteins in the whole proteome (PaxDB_INT), NULOC_JT, NULOC_CS, SimChrom and MS-based chromatomes and a nucleome.

**Supplementary Table ST11.** A list of SimChrom-SL proteins annotated with "RNA processing" category that are uniquely localized in the nucleus (NULOC_CS_UL). Columns: Gene, Entry, Protein names, localization by UniProt, HPA, OpenCell with filtration by evidence codes (see Methods), protein identification in MS-based studies, abundance by PaxDB_PA (ppm).

**Supplementary Table ST12.** Abundance data for all histone proteins and non-histone chromatin proteins with abundance values of more than 0.01% of Histone H4. Columns: SimChrom-SL, Gene, Abundance relative to histone H4 (%), Number of nucleosomes per protein, Abundance PaxDb_PA (ppm), Entry, Uniquely localized in nucleus, Details (for Figure 4D).

**Supplementary Table ST13.** Comparison of amino acid composition between chromatin and Cytoplasm$^{UL}$ proteins. Table includes columns: Feature (fraction of amino acid), Sequence (total, IDR, non-IDR), median fraction for chromatin and cytoplasmic proteins in percent, fold enrichment of median values for SimChrom vs Cytoplasm$^{UL}$, statistically significance according to Mann Uitney test with multiple test correction (TRUE, FALSE), fold enrichment in IDRs vs non-IDRs of SimChrom, Fold enrichment in IDRs vs non-IDRs of Cytoplasm.

**Supplementary Table ST14.** List of uncharacterized structural domains in chromatin proteins. Columns: Annotated in InterPro (YES/NO), Entry, TED sequence number, ted_range, ted_len, pLDDT, packing_density, norm_rg, consensus_level, num_segments, num_helix_strand_turn, num_helix_strand, num_helix, num_strand, num_turn, TED_link, InterPro_link, SimChrom-SL, ted_id. See also **Interactive Table 3** (https://simchrom.intbio.org/#novel_structural_domains).

**Supplementary Table ST15.** List of Pfam domain models in chromatin proteins. Columns: Entry, SimChrom-SL, Total number of Pfam domains, Number of distinct Pfam models, pfamA_ids (only unique ids), Domain annotations (according to manual annotation of EMVI-domains, see Methods), Distinct domain annotations (only unique annotations).

**Supplementary Table ST16.** Pfam domain models that are present in three or more chromatin/epigenetic regulator proteins. Columns: pfamA_id, pfamA_acc, pfamA_name, domain model functional subgroup, domain model functional group, SimChrom-SL ChromReg (the category which contains a large fraction of proteins with the domain model, where available), number of SimChrom proteins with the domain model, number of ChromReg proteins with the domain model, number of co-occurring Pfam domain models in SimChrom proteins, number of co-occurring Pfam domain models in ChromReg proteins, number of SimChrom proteins (where the Pfam domain models co-occurs with other domain models), number of ChromReg proteins (where the Pfam domain model co-occurs with other domain models).

**Supplementary Table ST17.** The conditional probability of EMVI-domains to co-occur in chromatin proteins, grouped by their main functions. The conditional probability of finding a corresponding domain A in a chromatin protein given that another domain B is already present (columns and rows correspond to domains A and B, respectively).

#### Supplementary Figures

##### 1. Sources of information about chromatin and nuclear proteins and their critical evaluation

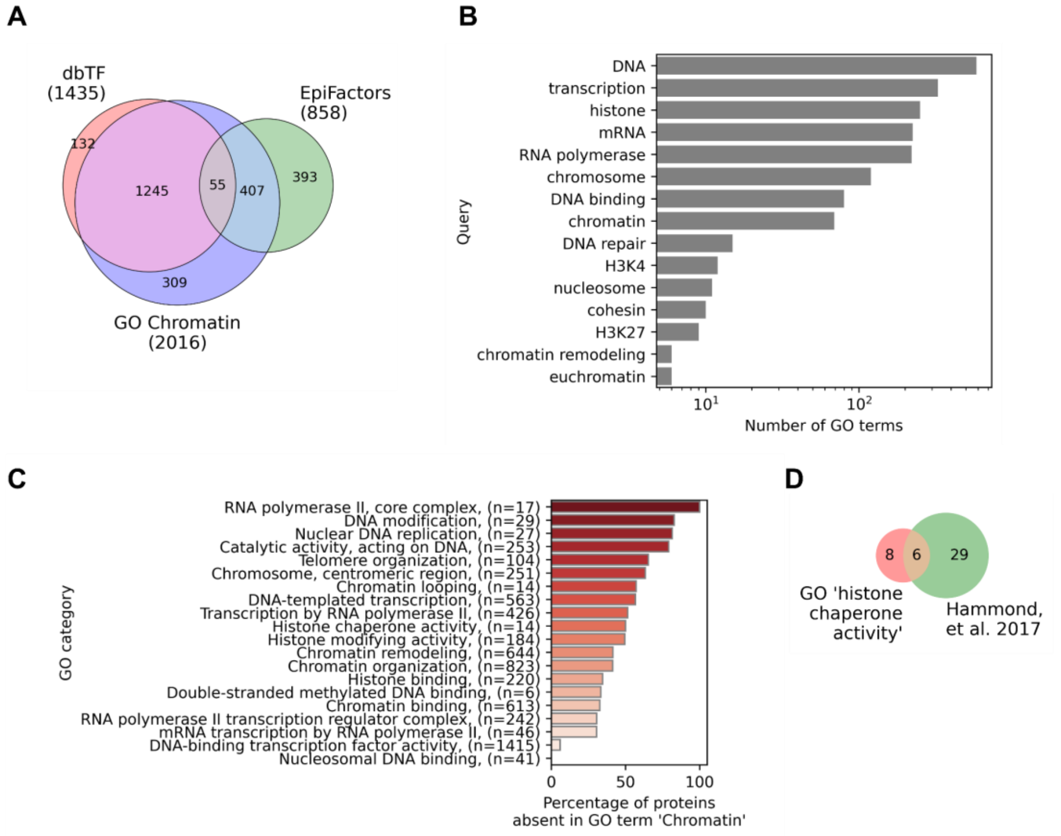

**Supplementary Figure SF2_1.** (**A**) The Venn diagram of protein repertoires from EpiFactors (epigenetic regulators), the GO catalogue of DNA-binding transcription factors, and GO term 'Chromatin'. (**B**) The number of GO terms which contain chromatin-associated keywords. (**C**) The fraction of proteins from suggested chromatin-associated categories and proteins annotated by the GO term 'Chromatin'. (**D**) The Venn diagram for histone chaperone proteins: proteins annotated by the GO term "histone chaperone activity" vs literature review by Hammond et al., 2017 [1].

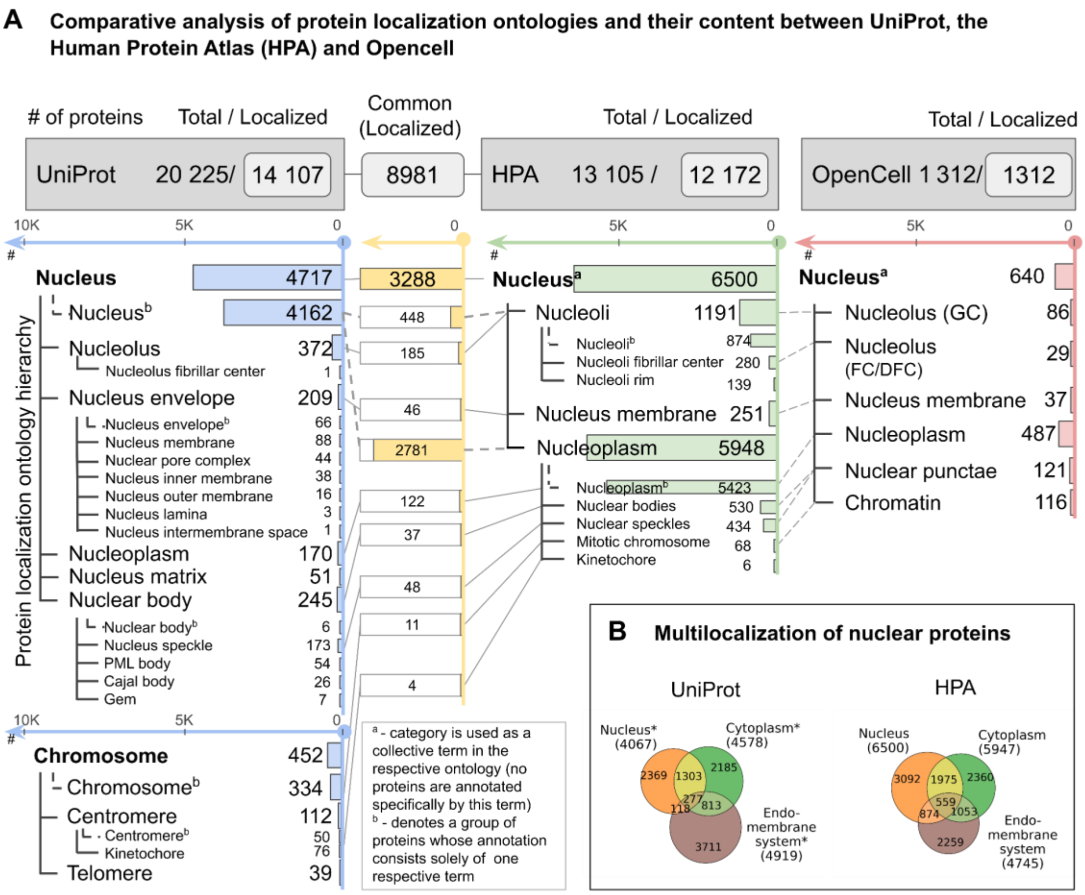

**Supplementary Figure SF2_2. Nuclear proteome content and protein subnuclear localization according to three major databases/proteome-wide experimental initiatives: UniProt, HPA, OpenCell.** See interactive version in **Interactive Figure 2** at https://simchrom.intbio.org/#localization. (**A**) A detailed comparative analysis of human nuclear proteome content and its subnuclear localization according to respective hierarchical ontologies between UniProt (blue), the Human Protein Atlas (HPA) (green) and Opencell (red). The scheme shows there ontologies for each localization annotation system, the number of proteins in each category (to the right of each category name), and the number of common proteins between different categories of UniProt and HPA (highlighted in yellow). Note that a protein may be present in several localization categories if it has multiple sublocalizations. On the top of the scheme the total number of human proteins in the database (total) and the number of proteins that have localization information supported by sufficient evidence (localized) is reported (see Methods) for the definition of the criteria). (**B**) A Venn diagram showing the overlap between the sets of nuclear, cytoplasmic and endomembrane proteins as defined by UniProt or HPA. UniProt proteins were considered with only three grouped localization tags; proteins with others localization were not considered.

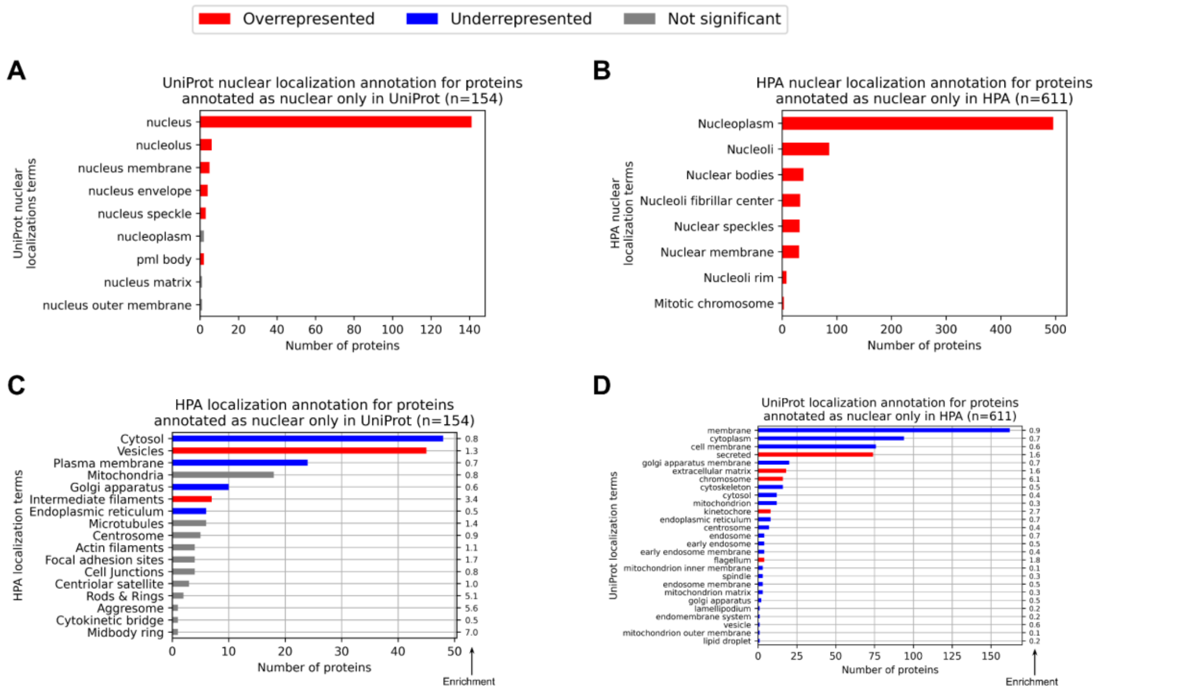

**Supplementary Figure SF2_3. A cross-annotation analysis of localization terms for proteins annotated as nuclear by one resource - either UniProt or HPA, while having only non-nuclear localization annotations in the other resource.** To identify significantly over- or under-represented localization terms, an enrichment analysis using the hypergeometric test (p-value < 0.05) was performed, see Methods Section 2.1.1. (**A**) The number of nuclear proteins in the UniProt with their subnuclear localization terms that lack nuclear localization annotation in HPA. (**B**) The number of nuclear proteins in the HPA with their subnuclear localization terms that lack nuclear localization annotation in UniProt. (**C**) The number of nuclear proteins in the UniProt that are non-nuclear in HPA, along with their non-nuclear localization annotation in HPA. (**D**) The number of nuclear proteins in the HPA that are non-nuclear in UniProt, along with their non-nuclear localization annotation in UniProt.

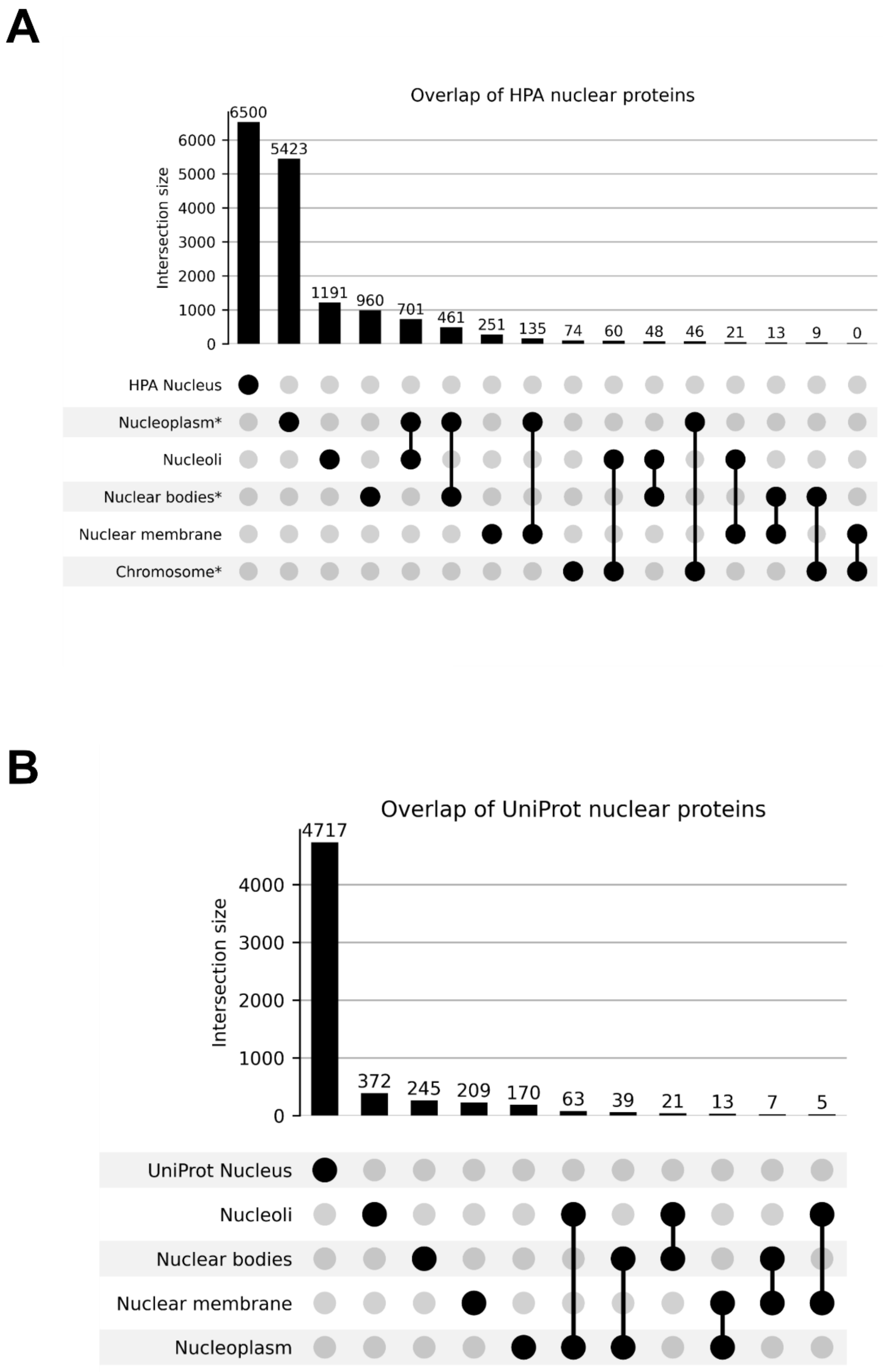

**Supplementary Figure SF2_4**. UpSet plots comparing sets of proteins according to their subnuclear localization as provided by HPA (**A**) and (**B**) UniProt. The generalized subnuclear localization terms are used as defined in Supplementary Table ST3.

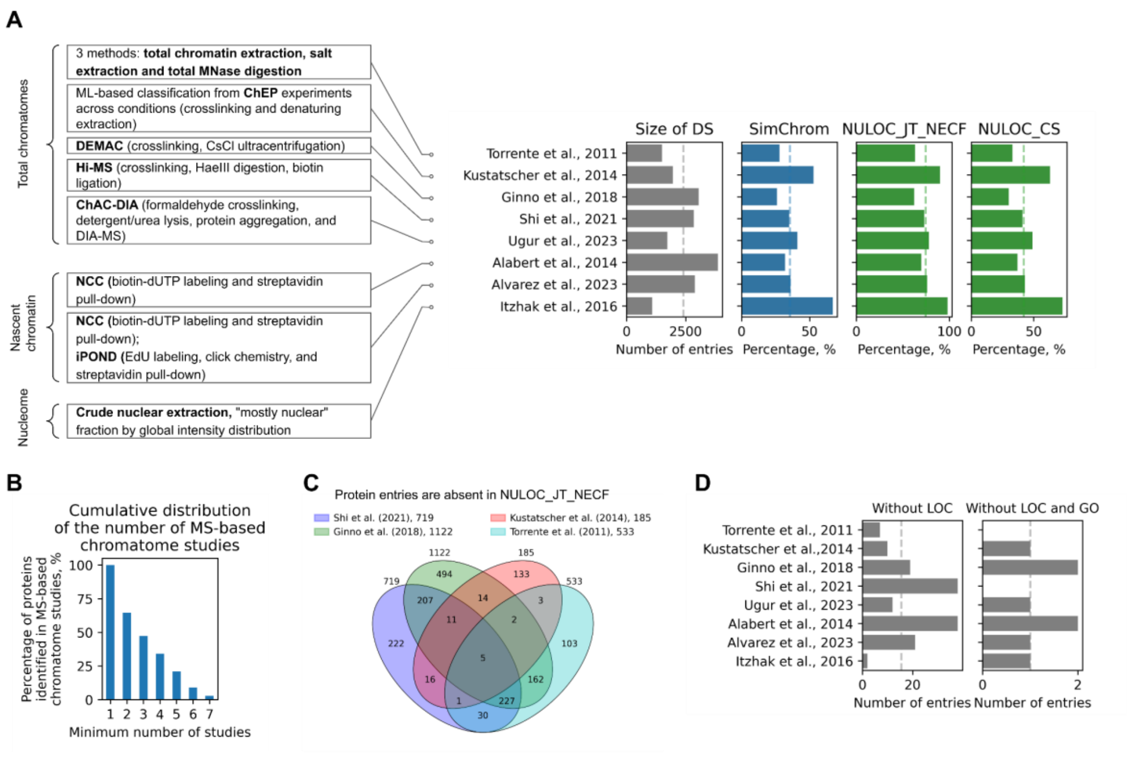

**Supplementary Figure SF2_5. The MS-based human chromatome (and nucleome) datasets examined through the lens of annotations provided by the localization databases and the SimChrom chromatin protein classification**. (A) The list of MS-based datasets of chromatin/nuclear proteins from the respective studies analyzed in this work together with a short description of the experimental/analysis workflow (left) and the plots (right) showing the size of the datasets and the fractions of the datasets that overlap with the SimChrom dataset or nuclear localization datasets (NULOC_CS and NULOC_JT_NECF). The median values are shown by dotted in the plots. (**B**) Cumulative fraction of proteins identified by at least N MS-based chromatin studies relative to the total number of chromatin proteins identified in at least one MS-based study. (**C**) Overlap of protein entries identified in MS-based studies that lack nuclear localization according to the database annotations (NULOC_JT_NECF dataset). (**D**) Number of protein entries identified in MS-based studies that have no annotations in the databases. Left: proteins lacking localization data in both UniProt and HPA. Right: proteins lacking both localization data and GO annotation.

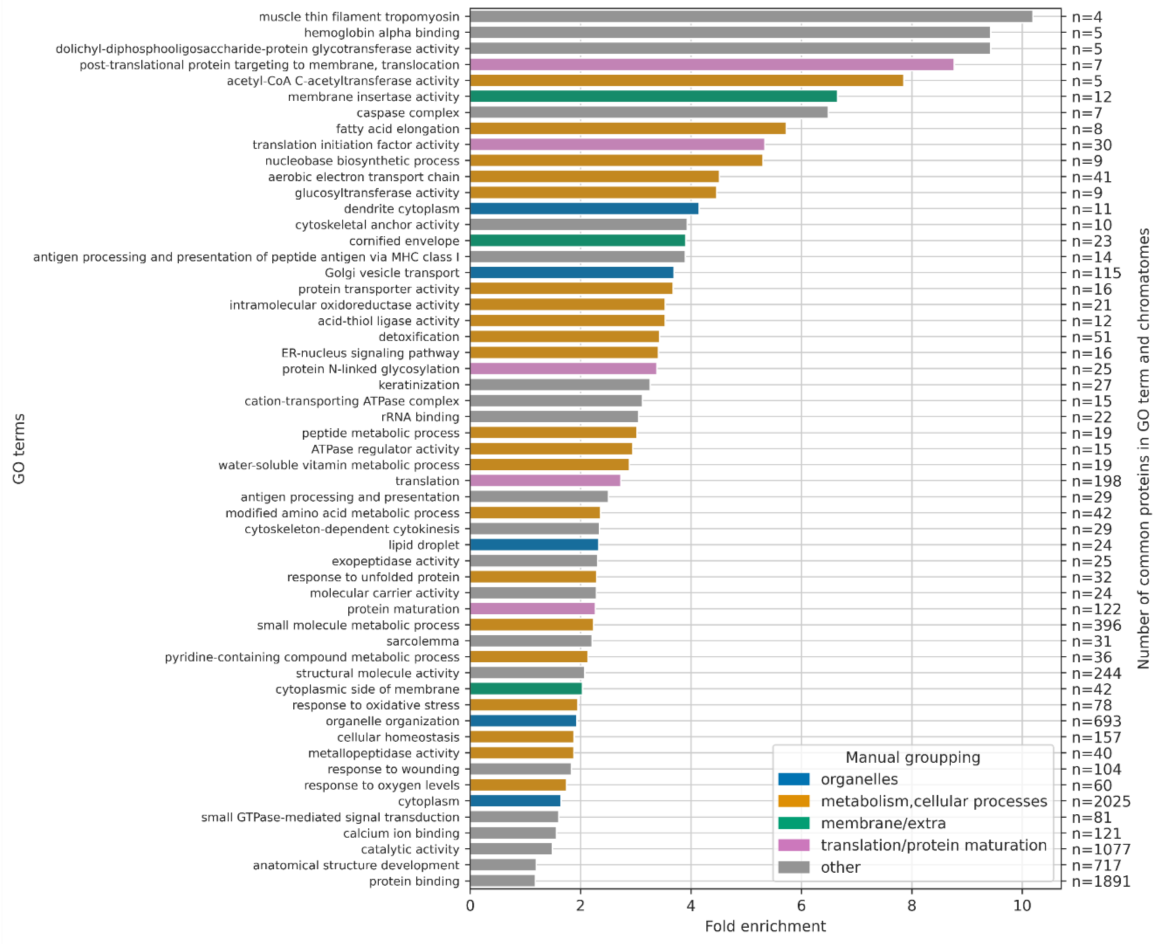

**Supplementary Figure SF2_6.** GO enrichment analysis for protein entries from MS-based chromatome datasets that are absent from both the NULOC_JT_NECF dataset and the SimChrom classification (n = 2232).

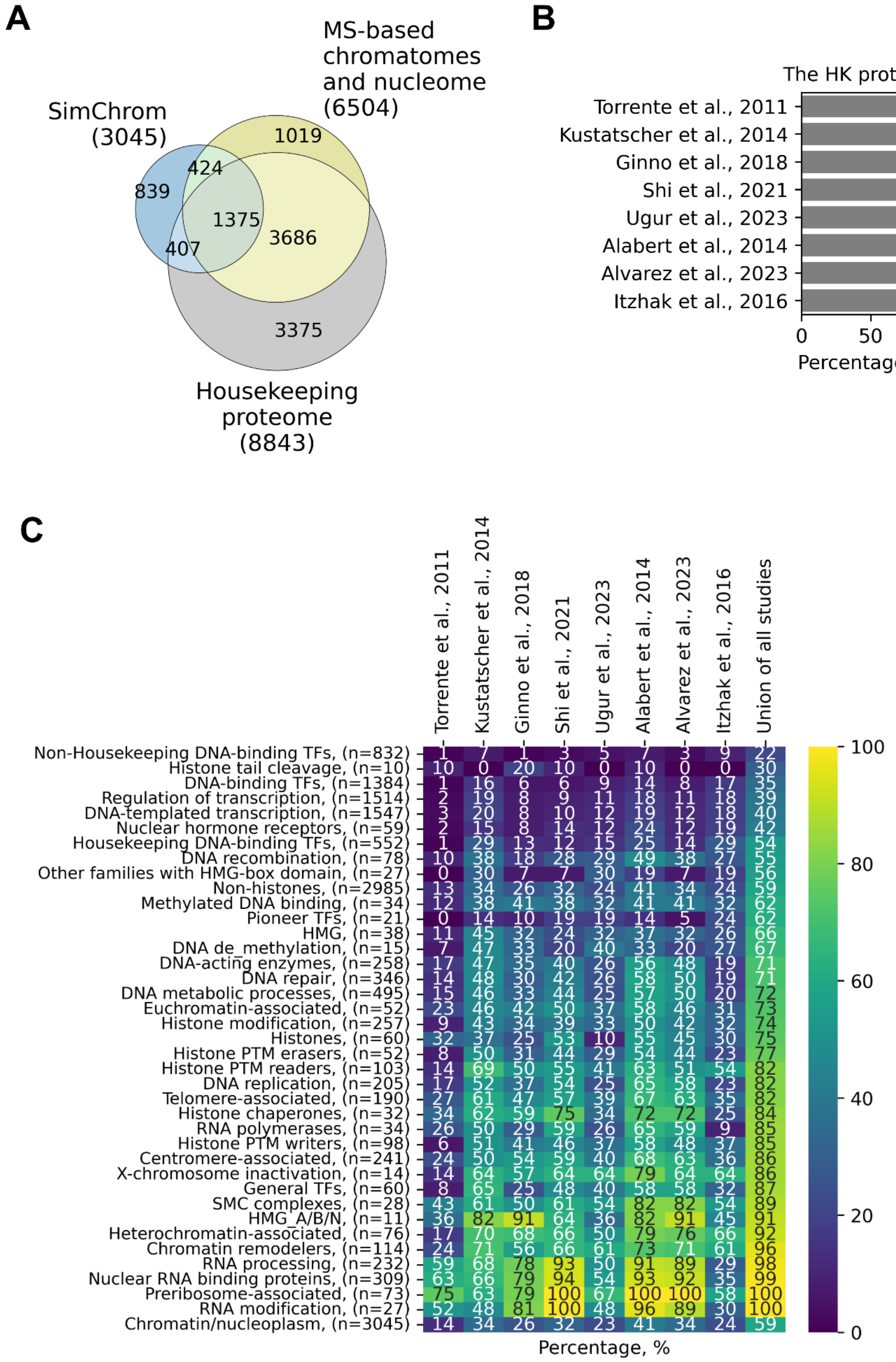

**Supplementary Figure SF2_7.** (**A**) The Venn diagram showing overlaps between the set of SimChrom proteins, the housekeeping proteome and the combined protein set of MS-based chromatome datasets (union of protein content from eight MS-based studies analyzed in this work). (**B**) The percentage of housekeeping (HK) proteins in MS-based chromatomes and nucleome (range: 76% - 90%). (**C**) The percentage of SimChrom proteins by SimChrom category in MS-based chromatomes and nucleome.

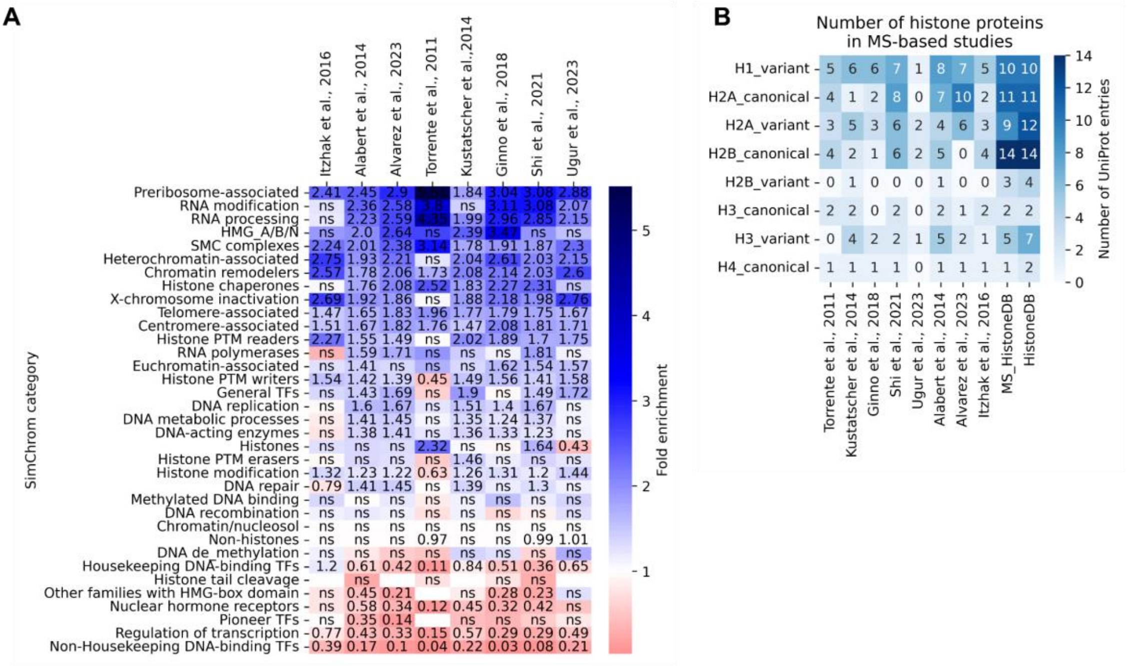

**Supplementary Figure SF2_8.** (**A**) Fold enrichment of chromatin-associated proteins identified in MS-based studies for every SimChrom category (fold enrichment is calculated with respect to the distribution of proteins among the categories in SimChrom). Only statistically significant values (p-value of Fisher exact test with Benjamini correction < 0.05) are shown. (**B**) Number of histone proteins detected in MS-based studies compared to the reference counts from MS_HistoneDB and HistoneDB 2.0.

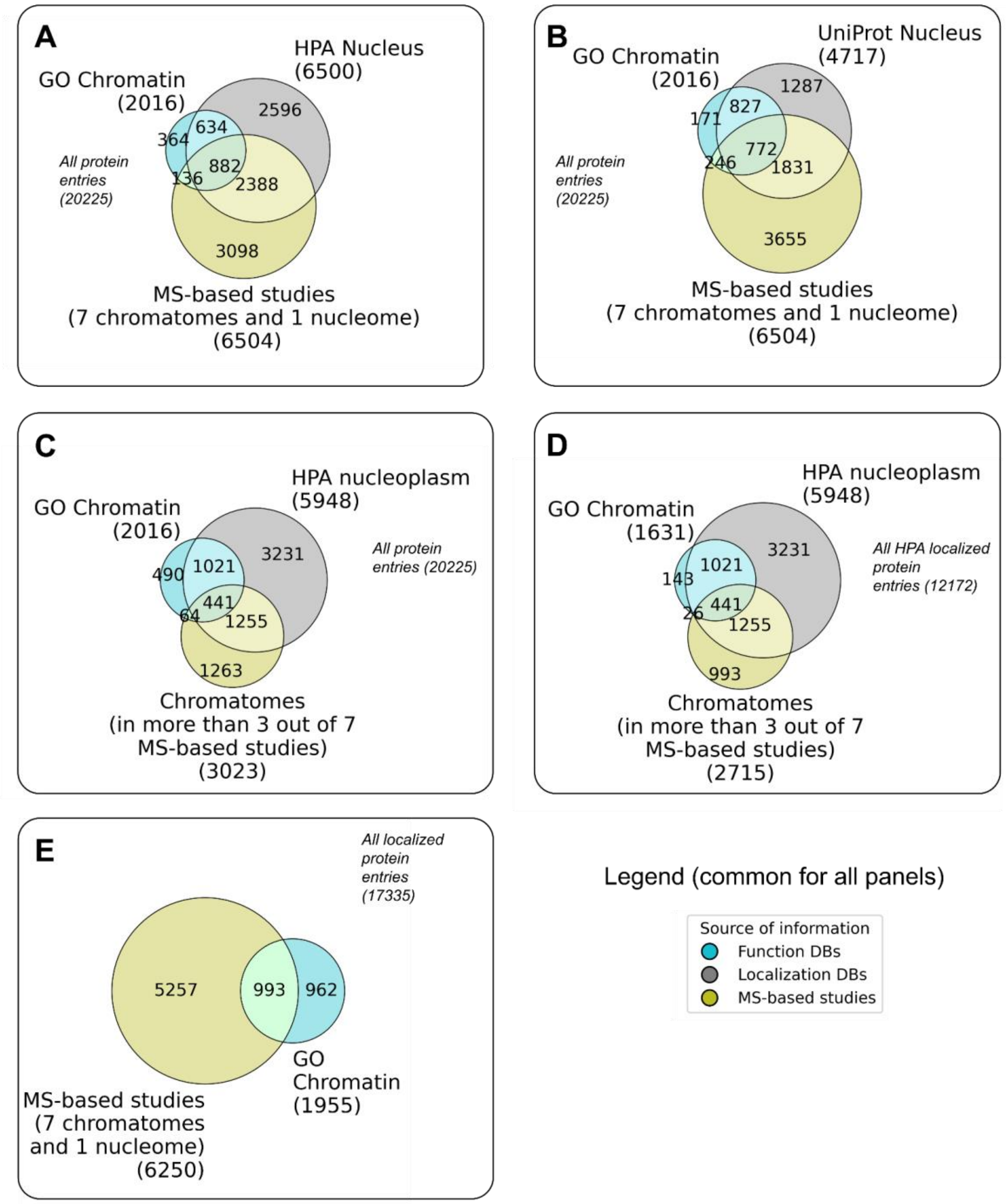

**Supplementary Figure SF2_9.** The overlap of chromatin/nuclear protein entries from different types of sources: protein function databases (GO "Chromatin"), protein localization DBs (Uniprot Nucleus, HPA Nucleus or Nucleoplasm), MS-based studies of chromatome and nucleome proteins (two protein sets are used - see legends: 1. union of protein entries from five total chromatin studies, two nascent chromatin and one nucleome; 2. protein entries that are present in three out of seven MS-based chromatin datasets). The "background" set of proteins for each panel is shown in italic.

#### 2. The SimChrom chromatin protein classification, the SimChrom dataset and other reference datasets

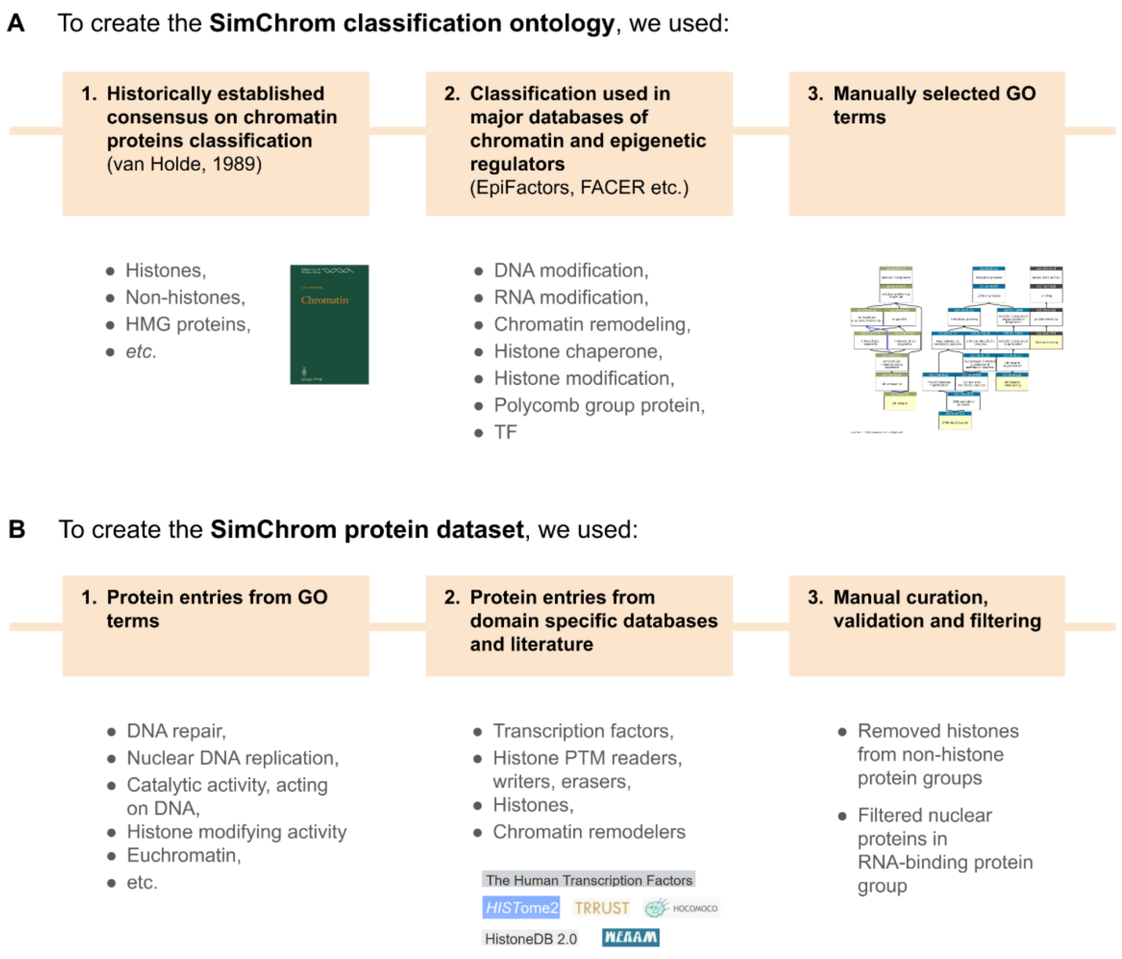

**Supplementary Figure SF3_1.** The scheme of creation of the SimChrom classification ontology and SimChrom protein dataset is shown in panels (**A**) and (**B**), respectively.

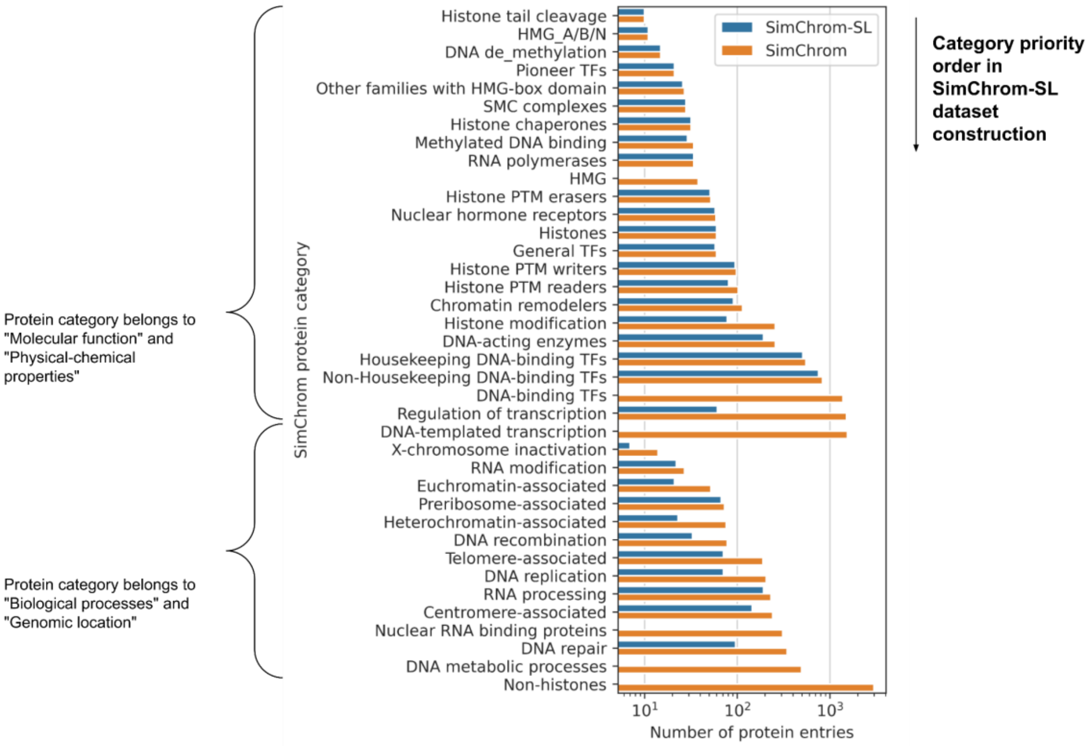

**Supplementary Figure SF3_2.** The order of SimChrom categories (from top to bottom) used to create the single-label SimChrom-SL classification of chromatin proteins. The categories were ordered as follows: molecular function and physicochemical properties were placed first, followed by the others. Among them, categories containing fewer proteins were ordered earlier. The number of proteins belonging to the respective SimChrom and SimChrom-SL categories is also shown.

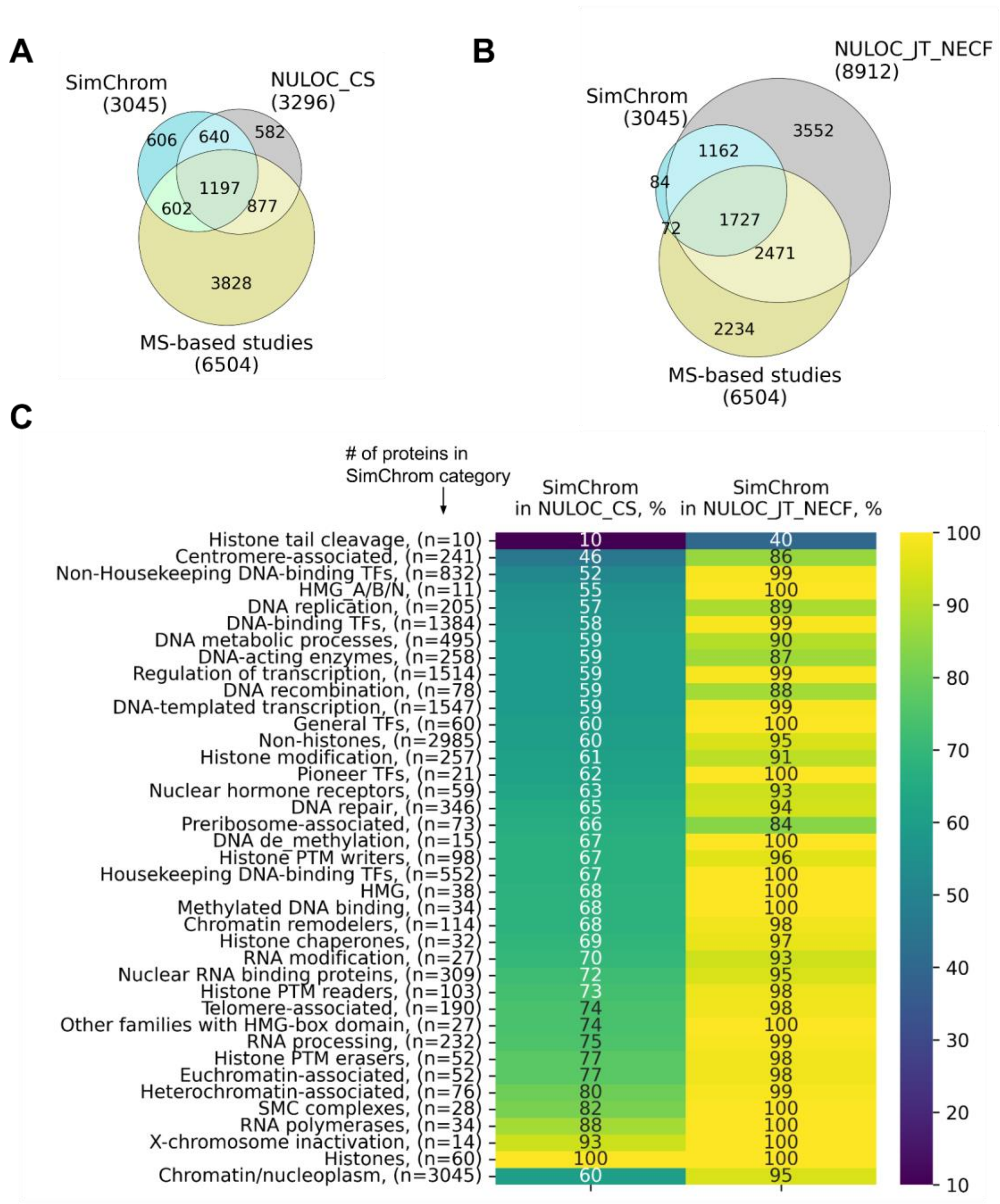

**Supplementary Figure SF3_3.** SimChrom protein analysis using protein localization information and MS-based chromatomes and nucleome. **(A-B)** A Venn diagram showing the overlap between the protein sets: SimChrom, proteins identified in MS-based studies and the reference nuclear protein sets NULOC_CS (**A**) or NULOC_JT_NECF (**B**). (**C**) The percentage of proteins from SimChrom categories that are found in NULOC_CS and NULOC_JT_NECF datasets.

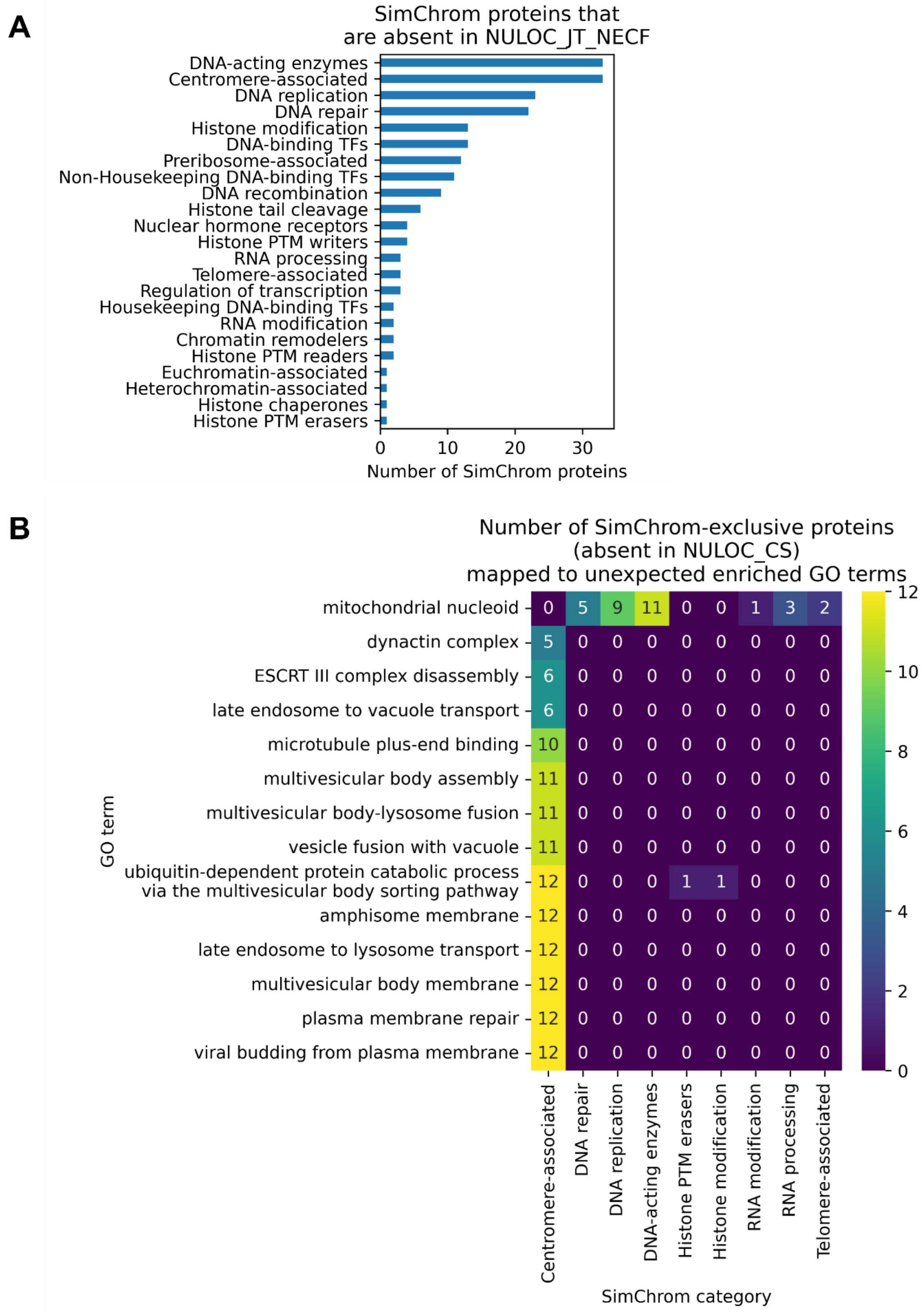

**Supplementary Figure SF3_4.** SimChrom protein dataset analysis using protein localization information. (**A**) The number of SimChrom-SL classified proteins without nuclear localization according to NULOC_JT_NECF (the broadest dataset that combined all nuclear protein entries from all protein localization databases at any level of confidence). (**B**) The unexpected enriched GO terms for proteins were identified for the SimChrom proteins that are absent in NULOC_CS.

#### 3. Analysis of the human chromatome

##### 3.1. The chromatome composition and abundance of chromatin proteins

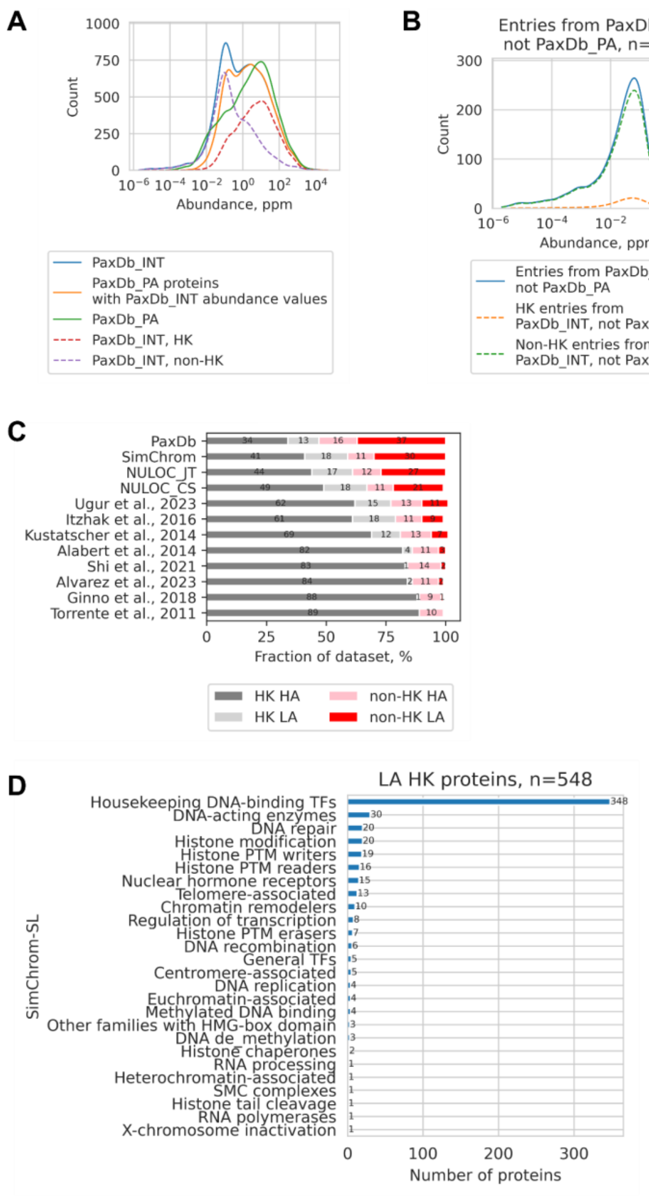

**Supplementary Figure SF4_1.** Chromatin proteins abundance analysis. (**A, B**) Distribution of proteins from PaxDb_INT and PaxDb_PA datasets according to their relative abundance values. The distribution for housekeeping (HK) and non-housekeeping (non-HK) are also shown (see legend). The distribution was constructed by taking the logarithm of the abundance values in ppm, making a histogram (bin size of 0.15) and smoothing it with a gaussian kernel for visual clarity. (**C**) Fraction distribution of low-abundant (LA) and high-abundant (HA) housekeeping (HK) and non-housekeeping (non-HK) proteins in the whole proteome (PaxDb_INT), protein localization datasets (NULOC_CS and NULOC_JT), and SimChrom and MS-based chromatomes. (**D**) The distribution of low-abundant (LA) housekeeping (HK) proteins among SimChrom-SL categories.

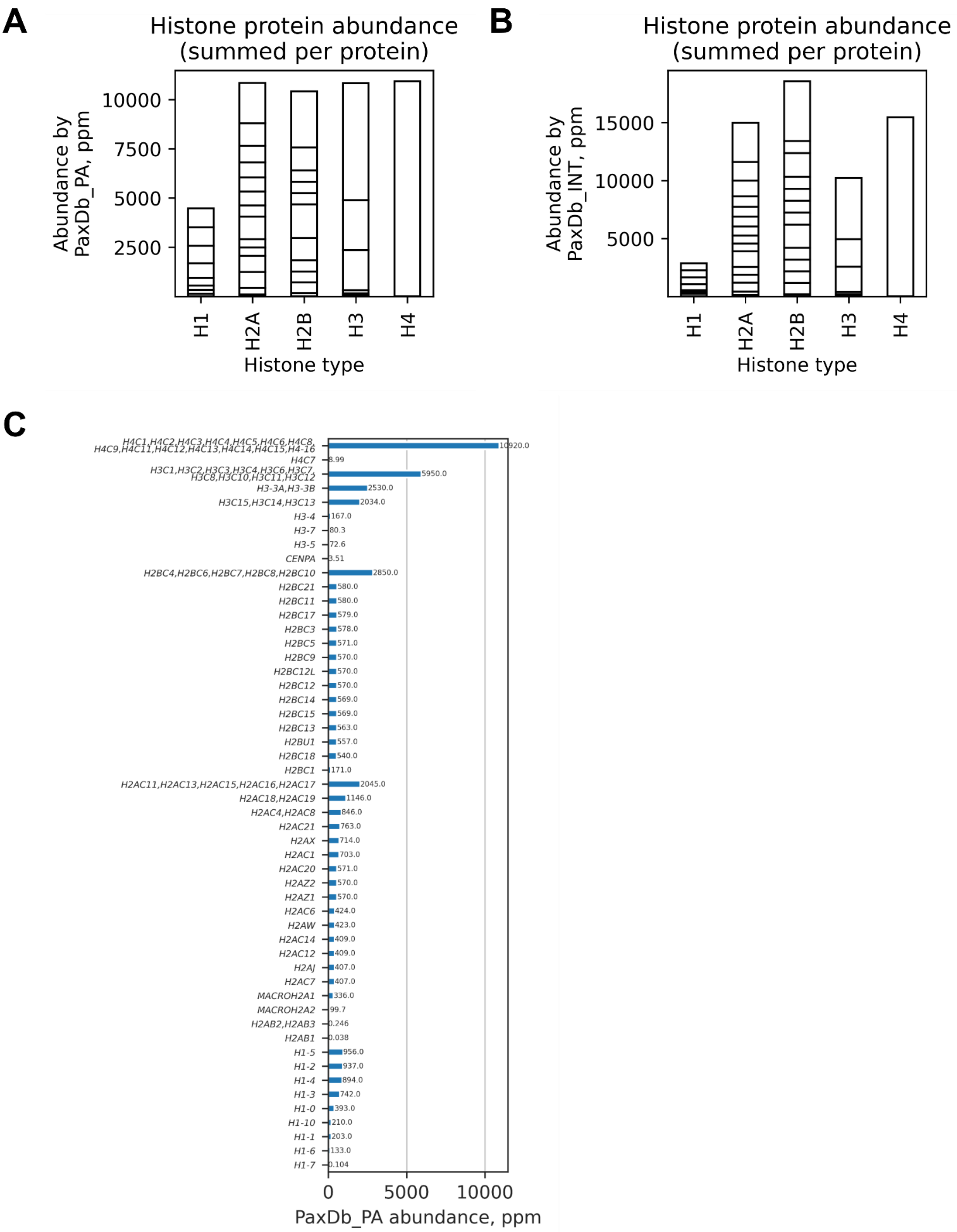

**Supplementary Figure SF4_2.** Abundance of histone proteins according to PAXDb datasets: PAXDb_PA "Whole organism, SC (Peptideatlas,aug,2014)" (**A**) and PAXDb_INT "H.sapiens - Whole organism (Integrated)" (**B**); ppm means part per million. Abundance of histone proteins grouped by histone types (each histone protein type is composed of one or several proteins that may in turn be encoded by several different genes) by PAXDb_INT. The horizontal lines on the bar plot delineate the contribution of different histone proteins belonging to the respective histone type. (**C**) Abundance of histone proteins according to PaxDB_PA.

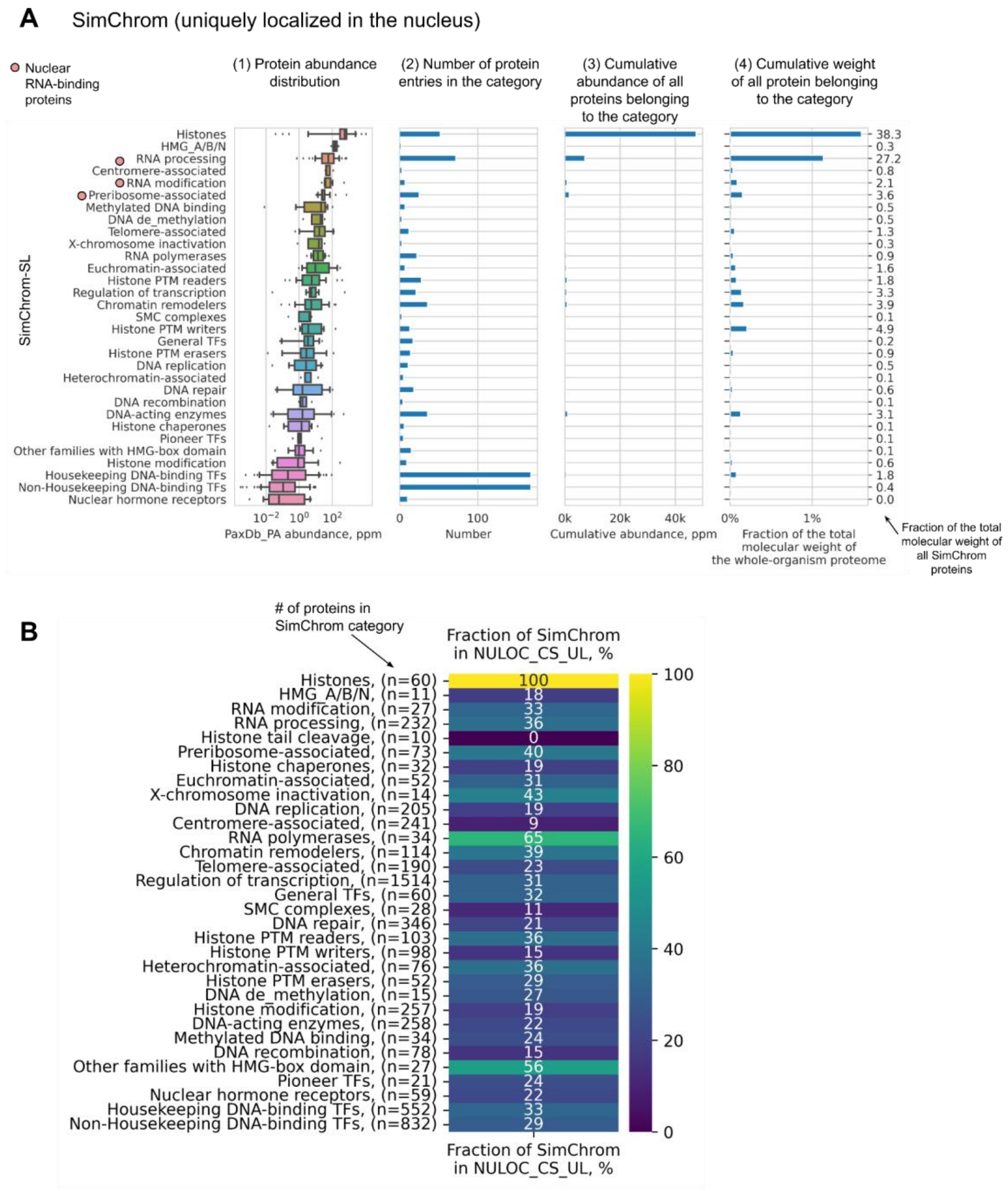

**Supplementary Figure SF4_3.** (**A**) This is a version of Figure 4C from the main text showing chromatin protein abundance for SimChrom categories, but created for the SimChrom proteins that are uniquely localized in the nucleus according to the NULOC_CS_UL dataset. (**B**) The fractions of proteins belonging to SimChrom categories (based on standard SimChrom classification) that are also present in NULOC_CS_UL dataset.

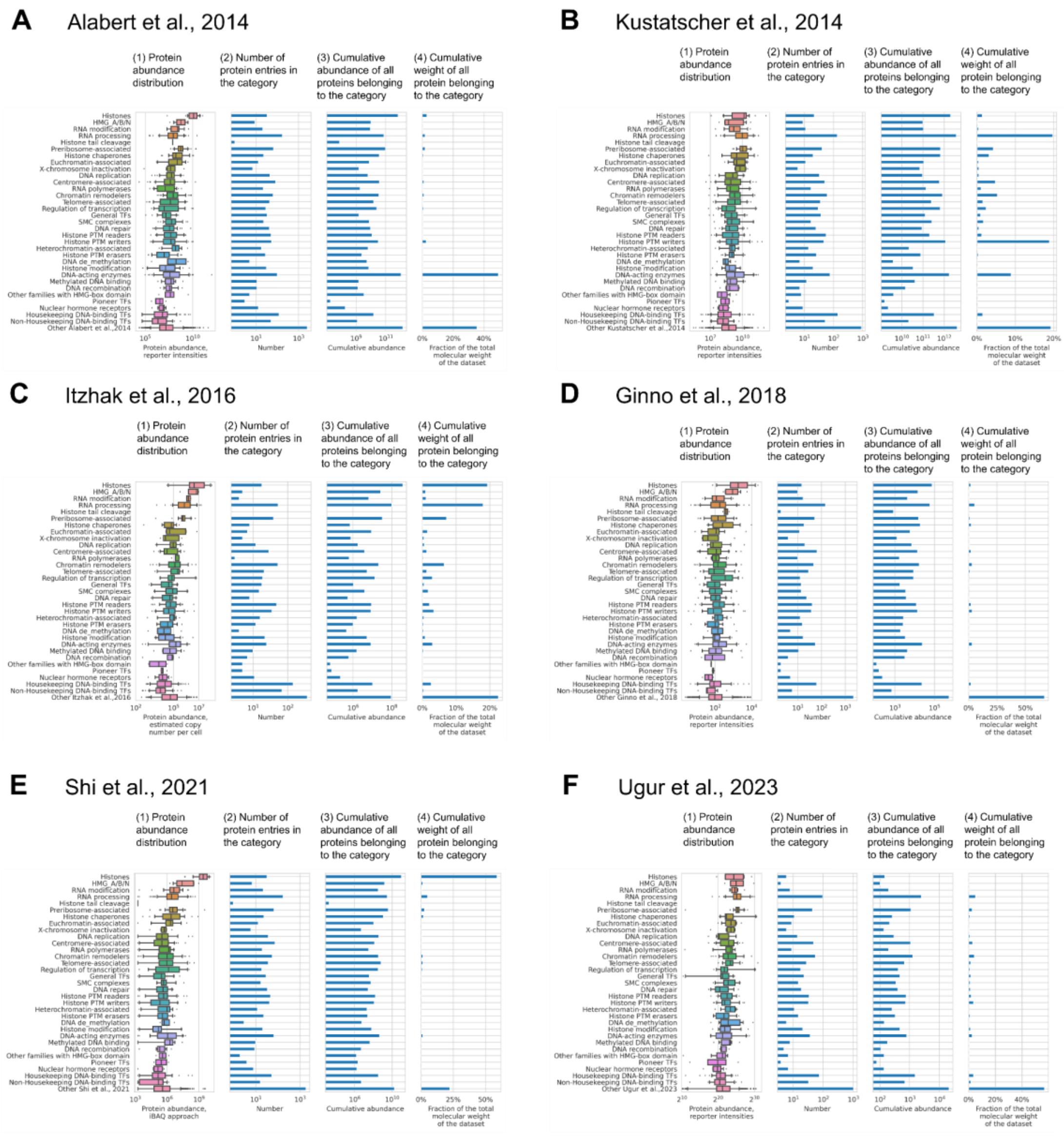

**Supplementary Figure SF4_4.** These are versions of Figure 4C from the main text showing chromatin protein abundance for SimChrom categories, but only for SimChrom proteins that were detected in MS-based chromatome/nucleome studies. The figure contains analysis for following chromatomes/nucleome (**A**) Alabert et al., 2014; (**B**) Kustatscher et al., 2014; (**C**) nucleome study from Itzhak et al., 2016; (**D**) Ginno et al., 2018; (**E**) Shi et al., 2021; (**F**) Ugur et al., 2023.

##### 3.2. Physico-chemical properties and amino acid composition

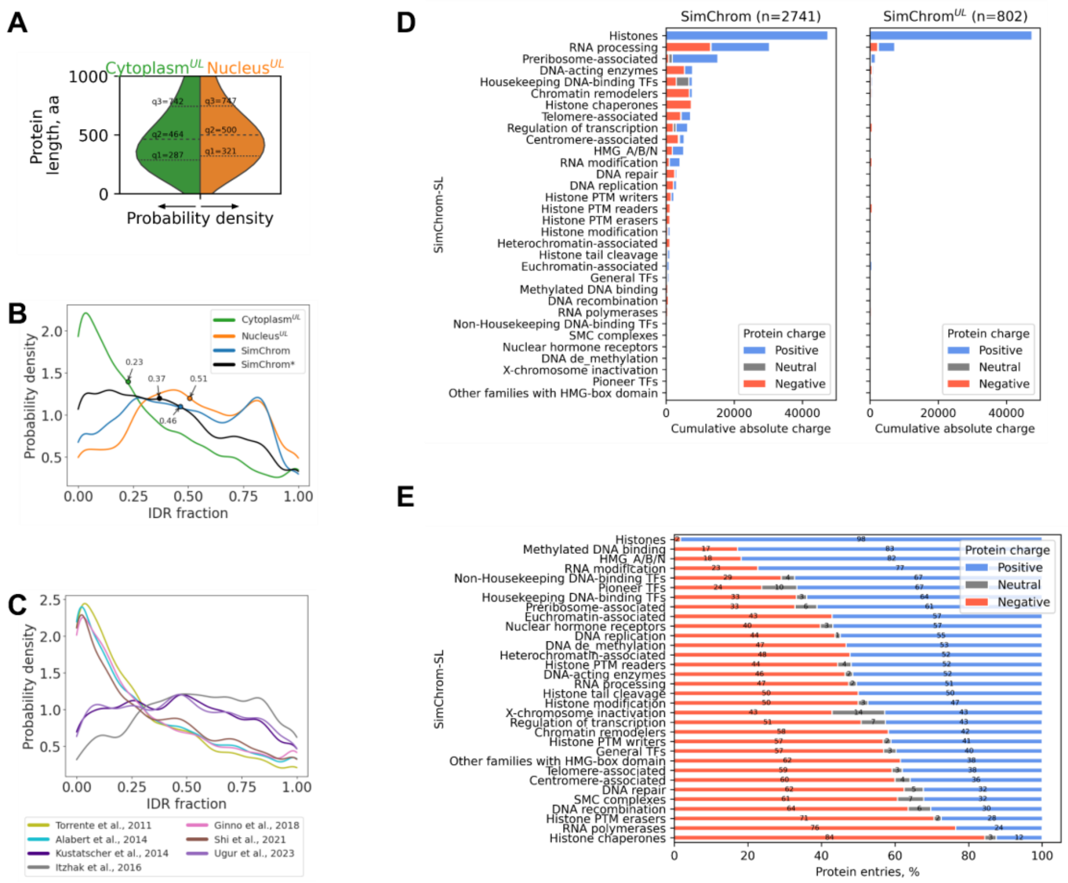

**Supplementary Figures SF5_1.** **(A)** The distribution of protein length for nuclear and cytoplasmic proteins. The q2, q3 (median) and q4 also shown. **(B)** IDR fraction distribution of chromatin, nuclear and cytoplasmic proteins (SimChrom, NULOC_CS_UL, CYLOC_CS_UL datasets). The median values are also shown. SimChrom* means SimChrom proteins without proteins from “Non-Housekeeping DNA-binding TFs” and "Housekeeping DNA-binding TFs" categories. **(C)** IDR fraction distribution for protein sets from MS-based chromatomes and nucleome. **(D)** Cumulative absolute charge of chromatin proteins from SimChrom (left) and chromatin proteins uniquely localized in the nucleus (right). **(E)** The fraction of protein entries with different types of charges (positive, negative, neutral) according to SimChrom categories.

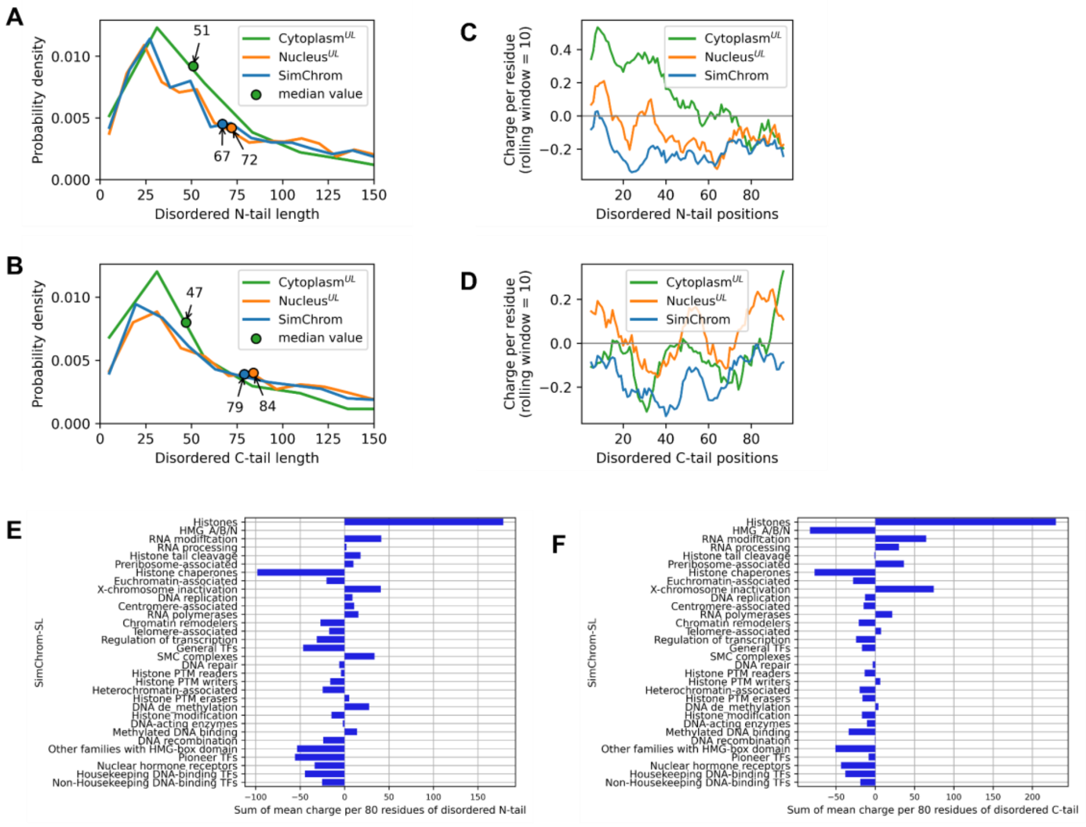

**Supplementary Figures SF5_2. (A-B)** The distribution of disordered tails' (N and C, respectively) length for chromatin, nuclear and cytoplasmic proteins. The medians are also shown. **(C-D)** Charge profile for N- and C-tails, averaged per amino acid residue with rolling window (10 residues) for each set of proteins (chromatin and uniquely localized in the nucleus and cytoplasm). **(E-F)** The charge of C- and N- disordered chromatin tails for each SimChrom category. For each set of proteins for each residue the mean charge was calculated, the bars represent the sum of these means. The analyzed length of disordered tails was less than 80 residues.

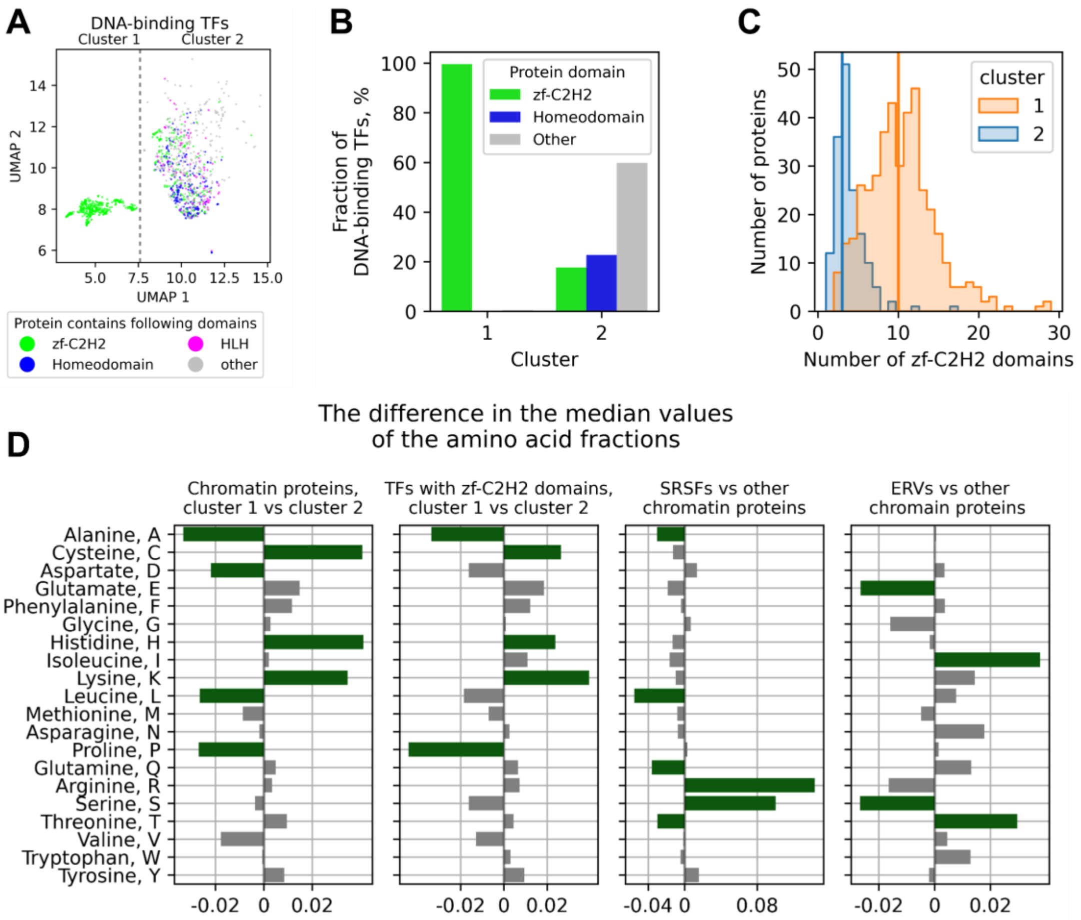

**Supplementary Figures SF5_3. (A)** This is a version of Figure 5K from the main text showing grouping of chromatin proteins by their amino acids composition using UMAP dimensionality reduction technique, but only transcription factors containing distinct DNA-binding domains are shown in the plot (the UMAP projections are the same as in Figure 5K calculated using the full set of chromatin proteins). **(B)** The fraction of DNA-binding transcription factors with zf-C2H2, homeodomain or others in the clusters from UMAP map in panel (A). **(C)** The number of DNA-binding TFs with different numbers of zf-C2H2 domains from clusters 1 and 2, median values are shown. **(D)** The difference in the median values of the amino acid fractions. The subpanels show the following comparison of: 1) chromatin proteins in cluster 1 with proteins in cluster 2; 2) TFs with the zf-C2H2 domain between clusters; 3) (Serine/Arginine-Rich Splicing Factor family (SRSFs) with other chromatin proteins; 4) proteins of the Endogenous retrovirus group (ERVs) with other chromatin proteins.

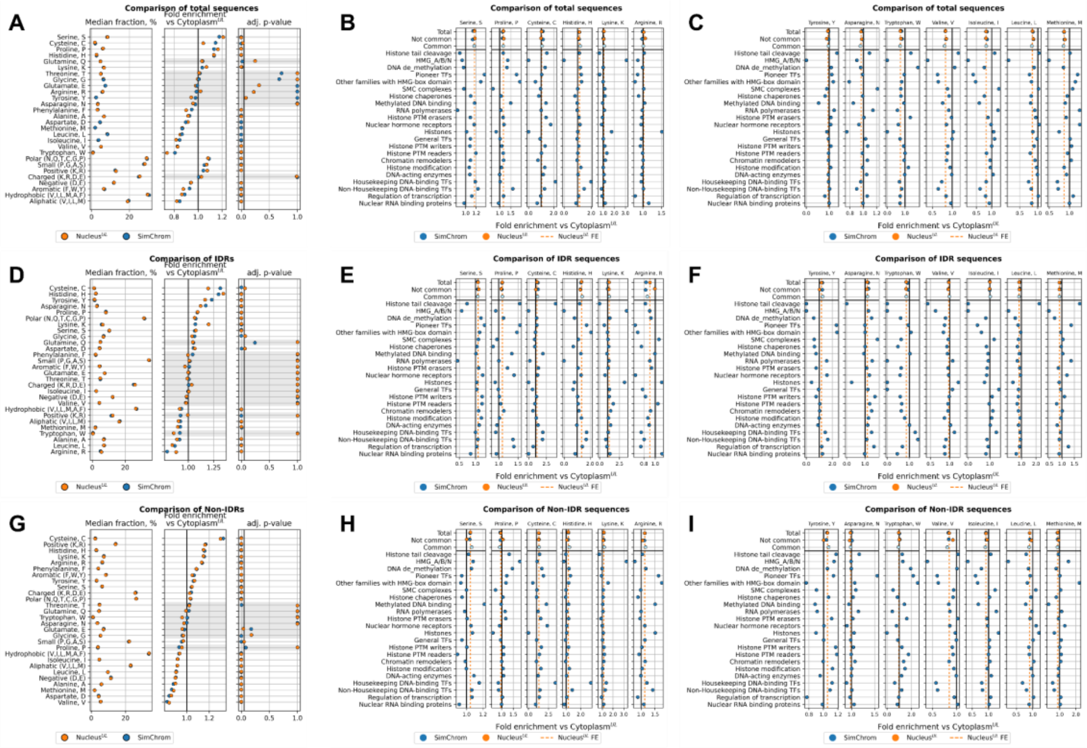

**Supplementary Figures SF5_4.** Comparison of amino acid composition between chromatin and uniquely localized nuclear proteins relative to cytoplasmic proteins (SimChrom, NULOC_CS_UL, CYLOC_CS_UL datasets). The comparison is done separately for the total protein sequence (**A,B,C**), IDRs (**D,E,F**), and non-IDRs (**G,H,I**). Subplots (**A, D, G**) presents the median fractions of amino acids for chromatin and nuclear proteins (subpanel 1 on each plot), the fold enrichment (FE) of these fractions relative to the cytoplasmic proteins (subpanel 2 on each plot), the black line indicates FE = 1. The adjusted p-value is shown for the statistical tests (Mann-Whitney test) comparing the median values of amino acid fractions for chromatin and nuclear proteins with the cytoplasmic ones (subpanel 3 on each plot). Gray highlights indicate a lack of statistical significance (adj. p-value > 0.05). Detailed analysis of the distribution of the selected amino acids in the total sequence of proteins belonging to respective SimChrom-SL protein categories is presented in panels **(B-I)**: enriched amino acids in chromatin proteins are shown in panels **(B,E,H),** depleted - in panels **(C,F,I)**. In the top of each plot (the first three rows) the following datapoints for the fold enrichment are given: “Total” - for all proteins from SimChrom or NULOC_CS_UL datasets (the latter also depicted by dashed line), “Common” – for common proteins among SimChrom and NULOC_CS_UL datasets, “Not common” – for proteins not present in the partner dataset (e.g., for SimChrom those present in SimChrom but absent in NULOC_CS_UL will be depicted, and vice versa for Nuclear_UL dataset).

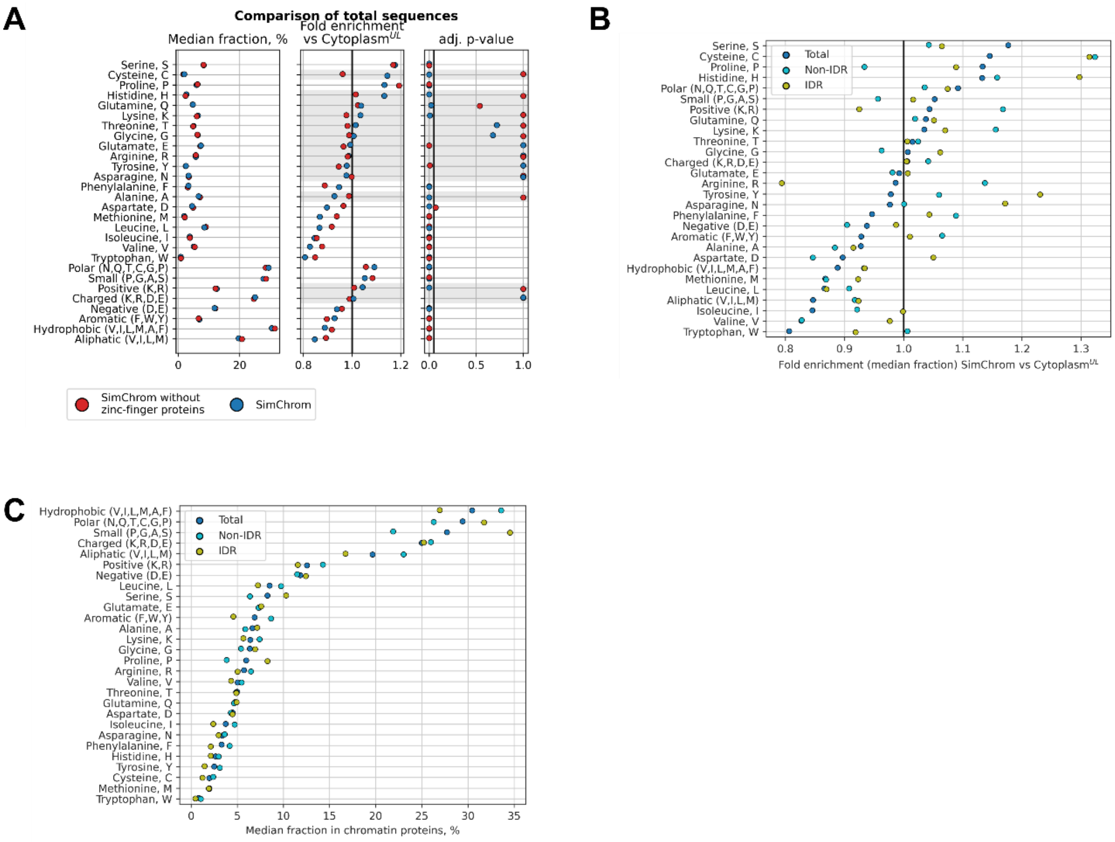

**Supplementary Figures SF5_5. Additional comparisons of amino acids composition for different protein groups. (A)** Comparison of amino acid composition in chromatin proteins and chromatin proteins without zf-C2H2 containing proteins relative to cytoplasmic proteins. The median fraction of amino acids for protein subsets (subpanel 1), the fold enrichment (FE) relative to the cytoplasmic proteins (subpanel 2), where the black line indicates FE = 1. The adjusted p-value is shown for the statistical tests comparing the median values of chromatin and chromatin proteins that lack zinc-finger domains with the cytoplasmic ones (subpanel 3). Gray shading indicates values that lack statistical significance (adj. p-value > 0.05). **(B)** Fold enrichment of amino acids' median fractions in chromatin proteins vs uniquely localized cytoplasmic ones, total sequences, IDRs and non-IDRs were analyzed separately. **(C)** Median value of amino acids' fractions in chromatin proteins for total protein sequences, IDRs and non-IDRs.

#### 3.3. Domain composition of chromatin proteins and identification of novel structural domains

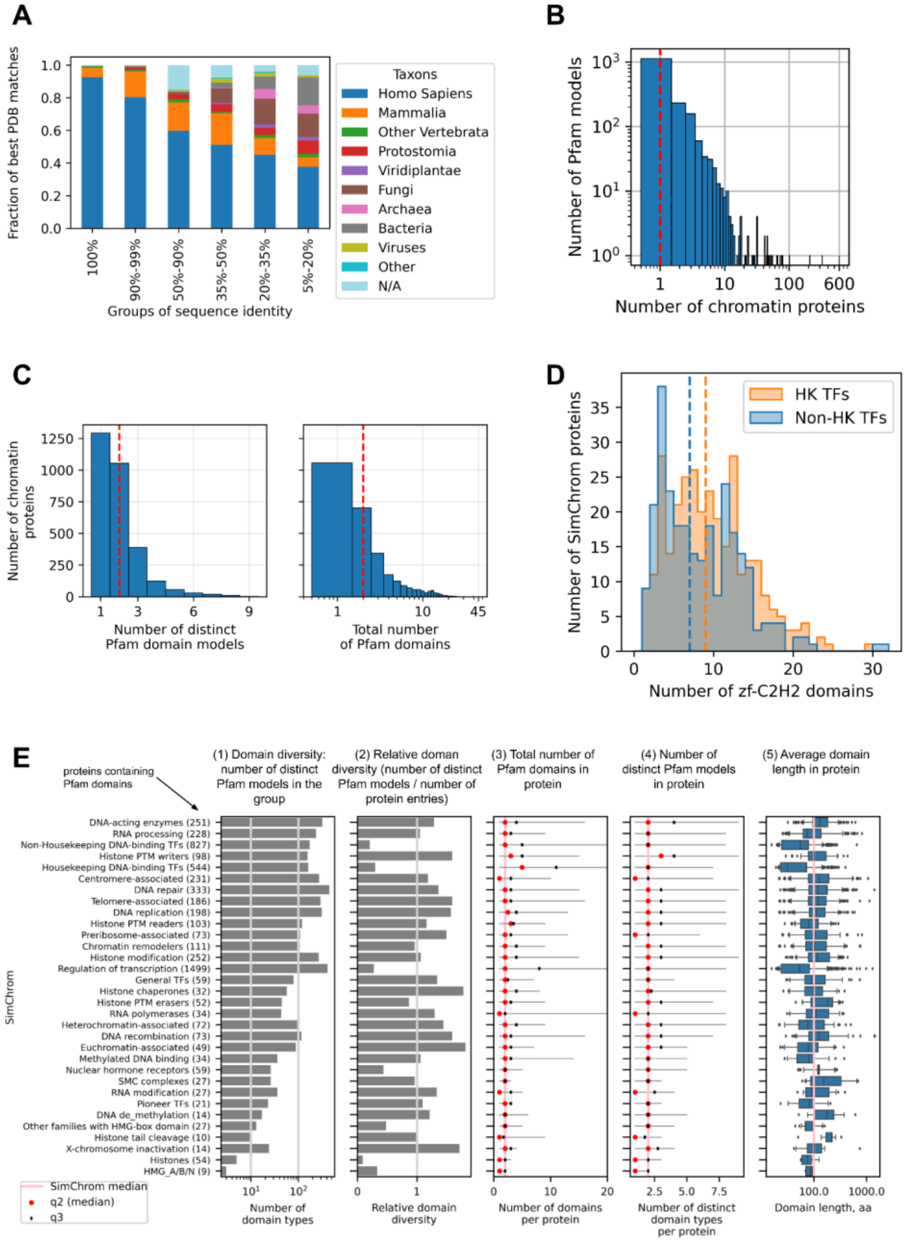

**Supplementary Figures SF6_1.** (**A**) Taxonomic distribution of source organisms for PDB structures with domains homologous to chromatin proteins (taxon of the best match to the structural domains identified by the TED resource, see Figure 6, Methods). (**B**) The histogram showing how many Pfam domain models (Y-axis) are found in exactly N (X-axis) chromatin proteins. One can see that the majority of Pfam domain models are represented only by domains found in one chromatin protein. The red line indicates median values. (**C**) The distribution of chromatin proteins according to the total number of Pfam domains identified in proteins (see also **Supplementary Table ST15**). (**D**) The distribution of proteins according to the number of zf-C2H2 domains in Housekeeping and Non-housekeeping DNA-binding transcription factors (HK TFs and Non-HK TFs, respectively). The lines indicate median values (7 and 9). (**E**) Analysis of functional domain diversity in chromatin proteins as identified by the Pfam database for proteins belonging to different chromatin categories according to SimChrom classification. Subpanel 1-5 represent various characteristics. This is the same as Figure 6E but SimChrom classification instead of SimChrom-SL classification is used.

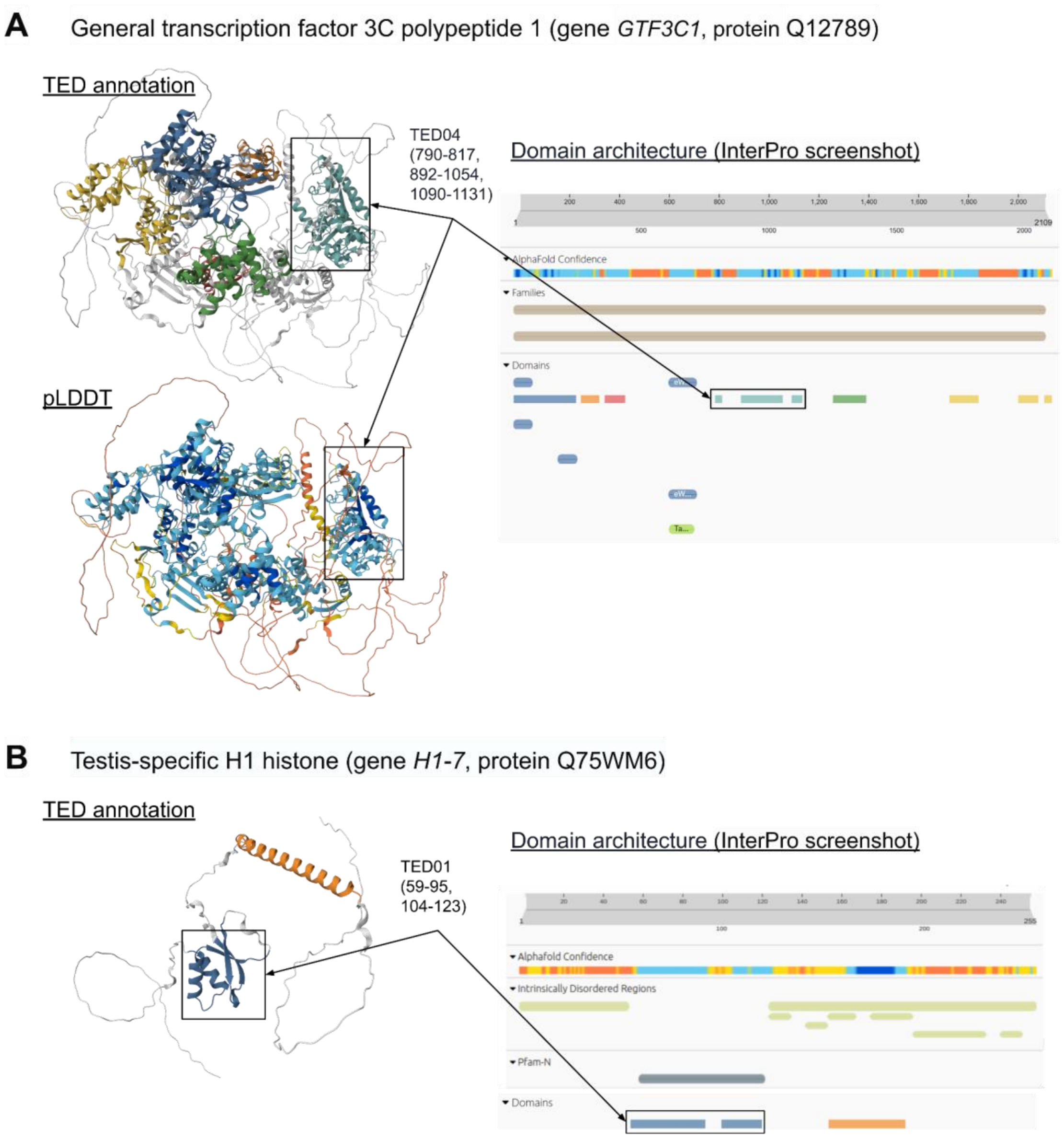

**Supplementary Figures SF6_2.** The examples of novel structural domains identified in chromatin proteins: structures, colored by TED domain annotation and AlphaFold2 pLDDT score, and its annotation in InterPro (screenshot). (**A**) General transcription factor 3C polypeptide 1 (gene GTF3C1, protein Q12789). (**B**) Testis-specific H1 histone (gene H1-7, protein Q75WM6).

**A**

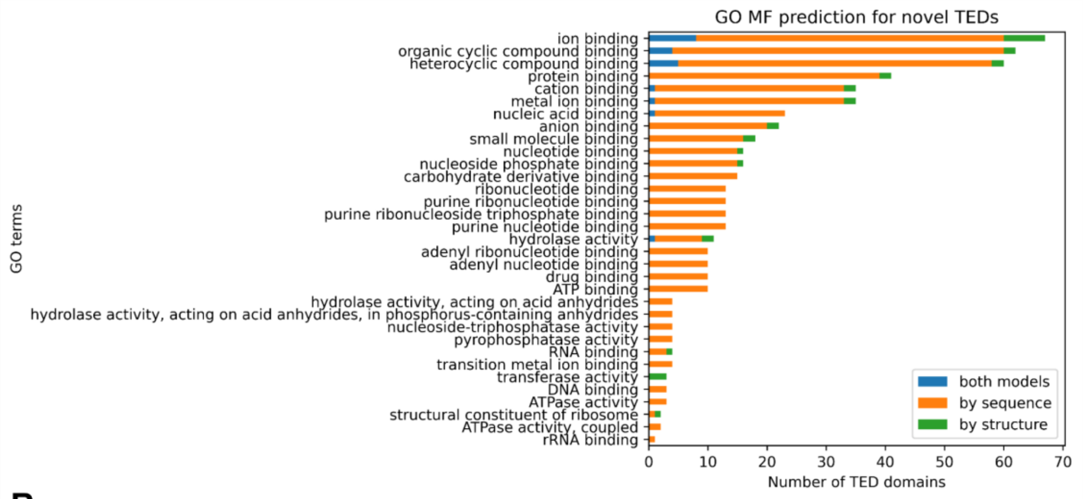

**B**

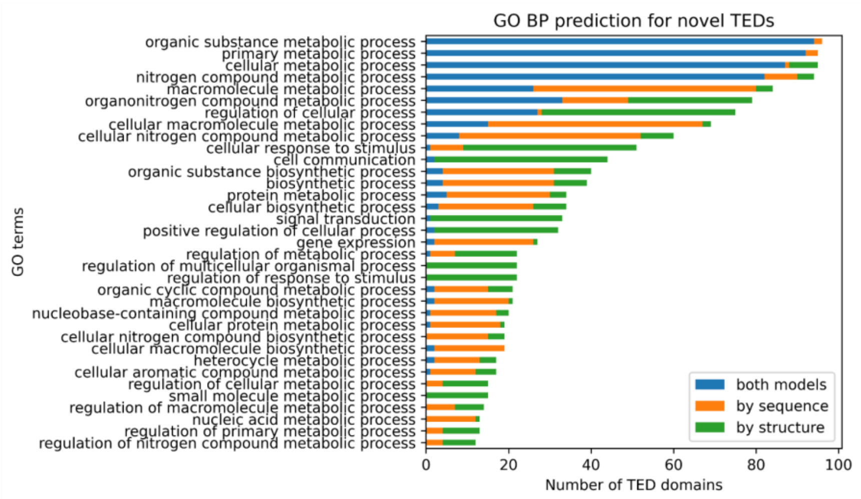

**Supplementary Figures SF6_3.** Predictions of GO molecular function (MF) (**panel A**) and biological processes (BP) terms (**panel B**) for novel structural domains without information in other DBs according to InterPro.

##### 3.4. Multivalent interactions in chromatin proteins

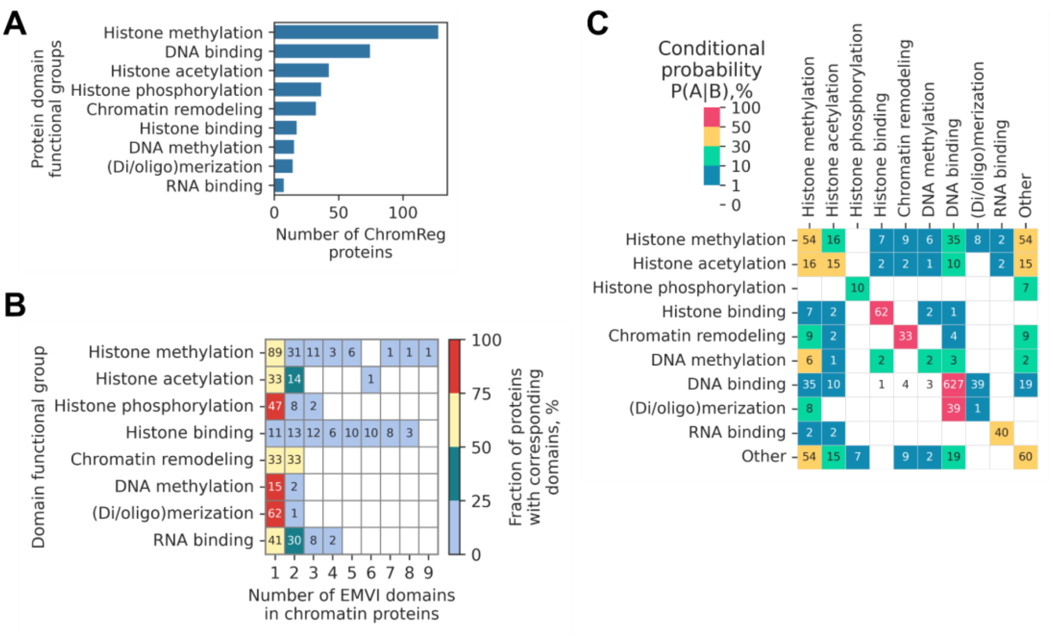

**Supplementary Figures SF8_1.** (**A**) The number of chromatin regulator proteins that contain EMVI-domains of certain groups. (**B**) The number of chromatin proteins with different numbers of EMVI-domains belonging to different groups ('DNA binding' domain functional group is not shown). (**C**) Co-occurrence of EMVI-domains belonging to different functional groups in chromatin proteins. The values indicate the estimated conditional probabilities to find in a chromatin protein a domain specified in the column name given that a domain specified in the row name is already present.

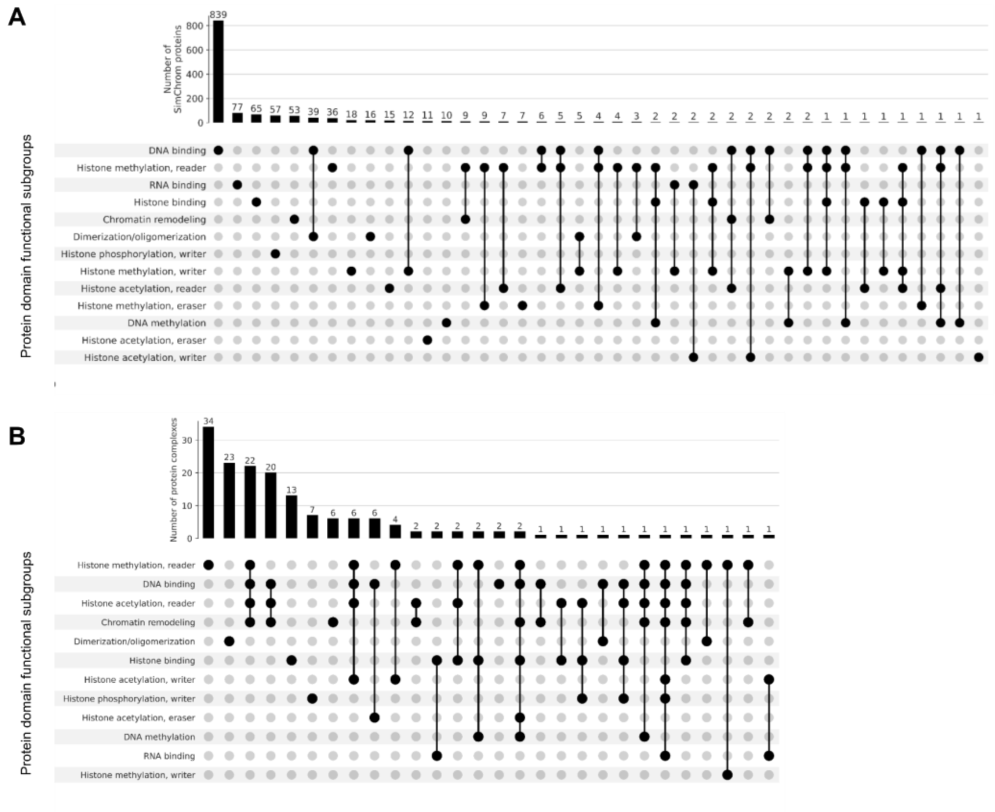

**Supplementary Figures SF8_2.** The UpSet plot shows combinations of EMVI domains classified by their functional groups/subgroups in chromatin proteins (panel **A**) and protein complexes that exclusively contain chromatin proteins (panel **B**).

### Supplementary Results and Discussion

#### 1. Sources of information about chromatin and nuclear proteins and their critical evaluation

This section includes supplementary results and discussion to section 3.1. Sources of information about chromatin and nuclear proteins and their critical evaluation in the main text.

##### A note on the distinction between and definition of nuclear proteome and chromatome

Nuclear proteome and chromatome are two terms that are historically used to describe the protein content of the nucleus and the proteins associated with genome packaging, maintenance and functioning (see Figure 1A for modern view of nucleus structure). The exact distinction between these two terms may be fuzzy and is often based on different consensus (protein localization or functional classification ontologies) or operational (experimental based extraction techniques) definitions. During interphase when the nucleus envelope is intact the chromatin proteins obviously reside inside the nucleus and are a part of the nuclear proteome. Hence, terminologically the nucleome seems to be more straightforwardly defined just by the protein contents of the nucleus. However, during mitosis and meiosis once the nucleus disintegrates as a distinct organelle, the situation becomes more complex. During these stages of the cell cycle there are no nuclear proteins *per se* while chromatin proteins can still be defined as those associated with the DNA in chromosomes.

Another debatable question is whether all of the proteins inside the nucleus can be considered chromatin proteins (even if nuclear envelope proteins are set aside). According to one modern view apart from the chromatin compartment the nucleus contains also interchromatin compartments [2] and nuclear bodies enriched in RNA and protein complexes (*e.g.*, nucleolus, nuclear speckles). Historically the soluble fraction of nuclear proteins was attributed to nucleosol or nuclear sap. However, to say that proteins localized in these compartments do not interact with genomic DNA at least transiently would be an oversimplification. Nucleoplasm proteins mostly also interact with genomes, some parts of the genome interact with the nucleolus (the so-called, nucleolar associated domains or NADs). Even proteins of the nuclear envelope – lamins do interact with the genomic DNA (*e.g.*, forming lamina-associated domains, LADs).

##### 1.1. Analysis of chromatin proteins' representation in the GO database and other protein-function oriented databases

The most comprehensive gene and gene products classification resource to date is GeneOntology (GO), which classifies proteins according to the three interrelated ontologies describing molecular function, biological processes, and cellular components (called aspects). GO annotates 97% of all proteins of the human reference proteome (as provided by UniProt), but this classification also has several drawbacks. GO combines many categories (currently around 42 thousand terms) of various scope describing various aspects of gene functioning connected via different types of relationships (such as "A is B", "A is part of B", "A regulates B", "A occurs in B", *etc.*) in a non-treelike structure (directed acyclic graph). This complex intertwined hierarchy of GO makes it difficult to get a holistic picture of various chromatin protein groups, and apply a reductionist way of thinking while interpreting the results of bioinformatics analysis of protein sets made using GO classification. Another drawback is that GO omits categories that are historically well established in the community of chromatin researchers (*e.g.*, such categories as "histone proteins", "high-mobility group proteins", *etc.*), again hampering interpretation of GO-based data analysis using the established knowledge (see discussion below).

The GO cellular component term "chromatin" is defined broadly as "the ordered and organized complex of DNA, protein, and sometimes RNA, that forms the chromosome." Consequently, functionally relevant chromatin terms — such as "nucleosomal DNA binding", "DNA-binding transcription factor activity', 'Histone H3K27 DNA-binding transcription factor activity', 'Nucleus', 'Histone H3K27 monomethyltransferase activity" — may not be linked hierarchically to the chromatin GO node, resulting in incomplete overlaps between protein lists. Based on our analysis, over 500 functionally defined chromatin proteins (inferred from literature and other sources) are absent from GO annotations (see **Supplementary Figure SF2_1A**).

While the GO provides nearly comprehensive coverage of the proteome, its classification structure presents challenges for extracting or comparing specific protein sets. The GO hierarchy is complex, with heterogeneous relationships between nodes, overlapping protein annotations across terms, and varying levels of detail and completeness. For instance, manually inspecting GO terms associated with chromatin-related keywords (*e.g.*, "DNA", "transcription", "histone", "RNA polymerase") is impractical due to their sheer volume (>100 terms; see **Supplementary Figure SF2_1B**). Furthermore, GO terms inherently include proteins from all child terms, which can lead to unintended inclusions. For example: the term "DNA-templated" transcription incorporates "mitochondrial transcription"; "gene expression" encompasses functionally distinct processes like "protein maturation" and "translation".

When compared with the EpiFactors database [3] (containing epigenetic regulator protein entries obtained by text mining) 46% of entries in EpiFactors are missing from the list of GO 'chromatin' proteins, see Figure 2D**.** Moreover, while a specialized review by Hammond et al., 2017 [1] lists 35 proteins of histone chaperone category, the GO term "histone chaperone activity" includes only 14, with just 6 overlapping entries (see **Supplementary Figure SF2_1D**).

Several functionally important but small chromatin protein categories are entirely missing from GO, including: HMG proteins, Histone tail cleavage proteins, general TFs. Even for well-annotated classes like histones, inconsistencies persist. Most are classified under "structural constituent of chromatin", but this term also includes two non-histone proteins (*HMGA1* and *LMNTD2*).

The Gene Ontology offers the most comprehensive coverage, annotating nearly the entire human reference proteome. In contrast, chromatin/epigenetic regulator databases typically include 400–800 proteins, while protein class-specific databases vary significantly in scope, ranging from as few as 30 proteins (*e.g.*, chromatin remodelers of the SWI/SNF family) to over 1500 (*e.g.*, transcription factors). Despite their utility, none of these resources fully captures the complexity of chromatin-associated proteins, either in terms of protein coverage or functional classification. While chromatin/epigenetic regulator databases encompass key cofactors for certain protein complexes, they exclude critical categories such as transcription factors, RNA polymerase subunits, DNA-modifying enzymes, DNA repair machinery, and HMG proteins. Conversely, protein class-specific databases are limited to at most six functional categories in total, including histones, chromatin remodelers (from select families), histone post-translational modification (PTM) writers/readers/erasers, and transcription factors. Additionally, several chromatin-related protein classes have been reviewed as the gene group in HGNC (*e.g.*, 'High mobility group', 'DNA polymerases') or in the literature but lack dedicated database resources. Examples include histone chaperones [1], SMC complexes [4], HMG proteins [5], pioneer transcription factors [6], nuclear RNA-binding proteins [7,8], and histone tail cleavage enzymes [9]. The full list of chromatin-associated protein class specific sources are available in **Supplementary Table ST1**. The absence of centralized repositories for these protein classes highlights a critical gap in current bioinformatics resources.

The revealed problems for using GO directly for extracting a set of chromatin proteins arise from several factors (see also [10] for a broader discussion of GO applicability): 1) protein multifunctionality: many proteins participate in diverse processes or localize to multiple compartments, 2) ambiguous term definitions: brief GO descriptions may lead to inconsistent interpretations, 3) annotation delays and errors: lag times in updates and propagation of errors in curated datasets, 4) curation bias: well-studied proteins are annotated more thoroughly than niche categories.

**1.2. Detailed comparative analysis of nuclear proteins subcellular localization between UniProt, HPA, and OpenCell**

Note: **Interactive Figure 2** (https://simchrom.intbio.org/#localization) is the interactive version of **Supplementary Figure SF2_2**, which is the key source of information for the analysis presented below.

In this section we analyzed in detail the information on subcellular localization on nuclear proteins provided by three database/proteome wide studies: UniProt, HPA, OpenCell. Each resource uses its own hierarchical subcellular localization ontology and a system of evidence tags (UniProt), reliability scores (HPA) or annotation grades (OpenCell) to annotate proteins with localization terms with different levels of confidence or types of evidence. From the three databases we obtained the lists of proteins that had their localization annotation(s) supported by at least one evidence tag (in the case of UniProt), reliability score higher than "uncertain" (in the case of HPA) or annotation grades better than the worst one (in the case of OpenCell) (see Methods 2.1.1 for details). UniProt is currently the main state-of-the-art reference data source about the human proteome, estimating it at 20225 proteins (release 2022_2). From these around 14 thousand are "localized" according to our criteria (see **Supplementary Figure SF2_2A**). HPA has so far experimentally analyzed around 13 thousand human proteins from which 12 thousand are localized. Interestingly, the localized protein sets of UniProt and HPA overlap only by 60-70% (8981 proteins, see **Supplementary Figure SF2_2A**), leaving around three thousand proteins, whose localization is characterized by HPA, but not available in UniProt. OpenCell has so far analyzed only a minor fraction of human proteome (all of them have been localized), although it provides localization information for another 37 proteins that are not "localized" by either UniProt or HPA (see **Supplementary Figure S2_2A**). In total the three databases collectively provide localization information for 17335 proteins amounting to 86% of the known human proteome. The fraction of "localized" proteins relative to each dataset constitutes 70%, 93%, 100% of the total number of proteins listed in UniProt, HPA, and OpenCell, respectively (see **Supplementary Figure S2_2A**).

**We next set to analyze information provided by these resources about the nuclear/chromatin proteins including their sublocalization**. The representation of different localization ontologies, the number of proteins annotated by the respective annotation terms (each individual term had to be supported by the criteria used to classify a protein as "localized" described above) and the overlap between the number of proteins annotated by different terms in UniProt and HPA is given in **Supplementary Figure SF2_2A**. We note that localization ontologies used by the three resources are somewhat different reflecting different approaches to protein classification, other ontologies used as basis (*e.g.*, GeneOntology (GO) in the case of UniProt), evolution of our understanding of chromatin organization (*e.g.*, some obsolete terms such as "Nucleus matrix" may be present in UniProt [11]), resolution of experimental methods and data sources used to classify protein localization. All ontologies converge on the presence of the following four common localization entities within the nucleus "Nucleus envelope/Nucleus membrane", "Nucleolus", "Nucleoplasm", "Nuclear bodies/speckles/punctae". However, the hierarchical relations between these terms within each ontology and their further subdivision into sub-localization (subcompartment) entities differ. Moreover, the terms with the identical names may be used at different levels of the hierarchy as both the names of specific annotation terms and groups of annotation terms, these have to be treated as separate terms when using this ontology for the analysis of protein sets (*e.g.*, the terms "Nucleoplasm" in HPA, or "Nucleus envelope" in UniProt, see **Supplementary Figure SF2_2A**). The term "Chromatin" is only explicitly present in OpenCell, suggesting that chromatin proteins in UniProt and HPA should be found in localization subcategories (Nucleus, Nucleoplasm, etc.). However, UniProt has a separate localization term "Chromosome", which according to its classification is a separate entity not being a part of "Nucleus". This term was inherited by UniProt from GeneOntology (GO) cellular component classification. According to GO "chromatin" is a part of "chromosome", while the term "chromosome" may be used to describe both components located inside (e.g., "nuclear chromosomes") and outside (e.g., "mitochondrial chromosomes") the nucleus. To add to this complexity, HPA considers "mitotic chromosomes" a part of "Nucleoplasm", but does not explicitly include the term "Nucleus" in its cellular sublocalization ontology (the subcellular localization ontology at the most general level already is subdivided into "Nucleoli", "Nucleus membrane" and "Nucleoplasm", as well as, "Cytosol", "Mitochondria", etc). The above-mentioned discrepancies reflect the dynamic complexity of cellular organization, our constantly evolving understanding of nuclear organization, and the resulting difficulty in describing subcellular localization in a form of a simple hierarchical tree-like ontology.

**We next performed comparative analysis of the quantitative and qualitative composition of nucleomes as provided by the three resources**. We first addressed the question of the number of nuclear proteins known to date and the consistency in these estimates between the resources. The number of proteins having their localization assigned to the nucleus is estimated as 4717, 6500, and 640 by UniProt, HPA, and OpenCell respectively, which gives an estimate of 33%, 53%, and 49% of human proteome to be present in the nucleus according to these resources, respectively. The **liberal estimate** of the number of nuclear proteins (combined from three resources) is 8035, which amounts to 46% of the known localized proteome (17335 proteins in all resources combined).

**Analysis of multi-localization of nuclear proteins in other cellular compartments**

A significant proportion of the nuclear proteome is considered to be also localized in other cellular compartments. We have aggregated localization terms in HPA and UniProt ontologies into the following generalized categories Nucleus, Cytoplasm, Endomembrane system (see Methods Section 2.1.1). **Supplementary Figure SF2_2B** shows the multi-localization of proteins across these categories in the form of Venn diagrams. According to UniProt and HPA 41% and 39% of nuclear proteins are localized in cytoplasm, respectively, 12% and 22% of nuclear proteins are localized in the endomembrane system, respectively. Among the nuclear proteins whole localization is available in both HPA and UniProt, only for ~40% of proteins the two databases reach consensus for their unique nuclear localization (1262/3288).

**We next analyzed consistency between UniProt and HPA** in annotating nuclear proteomes. The overlap between the nucleomes from UniProt and HPA amounts to only 3288 proteins, meaning that only 51% of nuclear proteins reported by HPA are supported by UniProt, and vice versa 70% of nuclear proteins reported by UniProt are supported by HPA. However, since both HPA and UniProt simultaneously provide localization information only for 8981 proteins, it is better to estimate consistency between the resources only within this set of proteins. Among these 8981 proteins, nuclear proteins constitute 3896, 4699, and 5307 according to UniProt, HPA, and HPA or UniProt, respectively. Hence, we estimate the consistency between UniProt and HPA in annotating the proteins as nuclear at 62% (Jaccard similarity measure) (3288/5307). Hence, 2019 proteins were annotated as nuclear by one resource but not the other, and at the same time had localization annotation in both resources. These discrepancies may in part come from the fact that proteins may have multiple localization, and one of the resources misses one of the localizations. We see that  among these 2019 proteins, 1670 (83%) proteins were simultaneously localized in two or more major cellular/extracellular structures (Nucleus, Cytoplasm, Endomembrane system, Other (including Secretory/Extracellular) localization - see Methods Section 2.1.1) according to at least one of the resources. Among those for 1254 proteins HPA and UniProt were consistent in identifying at least one common localization, suggesting that the discrepancy in their nuclear localization between HPA and UniProt may be because one resource misses its additional localization in the nucleus, while they are consistent about their localization in other cellular compartments. For 765 proteins that were considered as nuclear only by one resource, no alternative common localizations were reported by both resources. To further understand potential sources of these discrepancies and annotation biases in HPA and UniProt we performed a cross-annotation analysis for this particular “discrepant” set of proteins: analyzed the annotations provided by one resource for the proteins that were considered nuclear by the other resource. The results of our analysis with respect to detailed annotation categories is presented in **Supplementary Figure SF2_3**. These “discrepant” proteins are mainly annotated as belonging to nucleus or nucleoplasm in one resource and fall into a wide range of categories in the other resource. Among this wide range of localization categories in both cases the cytoplasmic proteins form the largest category by size, but account only for 20%-30% of all “discrepant” proteins, and are statistically underrepresented (fold enrichment of 0.7-0.8, compared to the case if they were selected at random from the respective set of proteins - see Methods Section 2.1.1). Among nuclear proteins annotated by UniProt but whose annotation was missed by HPA (154 proteins), proteins with "vesicle" localization category according to HPA were overrepresented (1.3 fold enrichment). Alternatively, among nuclear proteins annotated by HPA, whose annotation was missed by UniProt (611 proteins), "secreted" and "extracellular matrix" proteins (according to UniProt) were mainly overrepresented (1.6 and 1.6 fold enrichments, respectively) with largest number of proteins, while overrepresented "chromosome" and "kinetochore" (6.1 and 2.7 fold enrichments, respectively) contains less than 20 proteins. *Taken together this suggests that discrepancies in nuclear protein localization between UniProt and HPA have a complex origin, some effects may be in part due to 1) the tendency of HPA to mislabel nuclear proteins as vesicular proteins (may explain up to around 30% of proteins with mis-annotated localization), and 2) the tendency of UniProt to mislabel nuclear proteins as belonging to the secreted proteins and proteins of the extracellular matrix (may explain up to around 15% of proteins with mis-annotated localization). However, (see main text section 3.1) one of the main sources of these discrepancies is also the incompleteness of the databases (especially, UniProt) with respect to annotating multilocalization of proteins (e.g., the databases may agree on cytoplasmic localization of a protein, but one database may additionally annotate it as nuclear).*

**We next performed comparative analysis for the number of nuclear proteins assigned to different subnuclear localization categories as provided by the three resources**, and particularly analyzed the quantitative similarities and differences between the data provided by UniProt and HPA (see **Supplementary Figure SF2_2A**). A conceptual difference between UniProt and other resources, is that UniProt provide subnuclear localization data only for 19% (882 proteins) of its nuclear proteome, while HPA and OpenCell *per se* do not have a localization term “Nucleus” and provide protein localization by default at a finer level. Particularly, the largest subset of nuclear proteins in HPA and OpenCell (5948 and 487 proteins, respectively) belongs to nucleoplasm category, while only 170 nucleoplasmic proteins are found in UniProt (245 more proteins in UniProt are classified as belonging to nuclear bodies, which is not considered are part of nucleoplasm by UniProt, but is considered by HPA). Among the consensus set of nuclear proteins between UniProt and HPA (3288 proteins) 95% of proteins annotated as nucleoplasm by HPA are annotated solely by the term “Nucleus” in UniProt (2781 out of 3053). The second largest category of 1191 proteins according to HPA are proteins belonging to the Nucleolus (this is not the second largest category in OpenCell, but OpenCell is biased towards the most abundant proteins), the same is true for UniProt (372 proteins), however, the number of proteins is much less in accordance with the small proportion of proteins that UniProt provides subnuclear localization annotation for. Among these 372 proteins only around 50% (185 proteins) are also annotated as belonging to Nucleolus by HPA. The consistency between other subnuclear localization annotations is even less (see **Supplementary Figure SF2_2A** and **Supplementary Table ST2**). For example, in the case of nuclear envelope/nuclear membrane only 46 proteins have a consistent annotation between UniProt and HPA, while for 368 proteins the annotations do not match. Taken together our current understanding of sub-nuclear localization annotations are not very much consistent between UniProt and HPA.

**The subnuclear multi-localization of nuclear proteins was another aspect of our analysis.** UpSet plots comparing the number of proteins that have multiple subnuclear localizations according to HPA and UniProt were used for this analysis (see **Supplementary Figure SF2_4**). To be able to compare the data between the two resources, given the difference in localization ontologies as described above, in this analysis we used a manually created common localization ontology consisting of four categories (nucleoplasm, nuclear membrane, nucleolus, nuclear bodies) separately mapped to HPA and UniProt as detailed in **Supplementary Table ST3**. According to HPA the majority of nucleoli proteins (59%) are also annotated as localized in nucleoplasm, which is not unexpected because nucleoli are suspended in nucleoplasm with proteins diffusing to and from the nucleoli. In the case of UniProt only 17% of nucleoli proteins are also localized in nucleoplasm, consistent with a small number of nuclear proteins annotated as nucleoplasmic proteins in UniProt. A similar situation is found when comparing multiple localization between nucleoplasm and nuclear bodies (48% and 16% overlap according to HPA and UniProt, respectively). Another interesting fact is that according to HPA the majority of nuclear membrane proteins (54%) have a second localization in the nucleoplasm (in the case of UniProt only 6% have the said location). This suggests that many of the inner nuclear membrane proteins may be tethered to the nuclear membrane transiently through interactions with the bona fide membrane proteins that have transmembrane domains.

Given the complexity of nuclear sublocalization ontologies, significant differences in certain aspects of data provided by UniProt, HPA, and OpenCell (as discussed above), and different comparative analyses between the datasets that may be done to answer various specific questions about nuclear localization/subnuclear localization/multi-localization of proteins, we have implemented an interactive data viewer based on the analysis present in this work. This viewer is available https://simchrom.intbio.org/#localization and allows to interactively select and analyze various subsets of nuclear proteome based on the data collectively provided by the three resources.

##### 1.3. Detailed comparative analysis of sets of chromatin proteins identified in MS-based studies

An important source of information about chromatome content may be obtained from MS-based studies of chromatin or nuclear extracts (reviewed in [12,13]). We aimed at comparing these data to the chromatome and nucleome datasets developed above. We have chosen a number of high quality studies published since 2010 with publicly available data that analyzed human chromatin protein content (Table 1, **Supplementary Table ST1**, **Supplementary Figure SF2_5**, **Supplementary Figure SF2_6**, **Supplementary Figure SF2_7**). These studies used different human cell lines (Torrente et al.,

2011 - HeLa S3; Kustatscher et al., 2014 - HeLa, MCF-7, HepG2, HEK293, U2OS, DT40; Ginno et al., 2018 - T98G (derived from a human glioblastoma multiforma tumor); Shi et al., 2021 - K562, Ugur et al., 2023 - hESCs). For additional comparison we also included nascent chromatomes (Alabert et al., 2014 - HeLa S3; Alvarez et al., 2023 - HeLa S3 and TIG-3 fibroblasts) and MS-based study that reported a set of proteins predominantly localized in the nucleus (Itzhak et al., 2016 - HeLa cells).

While aiming at analyzing the chromatin content directly, MS-based studies are not without known limitations that should be kept in mind during the analysis. One previously reported limitation is the difficulty of extracting the chromatin associated proteins without the contamination from cytoplasmic and mitochondrial proteins, another one is the difficulty in retaining the proteins that associate with chromatin dynamically, transiently, or are present in small amounts [12,13]. Over the years different approaches to chromatin extraction have been suggested: (1) techniques based on crude differential detergent/salt extraction of chromatin proteins (which are known to result in a substantial contamination by cytoplasmic proteins, in part due to the sticky nature of chromatin which captures other proteins during extraction [14], (2) techniques that rely on formaldehyde cross-linking prior to the biochemical extraction of the chromatin in order to reduce contamination by cytoplasmic proteins (ChEP, DEMAC), (3) techniques that add additional steps of chromatin fractions' purification using ultracentrifugation (DEMAC), (4) techniques that are based on adaptation of *in situ* Hi-C methods that enrich DNA-protein complexes through extraction of biotinylated DNA fragments and are claimed to capture more transiently interacting chromatin proteins (Hi-MS), (5) techniques that use MNase digestion to aid in extracting chromatin associated proteins (total MNase digestion). Additionally, some studies compared protein MS-intensities between different cellular components to reliably establish the protein contents of a particular cellular compartment (e.g. the approach used by Itzhak et al., 2016) or supplemented MS-analysis with machine learning classification of proteins based on the changes in protein MS-intensities when cells under different biological and biochemical conditions were analyzed (e.g., the approach used by Kustatscher et al., 2014). The chosen datasets employed various of the above mentioned techniques (see **Supplementary Figure SF2_5A**, **Table 1**). Additional known limitations of MS-based studies are due to: (1) the variable sensitivity of MS-based analysis techniques combined with highly variable levels of expression of chromatin proteins, (2) the variation in chromatin protein expression between the cell lines used for analysis, and the fact that not all chromatin proteins are expressed in all cell types and during all stages of the cell cycle and development, (3) difficulties in interpretation of MS-data for proteins with high sequences similarity, such as histones [15].

First, we analyzed the absolute number of proteins in MS-based chromatin datasets and the fractions of those datasets that were present in the SimChrom dataset of chromatin proteins that was developed by us as a part of this study (see Results Section 3.2) and nuclear localization-based datasets (NULOC_JT_NECF, NULOC_CS see Results Section 3.2) (see **Supplementary Figure SF2_5A**). The

MS-based datasets showed high variability, only 179 proteins were in common between the chromatin datasets only (see **Supplementary Figure SF2_5B**). The largest MS-based datasets of total chromatin (Shi et al., 2021; Ginno et al., 2018) and nascent chromatin (Alabert et al, 2014; Alvarez et al., 2023) contained around three thousand proteins – approximately the same amount as the number of proteins in SimChrom. However, surprisingly the number of proteins for these datasets that were present in SimChrom was small (25-35%). This low consistency with SimChrom was also observed in the Torrenete et al., 2011 dataset. These three datasets (Shi et al., 2021; Ginno et al., 2018; Torrente et al., 2011) were based purely on experimental chromatin extraction techniques (**Supplementary Figure SF2_5A**). Alternatively, the consistency with SimChrom was twice as high (50-65%) for the Kustatscher et al., 2014 and Itzhak et al., 2016 datasets. These datasets stand out from the other datasets. Kustatscher et al., 2014 used a machine learning classification approach based on MS-data signals with a manually provided chromatin proteins training dataset, which was based on literature and database mining. In Itzhak et al., 2016 the proteins were considered nuclear if their MS-measured intensity in the crude nuclear extract exceeded 85% of the global intensity. We hypothesize that these two latter datasets have better consistency with SimChrom dataset in part because by their design they are initially biased either by the information already available in the literature and various databases (in the case of Kustatscher et al., 2014) or by the selection of proteins that are preferentially localized in the nucleus and hence having higher chances to be described in the literature and databases (in the case of Itzhack et al., 2016).

When MS-based datasets were compared to the nuclear localization datasets, we observed the same tendency. Around 95% of proteins in Kustatscher et al., 2014 and Itzhak et al., 2016 datasets may be found in our broad nuclear localization dataset (NULOC_JT_NECF), while for other MS-based datasets the proportion was around 60-70% (Torrente et al., 2011; Alabert et al., 2014; Ginno et al., 2018; Shi et al., 2021) and 75-80% (Ugur et al., 2023; Alvarez et al., 2023). The same tendency was observed for the consensus NULOC_CS dataset (the portion of the MS-based datasets present in NULOC_CS varied between 30% and 73%).

To further understand the origins of these discrepancies we analyzed proteins that were found in MS-based experimental chromatin datasets but not present in nuclear localization datasets or SimChrom (**Supplementary Figure SF2_5C**). The Itzhak et al., 2016 dataset had a 98% overlap with NULOC_JT_NECF and its inclusion would not affect the results presented below. In total there were 2232 of such proteins, and only a minor fraction of these (94 proteins) did not have localization annotation in the databases (see **Supplementary Figure SF2_5D**). Five proteins were present in four total chromatin MS-based datasets (see **Supplementary Figure SF2_5C**), they included proteins encoded by the FLNB (Filamin B, actin-binding protein), GMPS (Guanine Monophosphate Synthase), CHERP (Calcium Homeostasis Endoplasmic Reticulum Protein), ILKAP (ILK Associated

Serine/Threonine Phosphatase), PLEC (Plectin) genes. However, the Ugur et al., 2023 dataset includes only PLEC and CHERP. While according to UniProt and HPA all of these are non-nuclear proteins predominantly localized in cytoplasm (with additional localizations in cytoskeleton, intermediate filaments, endoplasmic reticulum, Golgi apparatus), manual literature mining confirmed experimental evidence supporting the presence of these proteins in the nucleus (e.g. [16]). We additionally randomly selected 20 proteins from a subset of proteins that were reported by at least five out of seven chromatin MS-based studies (there were 195 such protein coding genes) and manually performed literature searches. From those 15 for 5 genes literature evidence was found suggesting their nuclear localization (CALR [17], PDIA4 [18], ABCF2 [19], SEC23B [20], EIF3D [21]). Hence, it may be stated that MS-base studies currently have predictive power to identify new chromatin proteins that are not annotated as such by the localization and functional databases.

It is not straightforward to estimate the potential contamination of MS-based datasets with non-nuclear proteins since one cannot come up with an ultimate reference set of non-nuclear proteins. Even for well studied proteins there are still chances that they may have axillary functionality in the nucleus that have not yet been experimentally characterized. Still to address this problem we relied on analyzing the above mentioned set of 2232 proteins using information available in GO. GO enrichment analysis revealed that these proteins were mainly associated with a diverse set of GO-terms related to non-nuclear organelles/compartments, cellular metabolism, protein translation and maturation suggesting that no important chromatin associated categories were missed during the construction of SimChrom dataset that could account for this discrepancy (**Supplementary Figure SF2_6**). Previous studies suggest that the results of MS-based studies may be contaminated by cytoplasmic and mitochondrial proteins [12,22]. In our analysis 2025 proteins were related to cytoplasm, 115 to Golgi vesicle transport according to GO. No enriched terms include mitochondria proteins. It is still possible that among 2232 proteins there are still nuclear proteins, whose annotation by GO does not account for their additional, moonlighting functions in the nucleus. For instance, 30 proteins were associated with translation ("translation", “translation initiation factor activity”) according to GO, many proteins of the translational apparatus are known to be moonlighting proteins with functional roles in the nucleus [23]. Similarly, Golgi apparatus cooperating with nucleus and ER in vesicular transport.

In a different type of analysis we looked at chromatin proteins that were not identified by the MS-studies but were included in our SimChrom dataset. There were 1246 such proteins (or 41% of SimChrom). According to the HPA classification of housekeeping proteins, 67% (839 proteins) of these were not housekeeping, consistent with the idea that they were missed by MS-based studies, because they were not expressed in the cell lines. However, it was also found that MS-based studies are biased towards identifying the housekeeping proteins. More than 75% of nuclear/chromatin proteins reported by the MS-based studies were from the housekeeping pool, while the average expected fraction of nuclear housekeeping proteins is around 62% (**Supplementary Figure SF2_7B**). Among the set of 1246 proteins not found in the MS-based studies the dominant SimChrom category was related to DNA-binding transcription factors (see **Supplementary Figure S2_7C**). 1148 DNA-binding TF were missed by MS-based studies, including 394 housekeeping TFs. This highlights another potential source of discrepancy - housekeeping TFs may be present in small amounts or be washed away during chromatin/nucleome extractions and thus be missed by MS analysis. To further understand the discrepancies between the MS-based datasets and SimChrom we performed enrichment analysis for the SimChrom categories in the experimental datasets (**Supplementary Figure SF2_8A**). It can be seen that categories related to DNA-binding transcription factors were mainly depleted in the MS-based datasets, consistent with the analysis described above. The separate analysis of housekeeping TF suggests that different experimental methods show high variability in their ability to recover TF in chromatin extracts. The Torrente et al., 2011 dataset includes only 8 housekeeping TF, while the Kustatscher et al., 2014 and Itzhak et al., 2016 datasets each contain 158. **The dynamic nature of transcription factors' interactions with chromatin likely explains these facts.** The most enriched categories were related to protein involved in RNA binding and metabolism, histone chaperone, remodelers and heterochromatin associated factors. This is likely related to the higher chances of these proteins to be detected due to their universal presence in the cells and high expression levels. Detailed analysis of the SimChrom categories representation in MS-based datasets further revealed some details of the differences between the datasets (**Supplementary Figure SF2_7C**). For instance, the ML-based Kustatscher et al., 2014 dataset was able to recover twice as many DNA transcription factors than Ginno et al., 2018 and Shi et al., 2021 datasets. The ratio of housekeeping and non-housekeeping TFs for Ginno et al., 2018 and Shi et al., 2021 datasets is the same, and is higher for the Kustatscher et al., 2014; Itzhak et al. 2016; Alabert et. al., 2014 datasets. Interestingly the nucleome Itzhak et al., 2016 dataset was also able to recover the same amount of TF as the Kustatscher et al., 2014 dataset, although the size of the dataset was three times smaller than Shi et al., 2021 and Ginno et al., 2018 datasets. This again points to the fact that TF may be lost during chromatin extraction. The representation of some other SimChrom categories had significant differences between the MS-based datasets likely attributed to the differential extraction probability. For instance, only 29% and 25% of RNA polymerases and histones, respectively, were present in the Ginno et al., 2018 dataset, while 50-60% were present in the Shi et al., 2021 dataset. A closer look at the histone proteins (for which currently information about all 62 expressed proteins is known [15,24,25] revealed that while certain tissue specific histone variants were missed as expected, all MS-based studies were not able to recover all canonical histones variants, especially for the H2B-histone (**Supplementary Figure SF2_8B**). For instance, from 14 canonical H2B proteins isoforms the Shi et al., 2021 dataset was able to recover six proteins (corresponding to the genes products of *H2BC1, H2BC3, H2BC4, H2BC13, H2BC18, H2BC26*), while the Itzhak et al., 2016 dataset four proteins (corresponding to genes products of *H2BC4, H2BC11, H2BC12, H2BC13*). While the canonical histones are likely to be expressed simultaneously in the cell, these discrepancies may be due to the sensitivity of MS-based analysis or expression variation in cell lines.

Taken together, our analysis indicates that MS-based chromatin datasets exhibit inconsistencies with subcellular localization or functional annotations available in UniProt, HPA, or GO. Up to 35% of chromatin proteins identified by MS-based techniques lack known nuclear localization according to the databases. Among these, approximately 30% of proteins identified simultaneously by at least three experimental studies may possess an additional nuclear localization that is not currently captured by the databases, but may have been reported in research papers. For other proteins, the most parsimonious explanation of their presence is the considerable contamination of chromatin extracts by mainly cytoplasmic proteins. From another point of view, many chromatin proteins annotated in the databases are not present in the MS-based datasets. While this is partially due to the limited number of genes expressed in the analyzed cell lines, our analysis also suggests that many proteins (such as housekeeping transcription factors) are lost during chromatin extraction, likely due to the dynamic nature of their interactions and their low expression. Finally, we showed that MS-based studies that filter their results by selecting the proteins that are highly enriched in the nucleus with respect to other cellular compartments, or use ML-assisted classification based on database/literature data, show better consistency with the information in the databases. However, this comes at the expense of decreasing the size of their datasets and likely limiting their ability to identify new proteins associated with the nucleus/chromatin.

#### 2. The SimChrom chromatin protein classification, the interactive SimChrom database and other reference datasets

This section supplements section 3.2. The SimChrom chromatin protein classification, the SimChrom dataset and other reference datasets in the main text. Note: **Interactive Figure 3** (https://simchrom.intbio.org/#classification) is the interactive version of Figure 3**,** which is the key source of information for the analysis presented below.

The largest subgroup of the SimChrom "non-histone proteins" category is the "DNA-templated transcription" group (1547 proteins), which consists of the proteins belonging to the "Regulation of transcription" subgroup (1514 proteins) and "RNA polymerases" subgroup (34 proteins). The "DNA metabolic processes" form the second largest protein subgroup (495 proteins) and include DNA replication, repair, and recombination machinery. "Nuclear RNA binding proteins" is another major group of proteins (309 proteins) present in SimChrom. There are a lot of RNA binding proteins in the cell (more than 1500 [26]), in the "Nuclear RNA binding proteins" category we aimed at including only those that are found inside the nucleus (see Methods Section 2.2 and **Supplementary Table ST4**), including those involved in preribosome formation, RNA processing and modification inside the nucleus. Other major subgroups of our classification included “Histone modification” (257 proteins), "DNA-acting enzymes" (258 proteins including DNA methylation and demethylation enzymes), “Centromere-associated” (241 proteins) (see Figure 3). Several specific subgroups of various scope containing proteins important for chromatin functioning were also included in our classification, such as “Histone chaperones" (32 proteins), ATP-dependent "Chromatin remodelers" complexes (114 proteins), “SMC complexes” (28 proteins, including cohesins implicated in chromatin loop extrusion).

To evaluate the contents of our SimChrom dataset we performed its cross-comparison to the localization-based datasets described above (NULOC_JT_NECF and NULOC_CS) (see **Supplementary Figure SF3_3**). The vast majority of SimChrom entries was also present in the NULOC_JT_NECF dataset (the broadest dataset that combined all nuclear protein entries from all protein localization databases at any level of confidence), which is consistent with chromatin proteins being only a subset of nuclear proteins. However, a minor subset of SimChrom (156 proteins) was not classified as nuclear by the localization databases (**Supplementary Figure SF3_3A**). To further understand the nature of this minor discrepancy we analyzed the SimChrom categories which contributed the most entries to this subset (**Supplementary Figure SF3_3C**) or where a significant proportion of entries in the respective category was absent in the localization databases (**Supplementary Figure SF3_4A**). In this subset 32 entries out of 156 were not annotated by the localization databases at all, 27 were considered by UniProt as only chromosomal (this is consistent with “Centromere-associated” category of SimChrom having the most entries not present in NULOC_JT_NECF), and 124 had only non-nuclear localization according to the localization databases. A manual review of the latter entries suggested that they included both bona fide nuclear proteins (such as histone acetyl and methyltransferases), other proteins such as ribosomal proteins, various kinases and proteins involved in mitochondrial DNA processing. Further research and data would be needed to clarify the localization status of the latter entries. The SimChrom category having the largest proportion of proteins absent from the localization databases was “Histone tail cleavage” (**Supplementary Figure SF3_3C**). This is consistent with the fact that for many of the histone tail cleavage enzymes (*e.g.*, metalloproteinases, cathepsins, neutrophil elastase) the histone tail cleavage activity in the nucleus is not their primary function and manifests only in specific conditions and cell development stages [9]. The comparison of SimChrom with the NULOC_CS (our stringent high confidence consensus dataset of nuclear proteins) showed a sufficiently higher number of SimChrom proteins that were not included in the NULOC_CS dataset (1208) (**Supplementary Figure SF3_3A**). This is not unexpected since only 44% of the human proteome has simultaneous localization annotations at sufficient levels of confidence from UniProt and HPA (see Results Section 3.1). The intersection of SimChrom and NULOC_CS datasets included 1837 proteins (**Supplementary Figure SF3_3A**), and may be considered as a set of chromatin proteins with a high level of confidence. To further validate that the subset of SimChrom that was not present in NULOC_CS (1208 proteins) represented nuclear proteins we performed GO enrichment analysis of this subset against a list of all GO-terms and then select non-nuclear associated terms for further analysis (see Methods Section 2.2, **Supplementary Figure SF3_4B**, **Supplementary Table ST8**). The analysis confirmed the low number of SimChrom proteins that were associated with *bona fide* non-nuclear GO categories (the "Centromere associated proteins" having the highest number – around a dozen out of 156 – of proteins that had "non-nuclear" GO-annotation terms, that belongs to charged multivesicular body proteins and dynactin complex subunits). Finally, as a byproduct of NULOC_CS and SimChrom comparison we find that 1459 proteins were present in NULOC_CS but not in SimChrom - this set may be considered as both a high-confidence set of nuclear non-chromatin proteins and a set of proteins that should be added to SimChrom. The GO analysis of this dataset did not reveal any clear GO categories that should have been included in SimChrom as categories related to chromatin functioning (see **Supplementary Table ST9**).

**Limitations** of the proposed chromatin classification SimChrom and its contents include the following: the absence of cell cycle control proteins and checkpoint signaling proteins, the lack of detailed classification for proteins involved in reading, writing and erasing DNA and RNA modifications. The 'Genomic location' categories require additional curation supported by experimental evidence to enhance the accuracy and reliability of their protein content. In addition, we did not consider the protein components of nonmembrane nuclear organelles whose proteins may also functionally interact with nucleic acids (directly or through phase separation). The classification does not include protein isoforms. Also, SimChrom is limited currently to human proteins only. These limitations will be addressed in the future versions of SimChrom.

#### 3. Analysis of the human chromatome

##### 3.1. The chromatome composition and abundance of chromatin proteins

This section supplements and expands section 3.3.1. The chromatome composition and abundance of chromatin proteins in the main text.

To understand chromatin functioning it is important to know the chromatome content not only in terms of the set of proteins associated with chromatin, but also in terms of their abundance (*i.e.*, the (relative) number of proteins per cell or organelle). Hence, we aimed at analyzing the available mass spectrometry data to address this question. The analysis of MS protein intensities from the experimental chromatome/nucleome studies discussed above, revealed a high degree of variability (see Figure 4A, **Supplementary Figure SF4_1**). For instance, the estimated relative mass of histone proteins varied from 0.1 to 58 % depending on the study, suggesting a high degree of bias due to different experimental techniques and analysis pipelines used to process raw mass spectrometry data (see Figure 4A). Hence, for further analysis we relied on the "whole-organism" protein abundance information available in PaxDB for *H. sapiens* [27]. PaxDb provides high quality information on protein abundance combined from many experiments with high coverage, dynamic range and interaction consistency (estimated consistency of abundance data with data on protein functional interactions) integrated over many cell types and conditions. Among the datasets available in PaxDb we have chosen two whole-organism datasets: the dataset with the highest proteome coverage ("H.sapiens - Whole organism (Integrated)" - covers 99% of human proteome according to PaxDb, referred to as "PaxDb_INT" in this paper) and the dataset with the highest interaction consistency score ("Whole organism, SC (Peptideatlas,aug,2014)" - covers 84% of human proteome according to PaxDb, referred to as "PaxDb_PA" in this paper), see **Supplementary Figure S4_1A**. Our analysis showed that with respect to PaxDb_PA, PaxDb_INT dataset has additional abundance information for around 2700 human proteins that almost exclusively have low levels of expression (less than 1 ppm, see **Supplementary Figure SF4_1B**). Among these proteins there are up to around 700 nuclear/chromatin proteins, hence we opted to use PaxDb_INT for general characterization of the abundance distribution of chromatin/nuclear proteins (presented in Figure 4B). PaxDb_PA dataset showed a higher consistency with respect to the relative abundance of functionally interacting chromatin proteins. The total abundance of different types of histone proteins (H3, H4, H2A, H2B) matched their expected equimolar ratio (see **Supplementary Figure SF4_2A,B**). Hence, PaxDb_PA was used for a detailed analysis of chromatin protein abundance distribution between chromatin protein groups and individual proteins (Figure 4C,D).

Around half of the whole-organism human proteome consists of low abundance proteins with expression levels of less than 1 ppm (~50%, see Figure 4B, **Supplementary Figure SF4_1C**, **Supplementary Table ST10**). The whole-proteome abundance distributions are positively skewed towards the low abundant proteins. In PaxDb_INT dataset this skewness is additionally supplemented by a second peak at the low abundance values (Figure 4B, **Supplementary Figure SF4_1A**). Among the low-abundance proteins only 25% of them correspond to the housekeeping proteins, while the proportion of housekeeping proteins among high-abundance proteins (abundance of more than 1 ppm) is 68% (see distribution in Figure 4B, **Supplementary Figure SF4_1C**, and **Supplementary Table ST10**). We used PaxDb_INT data to analyze abundance distributions of database-derived chromatome/nucleome protein sets discussed in the previous sections of the paper.

The NULOC_CS, NULOC_JT, and SimChrom datasets reported above all manifested distributions mirroring that of the whole proteome for PaxDb_INT data (see Figure 4B) with a significant proportion of proteins in these datasets still represented by the low-abundance proteins (40%, 44%, and 48%, for NULOC_CS, NULOC_JT, and SimChrom, respectively, see **Supplementary**

Figure SF4_1C, Supplementary Table ST10). We next aimed at understanding the types of proteins contributing to the low and high-abundance portion of the nucleome/chromatome. The proportions of house-keeping/non-housekeeping proteins in low- and high-abundance fractions for different database-derived datasets are given in Supplementary Figure SF4_1C and show that around 60% and 40% of chromatin/nuclear proteins are housekeeping ones for the high- and low-abundance fraction, respectively. As discussed above nuclear and chromatin databased-derived protein sets are on average enriched in housekeeping proteins with respect to the whole proteome (~58.5 % vs ~47%, see Supplementary Results and Discussion Section 1.3, Supplementary Figure SF4_1C, Supplementary Figure SF2_7A,B). This increase in the proportion of housekeeping proteins stems from both the increase of the number of low-abundance and high-abundance housekeeping proteins relative to the respective total numbers of low-abundance and high-abundance proteins in nucleome/chromatome datasets. A detailed analysis showed that the increase of the fraction of housekeeping proteins among the low-abundance ones was more than expected, while for the high-abundance ones was less than expected (under the assumption that high- and low-abundance fractions should contribute to the increase proportionally to the number of housekeeping proteins belonging to these fractions, see Supplementary Figure SF4_1C). For instance, under parsimonious considerations the overall increase in the fraction of housekeeping proteins for SimChrom with respect to the whole proteome (60% vs 47%) should imply the increase of house-keeping proteins' fraction among the low-abundance proteins from 25% to 32% (25*60/47=32), yet an increase to 38% was observed. **This highlights the important role that low-abundance housekeeping proteins play in chromatin functioning.** A more detailed analysis revealed that 64% of these housekeeping low-abundance chromatin proteins belong to the housekeeping DNA-binding transcription factors group (Supplementary Figure SF4_1D).

We next applied similar analysis to the sets of chromosome/nucleosome proteins identified in MS-based studies. The resulting distributions differed considerably from the distributions of database-derived protein sets discussed above, having a single maxima centered at higher values of abundance (see Figure 4B). **This fact again suggests that MS-based studies of chromatin are able mainly to recover highly expressed proteins and miss low expressed proteins** (low-abundance proteins are in the range of 1%-27% of the identified protein sets, Supplementary Table ST10, Supplementary Figure SF4_1C). The protein sets recovered by MS-based studies were significantly enriched in housekeeping proteins when compared to database-derived datasets (Supplementary Table ST10, Supplementary Figure SF4_1C). The abundance distributions varied between different MS-derived chromatin/nucleome protein sets. The chromatome studies based on extraction and/or cross-linking techniques (Alabert et al., 2023; Shi et al., 2021; Ginno et al., 2018; Torrento et al., 2011) had distributions shifted towards higher values of abundance, than the studies by Kustatscher et al., 2014; and Ugur et al., 2023 suggesting that the latter studies were also able to capture more chromatin proteins with low abundance. The nucleome study by Itzhak et al., 2016 was also able to capture more lower-abundant proteins.

We next aimed at understanding the abundance of different chromatin protein groups and individual chromatin proteins in the cell relying on our SimChrom-SL classification using PaxDb_PA abundance dataset. The resulting diagrams depicting abundance variations of chromatin proteins, belonging to different SimChrom-SL categories, the number of proteins belonging to the respective categories, and the cumulative abundances (calculated both as the total number of protein molecules and the total molecular weight of protein molecules belonging to each SimChrom-SL category) are presented in Figure 4C. To gain additional insights into the functioning of chromatin in Figure 4D we plotted the abundance values of highly expressed chromatin proteins (abundance of more than 1% of the H4 histone abundance) belonging to SimChrom-SL categories of the "Molecular function" or "Physico-chemical properties" type. Abundance data for all histone proteins and non-histone chromatin proteins with abundance more than 0.01% of Histone H4 is presented in the **Supplementary Table ST12**. It is important to note that many chromatin proteins have additional localization in other cellular compartments, hence the presented data reflects the overall abundance of the chromatin proteins in the cell rather than their abundance in the nucleus. To shed more light on the protein abundance in the nucleus we have also built diagrams analogous to Figure 4C only for 802 SimChrom proteins that are uniquely localized in the nucleus (according to our NULOC_CS_UL dataset) (see **Supplementary Figure SF4_3A**). These proteins are also highlighted in Figure 4D. As seen in panel 1 of Figure 4C chromatin categories vary substantially by their median abundance from 0.09 ppm to 570 ppm and there is still considerable variation in the abundance values within the categories. The most abundant chromatin protein is histone H4 (~11000 ppm), which is expressed by a family of genes almost exclusively coding the same protein sequence (except for H4C7, which has a negligible abundance). It is convenient to measure the abundance of all other proteins in fractions of H4 abundance (see Figure 4D). Each nucleosome contains two copies of H4 histones, therefore the numbers are also easily converted to relative abundance of chromatin proteins per nucleosome. Detailed analysis of histone protein abundance is in **Supplementary Table ST12** and shown in **Supplementary Figure SF4_2A,C**. The total number of core nucleosomal histone types H3, H4, H2A, H2B expressed by various genes sums up to similar numbers (~10400-10900 ppm) consistent with their equimolar association within nucleosome core particles. The cumulative abundance of H1 histones (~4500 ppm) suggests that slightly less than one H1 histone is associated with each nucleosome. The most abundant core histone variants are H3.3 (23% of H4, 2530 ppm), H2A.X (6.5%, 714 ppm), H2A.Z (10%, 1140 ppm), H2A.W (3.8%, 423 ppm). The least abundant histone variants are H2A.B and H1.7 (less than 1 ppm). Despite the relatively small number of protein coding human histone genes (108), many of which code for identical sequences, the cumulative abundance of histone proteins exceeds that of all other chromatin protein categories even if proteins with multiple localization are taken into account (see panel 2,3 in

Figure 4C). However, when the total molecular weight of proteins belonging to different categories is compared, the relatively small size of histone proteins (median ~15 kDa) results in them yielding the first place to RNA processing proteins (see panel 4, Figure 4C). Collectively the cumulative weight of proteins belonging to "Nuclear RNA binding proteins" category (that combines Preribosome-associated, RNA modification, and RNA processing categories) amounts to 30.4% of all SimChrom proteins weight (4.8% of whole-organism proteome weight). However, many proteins from these categories are also localized in cytoplasm, and the major contribution to their cumulative molecular weight likely comes from the cytoplasmic fraction. If the same analysis is performed only for the SimChrom proteins that are uniquely localized in the nucleus (**Supplementary Figure SF4_3A**), the mass fraction of histone goes up to 38% of all the chromatin proteins that are uniquely localized in the nucleus.

Other functional chromatin protein groups (or groups with specific properties) with high values of median abundance and high level of individual protein abundance include HMG A/B/N, histone tail cleavage, histone chaperones, chromatin remodelers and other categories (see Figure 4D). The high mobility group proteins (HMG A/B/N) are the second group after histones ranked by their median abundance. Although grouped together due to historical reasons, they include three separate superfamilies: HMGA (contains AT-hook domains), HMGB (contains DNA binding HMG-box domain), and HMGN (contains nucleosome binding domain). Our analysis suggests that the ratio of HMG proteins to nucleosomes is 1:8, 1:2, 1:3 for HMGA, HMGB, or HMGN proteins, respectively. However, the majority of HMG proteins are not exclusively localized in the nucleus (Figure 4D). The histone tail cleavage proteins are another small group of proteins in our classification with high median abundance in the whole-organism proteome. These enzymes, however, are not exclusively specific for histone cleavage, and likely perform their main functions outside the nucleus by cleaving other proteins. Among histone chaperones the H3-H4 histone chaperone NPM1 and H2A-H2B histone chaperone NCL have the highest abundance, 32% and 13% of H4 abundance, respectively. The most abundant histone variant specific chaperone is ANP32E (specific for H2A.Z-H2B with abundance of 2%). The whole nucleosome chaperone FACT complex consisting of SSRP1 and SUPT16H gene products, has an abundance of around 1%, amounting to one FACT complex per around 50 nucleosomes. Among RNA polymerase subunits POLR2E the common subunit E of RNA polymerases I, II, and III is the most abundant protein (0.58% of histone H4 abundance or around 1 per 90 nucleosomes). The exclusive components of polymerase II (POLR2B, POLR2C, POLR2D, and others) have their abundances in the range of 0.05-0.3%. With a median human gene length of 24kb and nucleosomal repeat length of around 200 bp this gives a lower estimate of one polymerase II per approximately 10 genes. Among genes involved in chromatin remodeling actin encoding genes (*ACTB*, *ACTA1* and actin-like *ACTL6A*) are leading by the abundance of their protein products. While actin is a component of some chromatin remodeling complexes (e.g., SWI/SNF) the major contribution to its abundance clearly comes from its cytoplasmic fraction involved in cytoskeleton formation [28]. The next by abundance are RUVBL1/RUVBL2, members of the family of ATPases associated with diverse cellular activities (the so-called AAA+ proteins), their abundance is around 1.6-1.9%. They form heterohexameric subunits in INO80 remodeller complexes [29], but this is not their exclusive function [30]. The same is likely the case for UCHL5, a putative and optional component of INO80 having relative abundance of 0.6%. The main ATPase of the INO80 family (*INO80* gene) has a dramatically lower abundance (0.006%, suggesting the presence of one INO80 complex per 8680 nucleosomes). The other more abundant chromatin remodeler families include the ISWI family (ATPase subunit SMARCA5 and auxiliary subunits DDX21, SF3B1, MYBBP1A having abundance of 0.7%, 1.6%, 1.2%, 0.6%, respectively), the CHD family (ATPase subunit CHD4 – abundance of 0.6%), the SWI/SNF family (ATPase subunits SMARCA4 or SMARCA2, and axillary subunits SMARCC2, SMARCC1, SMARCE, having abundance of around 0.2 and 0.6, respectively). By the abundance of their catalytic subunits we estimate the presence of one ISWI, CHD, or SWI/SNF remodeler complex per around 70, 80, or 200-300 nucleosomes, respectively.

Among genes involved in formation of SMC complexes the leading are SMC3 and SMC1A with abundance of around 0.8%. These genes are known to form mitotic cohesin complex together with RAD21 (abundance of 0.46%), which has been recently found to perform DNA loop extrusion process in the human genome needed to maintain 3D genome organization and topologically associating domains [31]. Based on SMC3 abundance we estimate the presence of one SMC1-SMC3 complex per around 60 nucleosomes. Among PTM readers the most abundant are the HP1α, HP1β, and HP1γ proteins involved in recognition of methylated H3K9 and formation of heterochromatin (respective abundances are: HP1γ (*CBX3* gene) 3.65%, HP1β (*CBX1* gene) - 1.82% , HP1α (*CBX5* gene) - 1.33%). Among Histone PTM writers the most abundant is PARP1 which PARylates many targets including histones (3.97%). Other nonspecific PTM writers with high abundance are a group of kinases (STK4, PRKDC, PAK2,CDK1) and FBL methyltransferase, which methylates both RNA and histones (particularly at H2AQ104me [32]). Other known histone PTM writers with relatively high abundance include PRMT1 (involved in H4R3me1, H4R3me2 deposition), PRMT5 (involved in H2A, H4R3, H3R8 methylation), NAA50 (involved in H4 acetylation), TGM2 (catalyzing serotonylation and dopaminylation of histone H3) having abundance of 1.5%, 0.7%, 0.87%, 1.67% respectively. The abundance of Polycomb group (PcG) subunits, in particular, PRC1 complex, which monoubiquitinates H2AK119, is around 0.02−0.18%, suggesting one complex per 275 nucleosomes (based on RNF2 and CBX2/4/6/7/8 subunits). Subunits of PRC2 complex which trimethylates H3K27 approximately twice lower: abundance of 0.08%−0.09% of histone H4 for core subunit EZH2 and EED, SUZ12 (suggesting one complex per 550 nucleosomes). The most abundant Histone PTM erasers contain deacetylases: nonspecific to histones HDAC4 (abundance of 2.28%), HDAC2 (abundance of 1.337%), HDAC1 (abundance of 1.108%), and component of remodeling complex NuRD: MTA2 (abundance of

0.597%)), phosphatases (PPP1CC, PPPCA, PPP2CB, PPP5C, abundance of 0.6%−1.2%). The most abundant demethylase is JMJD6 (acts on H3R2me, H4R3me) - abundance of 0.18%.

The groups with least median abundance are those related to “DNA-binding transcription factors” (pioneer TFs, nuclear hormone receptors, housekeeping and non-housekeeping TFs) (note that this SimChrom-SL category excludes this group). Non-housekeeping DNA binding transcription factors have an abundance between 0.00008 and 23.9 ppm, suggesting their expression at minimal levels when averages across all body tissues. The majority of housekeeping transcription factors is also expressed only marginally (median abundance 0.314 ppm). However, the abundance of certain proteins classified as housekeeping TF may reach 91.8 ppm (RURB1 gene, 0.84% of H4) for TF uniquely localized in the nucleus or 3667 ppm for proteins that have multiple localization (ENO1 gene, 33.6% of H4). The DNA-binding transcription factor groups have the largest number of genes in SimChrom-SL (1345 genes in total), however, their contribution to the cumulative weight of chromatin proteins is a rather small 6.3%.

We also performed a similar abundance analysis of chromatin proteins grouped by SimChrom-SL categories for the MS-based experimental datasets, where the abundance values were taken from the respective studies (see **Supplementary Figure SF4_4**). The mass fraction of SimChrom proteins in experimental chromatomes/nucleosomes was between 33% and 81% (Alabert et al., 2014 - 65%; Kustatscher et al., 2014 - 81%; Itzhak et al., 2016 - 77%; Ginno et al., 2018 - 33%; Shi et al., 2021 - 78%; Ugur et al., 2023 - 43%). This reflects the fact that among the proteins detected in MS-based studies, SimChrom proteins constitute only a fraction (see **Supplementary Figure SF2_5A**). In Ginno et al., 2018 and Ugur et al., 2023 the total mass fraction was dominated by these non-SimChrom proteins. In Shi et al., 2021 and Alabert et al., 2014 the significant mass fraction was provided by one group of proteins (e.g., by "Histones" in Shi et al., 2021 and "DNA-acting enzymes" in Alabert et al., 2014). Itzhak et al., 2016 dataset (in which proteins from predominantly nuclear fraction were estimated in copy number per cell) showed a more balanced distribution of mass fraction between different SimChrom-SL categories: Nuclear RNA-binding proteins (26%), Histones (19%), Chromatin remodelers (6.4%), and Histone PTM writers (3.4%). Taken together our analysis revealed a high level of variation and potential experimental biases in quantitative proteomics data provided by MS-based studies.

##### 3.2. Detailed analysis of the physico-chemical properties and amino acid composition

This section supplements and expands section 3.3.2. Physico-chemical properties and amino acid composition in the main text.

Using the constructed datasets of chromatin, and uniquely localized nuclear and cytoplasmic proteins (SimChrom, NULOC_CS_UL, CYTLOC_CS_UL) we analyzed the physico-chemical properties of chromatin/nuclear proteins, their distinction from cytoplasmic proteins and peculiarities of chromatin proteins belonging to different classes of the SimChrom classification. Figure 5A,B summarizes the results, details are provided in Figure 5C-L and **Supplementary Figures SF5_1 - SF5_5**.

The length distribution of chromatin proteins is positively skewed with respect to that of cytoplasmic proteins but only in the range of protein length less than around 800 amino acids (Figure 5C). The same picture is seen for nuclear proteins (**Supplementary Figures SF5_1A**). The median length of cytoplasmic proteins is 464, while 496 for SimChrom and 500 for nuclear proteins (**Supplementary Figure SF5_1A** for nuclear vs cytoplasm comparison). Hence the median length of chromatin proteins is 7% longer than that of cytoplasmic proteins. The fraction of relatively short proteins (less than 200 aa) is 1.6 times smaller for chromatin proteins than for cytoplasmic ones (13% vs 8%), suggesting that smaller proteins are on average underrepresented in chromatin with respect to cytoplasm. The longest chromatin proteins in our dataset were those from following SimChrom categories: 'Preribosome associated': (MDN1 - 5596 aa); 'Histone PTM writers' (Histone methyltransferases: KMT2D (5537 aa), KMT2C (4911 aa), KMT2A (3969 aa), E3 ubiquitin-protein ligase HUWE1 (4374 aa), kinase PRKDC (4128 aa)); 'TFs': TRRAP (3859 aa), ZFHX3 (3703 aa), ZFHX4 (3567 aa), ZNF292 (2723 aa), HIVEP1 (2718 aa); 'Chromatin remodeler' subunits: SRCAP (3230 aa), EP400 (3159 aa), CHD7 (2997 aa), CHD9 (2897 aa), CHD6 (2715 aa), while the shortest from 'HMG', 'Histones' (90-130 aa) and 'RNA polymerase' subunits: POLR2K (58 aa), POLR2L (67 aa).

The fraction of intrinsically disordered regions in chromatin and nuclear proteins (as calculated by AlphaFold2 SASA with 20-residue smoothing, see Methods Section 2.4) and its distribution was significantly different from the one for the cytoplasmic ones (see Figure 5D). 46% of chromatin and 51% of nuclear proteins had an IDR fraction of more than one half, while for cytoplasmic proteins the value was only 23%. The median IDR fractions also differed considerably (46%, 51%, 23%, correspondingly). Further analysis showed that the increase of IDR fraction was both due to the increase in average IDR length and the number of IDRs (see **Figure 5E,F**). The fraction of proteins with two, three or four IDRs was significantly higher for chromatin/nuclear proteins than for cytoplasmic ones (e.g., fraction of proteins with two IDRs was 38%, 41%, 27% for nuclear, chromatin and cytoplasmic proteins, respectively) mostly at the expense of a lower fraction of proteins without or with one IDR (e.g., fraction of proteins with IDR was 20%, 18%, 33%, respectively). A similar tendency was observed for the increase in the number of non-IDRs in chromatin/nuclear proteins with respect to cytoplasmic ones, albeit to a lesser extent (see Figure 5F).

We performed a detailed analysis of protein length and IDR fraction for different categories of chromatin proteins (see Figure 5G). The protein categories with the shortest median length are histones (130 aa), HMG proteins (109 aa), and RNA polymerase subunits (220 aa). The longest ones are SMC complex subunits (1096 aa), Chromatin remodelers (835 aa), Histone PTM writers (766 aa), Histone PTM erasers (747 aa), DNA-acting enzymes (655 aa). The categories with the largest variation in size (ratio of 5% and 95%-percentiles) include Histone PTM readers (1741), General TFs (1746), Histone chaperones (1768), DNA-acting enzymes (1810), DNA (de)methylation (1830), Centromere-associated (2013), Chromatin remodelers (2314), Histone PTM writers (2988). The categories with the smallest variation in protein length (calculated for protein categories with more than 15 entries) include Histones (144), Pioneer TFs (321), Nuclear hormone receptors (598), Non-Housekeeping DNA-binding TFs (766), RNA modification (768), Methylated DNA binding (856), Housekeeping DNA-binding TFs (874), Other families with HMG-box domain (943).

The median IDR fraction was smaller than that of cytoplasmic proteins (13%) for proteins in the following categories: RNA polymerase subunits (0%), Histone tail cleavage (3%), and Preribosome-associated (10%). Other groups with relatively small fraction of IDR included: DNA-acting enzymes (16%), RNA modification (16%), DNA de_methylation (17%), SMC complex subunits (18%), DNA repair (19%). The categories with the highest fraction of IDRs are HMG (99%, 7 out of 11 proteins are completely disordered), Other families with HMG-box domain (74%), Pioneer TFs (73%), Methylated DNA binding (62%). The analysis allowed us to identify the chromatin categories contributing most to the high IDR fraction proteins observed in the distribution in Figure 5G. The large categories that contain many proteins with high IDR fraction are those belonging to transcription factors, especially the “Non-Housekeeping DNA-binding TFs” (contributes 453 proteins with IDR fraction of more than 50%). The distribution of IDR fraction for chromatin proteins without the transcription factor groups (“Non-Housekeeping DNA-binding TFs” and "Housekeeping DNA-binding TFs") resembles that of the cytoplasmic ones much better (see **Supplementary Figure SF5_1B**). Overall our analysis suggests that chromatin and nuclear proteins are significantly enriched in the number and length of IDRs, which likely are important for fuzzy interactions happening in chromatin including liquid-liquid phase separation behavior. This enrichment is particularly evident for non-housekeeping transcription factors.

We also performed the analysis of IDR fraction distribution among the chromatin/nuclear protein sets identified in experimental MS-based studies discussed in Results Section 3.1. The Kustatscher et al., 2014; Itzhak et al., 2016; and Ugur et al., 2023 datasets (that were previously shown to include a substantial fraction of low abundant proteins) had a considerably higher proportion of proteins with high IDR fraction. This can be explained by the presence of a larger number of non-housekeeping TF in these datasets (see **Supplementary Figure SF5_1C**, **Supplementary Figure SF2_7C**).

It is generally assumed that chromatin proteins are on average positively charged to compensate for the negative charge of the genomic DNA. Using the protein datasets compiled above and the protein abundance data from PaxDB (see Results Section 3.3.1) we set out to analyze the distribution of chromatin/nuclear proteins with respect to their overall electrostatic charge. To this end we used three metrics (see Figure 5H). In the first approach we classified the proteins as negatively/positively charged or nearly neutral (see Methods Section 2.4) and calculated the number of protein entries in our datasets belonging to these classes. According to this approach positively charged proteins dominate among nuclear and chromatin proteins, while the negatively charged proteins dominate among the cytoplasmic ones. The dominant charge group in each case consists of around more than half of protein entries (54-58%), while the opposite one consists of 38-43% of protein entries. This metric, however, does not account for protein abundance or total protein charge. In the second approach we adjusted the first metric by protein abundance, while in the third approach we additionally took into account the overall net charge of every protein, resulting in the analysis of the cumulative charge conferred by positively or negatively charged proteins (see Figure 5H). Interestingly, only among the uniquely localized nuclear proteins the results were significantly affected by these two adjustments. The amount of positively charged proteins increased to 82% once the protein abundance was taken into account, and the overall net positive charge coffered by these proteins was 89%. This is mainly due to the high abundance and high net positive charge of histone proteins present in NULOC_CS_UL dataset. Surprisingly, for the SimChrom dataset the distribution of positively/negatively charged fractions did not change much as the result of these adjustments (43% vs 37% vs 39%). A detailed analysis of cumulative charges conferred by proteins from different SimChrom categories showed that significant cumulative negative net charge is contributed by proteins involved in RNA processing, Chromatin remodeling, Histone chaperones, and DNA-acting enzymes (see **Supplementary Figure SF5_1D**, left). These categories, however, have many proteins that are localized both in the nucleus and the cytoplasm, which reconciles these results with the charge distribution analysis of NULOC_CS_UL dataset discussed above. The same analysis of SimChrom proteins that are uniquely localized in the nucleus revealed a prevalence of positive charge by histone proteins (see **Supplementary Figure SF5_1D**, right). To further elucidate the nature of negatively charged proteins in chromatin we analyzed the fraction of entries belonging to negatively/positively charged proteins in each SimChrom category (**Supplementary Figure SF5_1E**). Among the categories having the most number of negatively charged proteins are Histone chaperones (84%), RNA polymerases (76%), Histone PTM erasers (71%). This surprising presence of many negatively charged proteins in these categories is likely explained by their preferential association not with DNA, but rather with positively charged histones.

To further elucidate the peculiarities of charge structure in chromatin proteins we analyzed the average charge profiles of protein N- and C-terminal tails. The median length of N-/C-terminal tails was 51/47, 72/84, and 67/79 for CYTLOC_CS_UL, NULOC_CS_UL, and SimChrom (see

Supplementary Figure SF5_2A,B). Only for the N-terminal tail there was a clear difference in the average charge profiles between cytoplasmic proteins and nuclear/chromatin ones (see Supplementary Figure SF5_2C,D, Figure 5I). The average charge of the N-terminal tails (less than 80 amino acids in length) was -16 for SimChrom proteins, -6 for nuclear proteins, and +13 for cytoplasmic ones (see Figure 5I). The analysis of the N- and C-terminal tail charge for different SimChrom categories revealed that they varied between the different categories (see Supplementary Figure SF5_2E,F). Histones had the most positively charged tails, while histone chaperones and HMGs had the most negatively charged ones. Transcription factors on average also had negatively charged protein tails. This is an interesting fact because most transcription factors are positively charged (Supplementary Figure SF5_1E).

We next set to analyze in detail the amino acid composition of chromatin/nuclear proteins with respect to cytoplasmic ones and the variability of amino acids composition between different groups of chromatin proteins (see Figure 5J,K,L, Supplementary Figure SF5_3, SF5_4, SF5_5, Supplementary Table ST13). To this end we first used the UMAP nonlinear dimensionality reduction technique to see if significant variations between chromatin proteins can be identified in the space of their amino acid composition. The resulting 2D projections onto the main UMAP components revealed that (1) chromatin and cytoplasmic proteins occupied overlapping domains on the 2D map, but with a visible shift between their centers, suggesting there is an overall difference in the average physico-chemical properties, (2) some chromatin proteins manifested a significant difference from others (see cluster 1 and cluster 2 in Figure 5K,L). Further analysis revealed that in the 2D UMAP map transcription factors, containing zinc finger domains and homedomains formed distinct clusters (see Supplementary Figure SF5_3A,B). The most distinct group (cluster 1) was **almost exclusively** (415 out of 422) composed of zinc-finger containing DNA-binding transcription factors (240 housekeeping and 175 non-housekeeping) with the median number of zinc-finger domains (ZFD) of around 10 (Supplementary Figure SF5_3C). Zinc-finger containing DNA-binding transcription factors were also present in cluster 2, but the median number of zinc-finger domains (ZFD) in that cluster was only three, hence containing a lower proportion of amino acids specific to ZFD (Supplementary Figure SF5_3C,D). ZFD are enriched in histidine and cysteine, which were among the top four mostly enriched amino acids in chromatin proteins (see Figure 5J and discussion below). Other protein groups that occupied distinct positions on the UMAP map, included (1) histones, (2) serine/arginine-rich splicing factors (SRSFs) (enriched in serine and arginine), and (3) reverse transcriptases of endogenous retroviruses (ERVs) (enriched in isoleucine and threonine) (see Figure 5K, Supplementary Figure SF5_3D).

The analysis of amino acids composition revealed that among the top four enriched amino acids in chromatin proteins were serine, cysteine, proline, histidine (Figure 5J, Supplementary Figure SF5_5B). Their enrichment values are in the range 1.13-1.18. By classes of amino acids chromatin/nuclear proteins are mostly enriched in polar (N, Q, T, C, G, P), small (P, G, A, S), and positive (K, R) amino acids (**Supplementary Figure SF5_4A**, **Supplementary Figure SF5_5B**). It is important to note that such an analysis should be taken with a grain of salt, because the enrichment of certain amino acids may vary across different categories of chromatin proteins, and the categories with high number of protein entries have higher contribution to the overall average. To elucidate this variability we have also performed enrichment analysis for proteins in major functional SimChrom categories (see **Supplementary Figure SF5_4**).

Serine and proline are among amino acids that are relatively abundant in proteins (abundance of around 7-8% and 5-6% in chromatin and cytoplasmic proteins, respectively, **Supplementary Table ST13**). The total enrichment of serine in chromatin proteins is attributed simultaneously due to its enrichment in the IDR, non-IDR regions (relative to IDR and non-IDR regions of cytoplasmic proteins), and more importantly due to higher proportion of IDR regions in chromatin proteins (46% vs 23%) that in turn have a considerably higher proportion of serine than non-IDRs (**Supplementary Table ST13**). The slight enrichment of serine in IDRs was observed almost across all SimChrom categories (**Supplementary Figure SF5_4E**). In non-IDR regions serine showed both enrichment and depletion in certain categories, and the overall enrichment was driven mainly by transcription factors due to the large number of proteins in these categories. The total enrichment of proline in chromatin proteins is attributed due to its enrichment in IDRs (relative to IDRs of cytoplasmic proteins and more importantly due to higher proportion of IDR regions in chromatin proteins (proline is the most enriched amino acid in IDRs of both chromatin and cytoplasmic proteins versus the non-IDRs, fold enrichment 1.8-2.1, **Supplementary Table ST13**). The overall enrichment of proline in non-IDRs was close to one and statistically not significant. Certain small groups, such as histones and HMG proteins showed considerable deviations in proline content in their non-IDRs (**Supplementary Figure SF5_4H**). The enrichment of proline in IDRs was observed in many SimChrom categories (**Supplementary Figure SF5_4E**). In certain categories, such as Non-Housekeeping transcription factors and pioneer TFs enrichment was high (1.37 and 1.55 fold, respectively). Surprisingly, the enrichment of proline in IDRs of housekeeping TF was depleted (FE of 0.92). Suggesting that while there is still a considerable fraction of prolines in IDRs of housekeeping TF, this fraction is significantly lower than in IDRs of non-house keeping TF (7 % and 10.4 % median fractions, respectively).

Cysteine and histidine are among amino acids that have a relatively low abundance in proteins (abundance of around 1-2.5%, **Supplementary Figure SF5_5B**). The total enrichment of cysteine and histidine is mainly driven by the prevalence of zinc fingers containing transcription factors (**Supplementary Figure SF5_4B**). The exclusion of these proteins from analysis resulted in the disappearance of any statistically significant enrichment **Supplementary Figure SF5_5A**).

Interestingly, the enrichment of positive amino acids is only statistically significant for lysine, but not for arginine, and the enrichment is relatively moderate (1.03 in chromatin) (**Supplementary**

Figure SF5_4A). Lysines are enriched in IDRs and non-IDRs of chromatin proteins, while arginines are depleted in IDRs and enriched in non-IDRs (when compared with IDRs and non-IDRs of cytoplasmic proteins) (Supplementary Figure SF5_5B). The overall higher positive charge of chromatin proteins stems also from the depletion of negatively charged amino acids in their sequence. The depletion of aspartate in chromatin/nuclear proteins is statistically significant (fold enrichment is around 0.9), while the depletion of glutamate is statistically non-significant (Supplementary Figure SF5_4A). This suggests that the increased positive charge of chromatin nuclear proteins has its main contributions in the depletion of aspartate, and moderate enrichment of lysine.

Within the IDR regions of chromatin proteins tyrosine and asparagine were also significantly enriched (FE of 1.23 and 1.17 versus the IDRs of cytoplasmic proteins, respectively), mainly due to the contribution of transcription factors (Supplementary Figure SF5_4F).

Among the most relatively depleted amino acids in chromatin/nucleus are tryptophan and hydrophobic/aliphatic amino acids like valine, isoleucine, leucine, and methionine (Supplementary Figure SF5_4A). Tryptophan is the rarest amino acid in proteins (around 1%). Certain categories like histone and HMG proteins lack it completely (Supplementary Figure SF5_4C). It is depleted in almost all chromatin categories, except for a few small categories such as DNA (de)methylation, histone tail cleavage, and histone modification, where the enrichment comes from non-IDRs (Supplementary Figure SF5_4I). Hydrophobic/aliphatic amino acids are depleted in IDRs vs non-IDRs of proteins and hence the large proportion of IDRs in chromatin proteins accounts for a lower fraction of these amino acids in chromatin proteins (Supplementary Figure SF5_4F,I).

The most enriched amino acids in the uniquely localized nuclear proteins were the same except for cysteine (this difference may be traced to the diminished number of transcription factors enriched in cysteines in the NULOC_CS_UL dataset, apparently because of their multiple localization, see Supplementary Figure SF5_4A, Supplementary Figure SF4_3B,).

##### 3.3. Detailed analysis of the domain composition of chromatin proteins and identification of new structural domains

This section supplements and expands section 3.3.3. Domain composition of chromatin proteins and identification of new structural domains in the main text.

Next we set out to systematically analyze the available data on structural characterization, domain annotation and domain composition of chromatin proteins. We specifically explored the structurally uncharacterized portion of the chromatome ("dark" proteome) and identified potential new structural domains that are predicted by AI-based protein structure prediction tools (see Figure 6A).

Historically, protein domains are loosely defined as evolutionary conserved units with similarities at functional, structural and/or sequence levels [33]. Domains may represent single proteins or exist in a variety of various sequence contexts. Sequences of related individual protein domains may be grouped and aligned to produce domain models. Domain models are catalogued and annotated by a number of resources/databases such as PFAM [34], CDD [35], CATH [36], InterPro [37], and may be further grouped into superfamilies, clans, folds, etc [38,39]. Domain models are usually defined through multiple sequence alignments (MSA) and corresponding hidden Markov models (HMM). In structure-based approaches (e.g. CATH/Gene3D database) domain superfamilies are assigned through grouping and alignment of available experimental 3D structures. The ultimate experimental structural characterization of chromatin proteins is available in the PDB database, however, recent progress in protein structure prediction spurred by AlphaFold resulted in new approaches to the structural characterization and discovery of new structural domains (*e.g.*, as implemented in the TED database) [40] (Figure 6A). Structure prediction algorithms combined with structure similarity search algorithms, such as FoldSeek [41], now allow to find remote homologs and assign individual domains to their respective superfamilies.

**Figure 6B** shows the fraction of the aggregate number of amino acids in all human chromatin proteins (referred below to as "aggregate chromatome sequence", or ACS) which are structurally characterized or have domain annotations in different databases. According to AlphaFold approximately one half of the ACS (47%) is predicted to be intrinsically disordered, or to become ordered within protein-protein complexes (the structurally uncharacterizable "dark" chromatome) (see Methods), and the rest as having distinct 3D structure. Direct experimental structural data in PDB is available for only one fourth of the ACS (20% of ACS are simultaneously considered ordered by AFDB and available in PDB). Hence, we envision that at least one third (34%) of the aggregate human chromatome sequence is amenable to characterization with structural biology methods but has not yet been characterized (constitutes the potentially structurally characterizable "dark" chromatome). The Pfam database (the largest sequence-based database of protein domains and protein families to date) has annotations for around 39% of the aggregate human chromatome sequence. The CATH database, which focuses on identifying and annotating structural domains, annotates 25% of ACS, while the automated AlphaFold-driven TED resource finds structural domains in 35% of ACS. The difference between the fraction of ACS annotated by TED and that considered ordered by AFDB was traced to at least several facts: 1) AlphaFold is known to be biased to predict long solitary alpha-helices which are not considered domains by algorithms that identify structural domains, 2) the TED algorithm frequently fails to annotate repetitive regions that contain visually identifiable secondary structure elements within large multidomain proteins, 3) we identified non-IDRs as regions no less than 4 amino acids whereas median length of TED domain in human proteins were 108 aa. A caveat that has to be kept in mind, is that current automated analysis using AlphaFold is based only on predictions for single chain proteins, while in reality chromatin proteins engage in many intermolecular interactions. To some extent Pfam and CATH/TED are complimentary (see Figure 6B). In addition to 39% of ACS annotated by Pfam, TED annotates additionally 13% of ACS, and CATH adds annotations to 3% of ACS on top of it (yielding a combined annotation coverage of 55% by these three resources).

Next we analyzed the structural characterization of the aggregate human chromatome sequence from the point of view of structural domains present in chromatin proteins (as identified by the most comprehensive TED database, which automatically detects structural domains) (see Figure 6C). Chromatin proteins contain in total 6246 individual TED domains. Using FoldSeek and combination of FoldSeek and CATH resources (see Methods) we matched these domains to the structurally related domains in PDB or CATH superfamilies. The remaining domains were analyzed for the presence of previously uncharacterized structural folds/superfamilies and potential functional roles of these domains. Among the 6246 predicted structural domains constituting human chromatin proteins, 34% had exact matches in PDB structures (100% sequence identity, see Methods), 56% matched PDB structures of homologues with different levels of sequence identity (from 99% to 5%, for details see Figure 6C). The majority of these homologous domains were in fact different paralogous sequences found within human genes (even for domains with sequence identity of 35-50% the fraction of human sequences among the matches was 51%), for matches with sequence identity above 35% the second largest contribution came from structures of mammalian homologues, for matches with sequence identity below 35% significant contributions were from structures derived from proteins of fungi, protostomia and bacteria (see **Supplementary Figures SF6_1A** for details). Additionally, 6% of TED domains that lacked direct hits among the PDB structures were mapped to protein structural superfamilies in the CATH database (the information about potential sequence variation in each homologous superfamily collected in CATH database combined with AlphaFold structural predictions allowed to identify more distant structurally characterized homologues). The remaining 4% (241 TED domains) represented domains that could not be matched to any known protein structure or protein structure superfamily, potentially representing new types of structural superfamilies of even protein folds. These domains are presented in **Supplementary Table ST14** (see also **Interactive Table 3** at https://simchrom.intbio.org/#novel_structural_domains) and ranked via their structural complexity by the number of their secondary structure elements. Among these domains, 123 domains have annotations in Pfam or other domain annotation databases present in InterPro, leaving 118 domains that are completely without annotations. The latter domains belong to 106 chromatin proteins, which may be considered as perspective new targets for experimental studies of their function and structure. Among such proteins are, for example, a protein encoded by the *GTF3C1* gene, a General transcription factor

3C polypeptide 1 (it has a previously unannotated and uncharacterized structural domain with length of 233 amino acids) (see detailed characterization in **Supplementary Figure SF6_2A**). Another instructive example is the globular domain of the testis specific linker histone H1.7 (product of *H1-7* gene, see **Supplementary Figure SF6_2B**). Despite the considerable amount of studies dedicated to the elucidation of the structure of H1-linker histones [42], the H1.7 histone variant (previously, named HANP1/H1T2) has a quite different sequence resulting in a predicted structure that has a different topology than other known H1 histones (the "wing" of the globular domain consists of three beta-sheets rather than two). The relation of this domain to the H1 histone family cannot be identified with conventional sequence analysis methods (such as those implemented in the Pfam database), however, it should be noted that new deep-learning-based annotation approaches (such as Pfam-N) are able to annotate it (see **Supplementary Figure SF6_2B**).

Next we predicted GO molecular functions and biological processes for mentioned above 118 not-annotated chromatin protein domains using DeepFRI [43], a Graph Convolutional Network for predicting protein functions by leveraging sequence features extracted from a protein language model and protein structures, see Methods Section 2.5. The top-7 common GO MF terms: ion binding, organic cyclic compound binding, heterocyclic compound binding, protein binding, cation binding, metal ion binding, nucleic acid binding. Top-10 GO BP terms: organic substance metabolic process, primary metabolic process, cellular metabolic process, nitrogen compound metabolic process, macromolecule metabolic process, organonitrogen compound metabolic process, regulation of cellular process, cellular macromolecule metabolic process, cellular nitrogen compound metabolic process, cellular response to stimulus. 9 out of 118 TED domains lacked the predicted GO molecular function by DeepFRI: two of them were in members of the regulatory factor X (RFX) family of transcription factors (encoded by genes *RFX1, RFX5*).

Many chromatin proteins contain similar, evolutionary related individual protein domains whose kinship may be identified by matching them to the same Pfam domain sequence models. Hence, we used the Pfam domain annotation to characterize the diversity of protein domains found in chromatin proteins and typical domain composition thereof. In total 1753 different domain types (sequence models) were identified in chromatin proteins (Figure 6D). Next we analyzed the structural information available for these models. 76% of domain models had at least one individual domain among chromatin proteins that could be matched to a PDB structure using FoldSeek (*bona fide* structural domain in Figure 6D). To characterize the comprehensiveness of the structural characterization of each domain model we estimated the median sequence identity between all individual domains in chromatin proteins belonging to the said domain model and their best matches in PDB found via FoldSeek (see Methods Section 2.5, Figure 6D). 42% of domain models were considered fully characterized, *i.e.* every individual domain in chromatin proteins belonging to these models can be found in PDB, 34% of domain models are partially characterized. 14% of Pfam domains were not matched by FoldSeek to PDB structures, but could be still identified in PDB via sequence search methods – these represented IDR regions, repeats, etc. 3% (55 domain models) did not match any PDB structure but could be matched to structural domains predicted by AlphaFold and found in the TED database. These represent prospective targets for validation with structural biology methods and further investigation of their interactions. For instance, among these domains are domains, potentially associated with chromatin remodeling (SANTA, zf-C3Hc3H), histone PTM writing (DUF7030, COMPASS-Shg1), zinc fingers (zf_CCCH_4, zf-LITAF-like, zf-WIZ, SWIM) etc. 7% of Pfam domain models currently have no structural information that can be assigned either through the PDB or TED databases.

We next analyzed the diversity of Pfam domains in various SimChrom-SL protein categories (Figure 6E, subpanels 1,2) and the domain content of individual proteins belonging to these categories (Figure 6E, subpanels 3-5). Pfam identified 11147 individual domains in chromatin proteins belonging to 1753 domain types (Pfam models); only 70 chromatin proteins had no domain annotation at all. For the distribution of the total number of Pfam models and distinct Pfam models identified in chromatin proteins, see **Supplementary Figure SF6_1B**, **Supplementary Figure SF6_1C**. Expectedly, large SimChrom-SL categories consisting of more than one hundred proteins harbored the largest number of different domain types (*e.g.*, DNA-acting enzymes, histone PTM writers, transcription factors, etc.), while the smaller categories had less (see Figure 6E, subpanel 1). Although Pfam may not be comprehensive in its annotation, we estimated the number of distinct domain types per protein in each category (relative domain diversity, Figure 6E, subpanel 2). The average domain diversity was around one for all categories. The categories with the considerably lower domain diversity were transcription factors categories (their variability relies on different combinations of zinc-finger domains that are described through only a few Pfam domain models), histones (their functional variability is often conferred by only small changes in the sequence), and HMG-constainting proteins (this is a very small group of proteins with only nine proteins and three corresponding Pfam models). The median number of Pfam domains in human chromatin proteins was two (which corresponds to the structure based domain analysis presented above). Certain chromatin protein categories had a higher median number of domains, including Housekeeping TF, histone PTM writers and readers (Figure 6E, subpanel 3). Interestingly the median number of domains for Non-housekeeping TF was two (they more often rely on single homeodomains than on cassettes of zinc-finger domains), although the group is diverse and proteins with as many as 32 domains were present. This, however, is again explained by the large number of zinc-finger domains that may be present in such proteins (see **Supplementary Figure SF6_1D**). The presence of additional domains in PTM writers and readers may be hypothesized to have evolved due to the functional necessity for multivalent binding to different chromatin structures (see below for a detailed analysis). Some categories mostly consist of single domain proteins, such as Centromere-associated, DNA repair, Regulation of transcription, RNA polymerases (but this group also includes proteins with a maximum of 42 domains), DNA recombination, RNA modification, Histones, HMG_A/B/N, etc. The analysis of the number of distinct different domain types present in chromatin protein categories corroborates the above mentioned analysis (Figure 6E, subpanel 4). Proteins from PTM writers group have the median number of three distinct domain types, while all other categories have less. Still many chromatin proteins harbor many distinct domain types, DNA-acting enzymes, histone PTM writers, chaperones, remodelers, transcription factors with as much as 8-9 distinct domains are present (see **Supplementary Table ST15**). There are 118 chromatin proteins harboring at least five different domain types (see **Supplementary Figure SF6_1C**). This highlights the multivalency of protein interactions in chromatin, keeping in mind that many proteins further form protein-protein complexes increasing their interaction potential. The average individual domain length in chromatin proteins is around 65 amino acids (the median is 28 aa), however, this number is biased by the presence of many zinc-finger domains (around 22 aa in length). Subpanel 5 in Figure 6E gives a more balanced view for each SimChrom category. For the majority of protein groups the median domain length is around 100 amino acids (mean is 137, median is 134).

The birds-eye view of the most frequently occurring Pfam domains' in various functional SimChrom-SL categories is presented in Figure 7. The data is presented for domains that occur in at least five chromatin proteins and in at least 10% of proteins in a category (the threshold for data point depiction is 5%). The comprehensive interactive analysis figure with the ability to alter these thresholds and switch between SimChrom and SimChrom-SL classifications systems is available in **Interactive Figure 4** (https://simchrom.intbio.org/#domain_composition). In Figure 7 the following categories and their respective domains can be grouped revealing their partially shared domain composition: 1) the categories containing transcription factors and their zinc finger, homeodomains and KRAB domains form the most frequently occurring entities, 2) some chromatin regulators, such as PTM writers, readers, erasers and chromatin remodelers together with their Chromo, Bromodomain, PHD.

##### 3.4. Detailed analysis of the multivalent interactions in chromatin proteins

This section supplements and expands section 3.3.4. Multivalent interactions in chromatin protein in the main text.

The presence of multiple domains (belonging to the same or different domain models) in chromatin proteins is a known feature contributing to their ability to engage in multivalent interactions (Figure 8A) [44]. Below we present the analysis of such domains engaged in multi-valent interactions (referred to as EMVI-domains hereafter) that are found in chromatin/epigenetics regulator proteins (see Figure 3 for definition of this group). To limit our analysis to a manageable set of EMVI-domains, we selected those that were found in multiple copies or in combination with another Pfam domain in at least three chromatin regulator proteins (94 Pfam domains in total), and from those we selected 59 domains that we were able to manually classify based on the information currently available in the literature according to their functional binding modes. The following *functional groups* of domains were used: histone methylation/acetylation/phosphorylation, chromatin remodeling, histone binding, DNA binding, DNA methylation, protein dimerization/oligomerization, PPI, RNA binding. Histone post-translational modifications were further subdivided into readers, writers and erasers *functional subgroups* (see Figure 8C, **Interactive Figure 5 (https://simchrom.intbio.org/#domain_co-occurrence),** and **Supplementary Table ST16** for the list of domains and their detailed classification). We consider this subset of chromatin proteins' domains as representative to illustrate the concept of multivalency in chromatin regulators interactions, since the selected domains are extensively characterized and their functions are known. A comprehensive analysis would require characterization of all 409 Pfam domain models that are found in combination with other models or in multiple copies in at least one chromatin regulator protein. As a compromise our online database includes the analysis of 163 Pfam domain models that are present in at least two chromatin regulator proteins (see https://simchrom.intbio.org/#domain_co-occurrence).

First, we analyzed the co-occurrence of selected EMVI-domains in all chromatin proteins. There were in total 922 chromatin proteins (306 with the exclusion of transcription factors) that had more than one selected EMVI-domain (including multicopies). The conditional probability of finding a corresponding domain A in a chromatin protein given that another domain B is already present was estimated and is presented in Figure 8C (columns and rows correspond to domains A and B, respectively). The **Interactive figure 5** is available at https://simchrom.intbio.org/#domain_co-occurrence (also includes unclassified potential EMVI-domains found in at least two chromatin regulator proteins). The matrix in Figure 8C allows to trace the interplay between different domains employed in architectures of chromatin proteins. The largest groups of domains in Figure 8C are those involved in histone methylation and DNA binding, suggesting that these mechanisms are the most represented and employed in chromatin functioning regulation.

There were 49 cases where association between the presence of various domains in chromatin proteins was 100% (red squares in Figure 8C), among them for 18 cases (9 domain pairs) the association was reciprocal (i.e. P(A|B) = P(B|A)). In certain cases this exclusive association between domains may be traced to the fact that they form a larger structural complex with direct structural interactions between the domains as judged by the visual inspection of AlphaFold based predictions (MOZ_SAS and zf-MYST, ADD_DNMT3 and DNMT3_ADD_GATA1-like). In other cases the association is likely due to functional reasons, in our analysis in the majority of cases such domains were confined to the histone methylation readers subgroup (KDM3B_Tudor and PWWP_KDM3B; C5HCH, NSD_PHD, PHD-1st_NSD, and PHDvar_NSD found in Histone-lysine N-methyltransferases).

Among the EMVI Pfam domains that co-occur with the most number of other different Pfam domains in chromatin regulator proteins is the histone methylation/acetylation domains: PHD domain (45 other domains), Bromodomain (38), SET (40) and PWWP (28) and chromatin remodeling Helicase_C (33) and SNF2-rel_dom (31).

The diagonal elements in Figure 8C show that certain domains tend to be present in multiple copies, particularly often, MBT, WD40 and zinc finger (zf-C2H2) domains. However, all these are special cases of short repeat domains, where multiple copies are needed to form one functional unit. Not so often, but in a considerable number of proteins PHD and Bromodomain may be found in multiple copies.

Another more general view of multivalent interactions in chromatin proteins may be obtained if we trace the relationships within or between different functional groups of domains. One can see that domains from the same functional group (*e.g.*, histone methylation) tend to co-occur (Figure 8C) in chromatin proteins and may also be present in multiple instances in proteins (**Supplementary Figure SF8_1B**). For example, there may be up to nine histone methylation associated domains in chromatin proteins (**Supplementary Figure SF8_1B**). If a chromatin protein has a domain involved in histone methylation (either writing, reading or erasing) there is an estimated 38% chance that there will be another different functional domain from this group of domains (**Supplementary Table ST17**, **Supplementary Figure SF8_1C**). For acetylation this estimated probability is 31%, for phosphorylation 18%. Note, that domain categorization is not trivial. For example, WD40 is a repeat that folds into a higher order structure (median number in chromatin proteins are four). They can recognize both unmodified histone tails and methylated regions [45] and its classification may affect this part of study.

The associations between the occurrence of domains from different functional groups can also be observed. This can be seen in Figure 8C, and the upset plot in Figure 8D (see also **Supplementary Figure SF8_2A**). One can see that domains involved in histone methylation (one of the most abundant group by the number of Pfam models and the number of chromatin proteins) may be in a considerable number of chromatin proteins combined with other EMVI-domains (particularly DNA binding domains and histone acetylation), association association with with histone binding domains, chromatin remodeling, DNA methylation, (di/oligo)-merization and RNA binding domains was also observed, see **Supplementary Figure SF8_2C**. The same can be said about domains involved in histone acetylation, although in a somewhat smaller number of cases, and with exclusion of their combination with dimerization domains. Notably, domains involved in histone phosphorylation were not found in combination with domains from other functional groups in our analysis. This may reflect an evolutionary strategy whereby combinations of histone methylation and acetylation evolved to delicately regulate gene expression at the epigenetic level, while phosphorylation remained as a more general mechanism affecting a broad number proteins and pathways in the cell. DNA binding domains are found to be associated with methylation, acetylation, dimerization, and chromatin remodeling domains although the relative number of proteins harboring combinations of such domains is small compared to the number of protein (mainly transcription factor) that have DNA binding domains in their sequence. Domains associated with catalytic subunits chromatin remodeling complexes in certain cases are found together with histone acetylation, methylation or DNA binding domains. A more detailed analysis reveals that one "reader" domains for acetylation and methylation are found in these proteins.

For a more comprehensive view of multivalent interactions it is reasonable to (1) analyze not only pairwise co-occurence of different functional domains, but simultaneous co-occurrence of domains from several functional groups in one proteins, (2) extend the analysis to complexes of chromatin proteins. The results of such analysis are presented in Figure 8D (see Methods Section 2.1.4 for our selection of 513 protein complexes where all proteins are chromatin proteins from Complex Portal). In our analysis at the level of individual proteins, proteins harbored domains only from up to three functional groups. Particularly, DNA binding domains may be combined with (di/oligo)merization, chromatin remodeling, histone methylation, methylation and acetylation or histone acetylation and chromatin remodeling domains. The formation of chromatin protein complexes considerably affects the available combinations of functional domains. Among the 513 analyzed protein complexes, 181 complexes contained domains EMVI-domains from the analyzed functional domain groups, 101 complexes harboured more than one domain, 80 complexes harbored domains from different functional groups. From these 80 complexes the majority were various chromatin remodeling complexes (53 complexes), others representatives included acetyltransferase complexes (13), deacetylase (4), and DNA-methylation (2).

One can see that the largest number of analyzed complexes (22) simultaneously contained domains from four functional groups (DNA binding, histone methylation, histone acetylation, and chromatin remodeling), in a select number of complexes domains belonging to up to six functional groups were observed (all the above mentioned together with domains involved in DNA methylation and histone binding). These were all complexes involved in chromatin remodeling. For example, 'MBD2 or MBD3/NuRD nucleosome remodeling and deacetylase complex'. Notably, in chromatin complexes histone acetylation domains are found more often than histone methylation domains (unlike in the case when individual proteins are analyzed), this might, however, be biased by the current list of known chromatin complexes and their variability. Taken together chromatin protein complexes expand the multivalency of chromatin protein interactions and expand the functionality of the complexes.

As a final step we looked for chromatin proteins that had the most number of domain types (Pfam models) that belong to different functional groups/sub-groups and thus may engage in interactions of high multivalency (see **Supplementary Figure SF8_2A**). According to this analysis chromatin proteins could harbor in their domain composition domains from up to four functional groups/sub-groups. For example, histone-lysine N-methyltransferase ASH1L is involved in reading and writing of histone methylation, reading of histone acetylation and histone binding (Figure 8E). Proteins could harbor up to nine individual domains (where several domains belong to identical functional categories). One of such examples is Histone-lysine N-methyltransferase NSD2 (Figure 8E). This protein combines domains that likely engage in methylated histone binding (PWWP, PHD-1st_NSD, NDS_PHD, PHDvar_NSD, PHD, C5HCH), histone methylation (SET), histone binding (AWS) and may be DNA binding (HMG_box).
